# PlastidHub: an integrated analysis platform for plastid phylogenomics and comparative genomics

**DOI:** 10.1101/2025.03.19.644123

**Authors:** Na-Na Zhang, Gregory W. Stull, Xue-Jie Zhang, Shou-Jin Fan, Ting-Shuang Yi, Xiao-Jian Qu

**Affiliations:** College of Life Sciences, Shandong Normal University, Ji’nan 250358, China; Dongying Institute, Shandong Normal University, Dongying 257092, China; Department of Botany, National Museum of Natural History, Smithsonian Institution, Washington, DC 20013, USA; Germplasm Bank of Wild Species, Kunming Institute of Botany, Chinese Academy of Sciences, Kunming 650201, China

**Keywords:** Annotation, Comparative genomics, Plastid phylogenomics, Sequence processing, Visualization

## Abstract

The plastome (plastid genome) represents an indispensable molecular data source for studying plant phylogeny and evolution. Although plastome size is much smaller than that of nuclear genome, accurate and efficient annotation and of use plastome sequences are not simple tasks. Therefore, a streamlined phylogenomic pipeline from plastome annotation to phylogenetic reconstruction and comparative genomics would greatly facilitate research using this important organellar genome. Here we develop PlastidHub, a new web application that primarily uses novel tools to analyze plastome sequences. In comparison with existing tools, new functionalities in PlastidHub include: (1) standardization of quadripartite structure; (2) improvement of annotation flexibility and consistency; (3) quantitative assessment of annotation completeness; (4) diverse extraction modes for canonical and specialized sequences; (5) intelligent screening of molecular markers for biodiversity studies; (6) visual comparison of structural variations and annotation completeness at the gene-level. PlastidHub has the following features: cloud-based web applications that do not require users to install, update, or maintain tools; detailed help documents including user guides, test examples, a static pop-up prompt box, and dynamic pop-up warning prompts when entering unreasonable parameter values; batch processing functionalities for all tools; intermediate result documents for secondary use; and easy-to-operate task flows between file upload and download. Interrelated task-based user interface design is the most important feature of PlastidHub. Give that PlastidHub is easy to use without specialized computational skills or resources, this new platform should become widely used among botanists and evolutionary biologists, improving and expediting research employing the plastome. PlastidHub is available at https://www.plastidhub.cn.

## 1. Introduction

At present, analytical tools are divided into two categories: online web applications and local command-line scripts. The former are often more convenient, but their functionality is generally more limited, requiring users to visit multiple websites to fully process and analyze -omic data. The latter rely on the operation of scripts/codes, which can increase flexibility but requires users to have computation skills given complexities involved in installation and operation. Therefore, efforts to combine the benefits of both, by providing practical, integrative, and flexible tools through “one-stop” online platforms, are important for furthering genomics research. This is particularly true for plastome (plastid genomic) research, for which few *integrative* analytical platforms are available.

Due to its relatively small genome size, conservative gene content, and moderate evolutionary rate, the plastome is an indispensable molecular data source that has been widely applied in various fields including phylogenomics, evolutionary biology, comparative genomics, population genetics, phylogeography, and genetic engineering (Daniell et al., 2016; Jansen and Ruhlman, 2012; Jin and Daniell, 2015; Li et al., 2019; Straub et al., 2012; Stull et al., 2021; Wicke et al., 2011). There are many specialized tools commonly used for organelle or plastome analyses. Those for assembly include SPAdes (Bankevich et al., 2012), IOGA (Bakker et al., 2016), ORG.Asm (Coissac et al., 2016), NOVOplasty (Dierckxsens et al., 2017), Organelle_PBA (Soorni et al., 2017), chloroExtractor (Ankenbrand et al., 2018), GetOrganelle (Jin et al., 2020), and ptGAUL (Zhou et al., 2023). Those for annotation include DOGMA (Wyman et al., 2004), Plann (Huang and Cronk, 2015), Verdant (McKain et al., 2017), GeSeq (Tillich et al., 2017), AGORA (Jung et al., 2018), CpGAVAS2 (Shi et al., 2019), and PGA (Qu et al., 2019). Those for visualization include OGDRAW (Greiner et al., 2019), CGView (Stothard et al., 2019), and CPGView (Liu et al., 2023). Those for phylogenetics include HomBlocks (Bi et al., 2018), ORPA (Bi et al., 2024), and OGU (Wu et al., 2024). Finally, those for comparative genomics include VISTA (Frazer et al., 2004), progressiveMAUVE (Darling et al., 2010), IRscope (Amiryousefi et al., 2018), IRplus (Díez Menéndez et al., 2023), and CPJSdraw (Li et al., 2023). Despite the existence of multiple tools for analyzing plastomes, there are still various technical issues regarding plastome sequence processing and/or utilization, including standardization of genome structure, evaluation of annotation quality, accurate extraction of target sequences, streamlined pre- and/or post-alignment sequence processing, automatic screening of molecular markers, and visual comparison of genomes at the gene-level. These issues are described in detail below.

First, accurate identification of the quadripartite structure of plastomes remains a challenge. The plastomes of most photosynthetic seed plants are highly conserved, with a quadripartite structure comprising large (LSC) and small (SSC) single-copy regions and two identical inverted repeat (IR) regions (Jansen and Ruhlman, 2012; Wicke et al., 2011). In some cases, two IR copies are not completely identical, due to the imperfections (mismatches) or heterogeneities introduced into genome sequences (Díez Menéndez et al., 2023; Turudić et al., 2022), suggesting there is still room for improvement in IR identification (Turudić et al., 2022). The standardization of quadripartite structure cannot be separated from identifying the IR regions (Turudić et al., 2022). At present, the main plastome annotation tools, web application GeSeq (Tillich et al., 2017) and command-line application PGA (Qu et al., 2019), consider the identification of IR copies, but cannot adjust the quadripartite structure. IRscope (Amiryousefi et al., 2018) and CPJSdraw (Li et al., 2023) are commonly used tools only for identifying IR copies, with the latter one considering accurate junction of LSC-IRa, but these two tools cannot output post-adjusted plastome sequences. In addition, NOVOWrap (Wu et al., 2021) and Plastaumatic (Chen et al., 2022) directly claim to have the functions of adjusting quadripartite structure from non-standardized form to standardized form. However, the “validate.py” Python script in NOVOWrap depends on the reference sequence to finish the adjustment process, and the incorrect SSC orientation of the target species will not be reversed when selecting distantly related or unsuitable reference species. The “standardize_cpDNA.sh” bash script in Plastaumatic cannot be run without a Linux operating system.

Second, accurate annotation of whole plastomes remains a challenging task, despite the availability of multiple plastome annotation tools, including web tools such as GeSeq (Tillich et al., 2017), CPGAVAS2 (Shi et al., 2019), and Chloe (https://chloe.plastid.org/annotate.html), and the command-line tool PGA (Qu et al., 2019). The principles of available plastome annotation tools were reviewed and compared in Qu et al. (2023). The specific genes that need special attention are pseudogenes, redundant genes, and the trans-splicing gene *rps12*. Pseudogenes are non-functional gene fragments that are derived from and similar to functional genes (Goodhead and Darby, 2015). Any gene could become a pseudogene due to mutations resulting in premature stop codons or frameshifts (Xiao et al., 2016). The existence of pseudogene complicates genome annotation (Qu et al., 2023). Annotation of these evolutionary remnants in the intergenic regions is very important for further exploration of plastomes, including but not limited to the generation of pseudogenes, the origin of new genes, and the occurrence of rearranged plastomes. However, there is almost no tool that has the function of annotating evolutionary remnants. In addition, trans-splicing refers to the linking of two or more exons from two different pre-mRNAs. In contrast to canonical cis-splicing, two or more exons are from different pre-mRNAs, but also may be from the same gene. As a trans-splicing gene containing three or two exons of two pre-mRNAs, the *rps12* gene is often annotated incorrectly or incompletely, because the three or two exons are usually difficult to link manually (Qu et al., 2023; Shi et al., 2019; Tillich et al., 2017). Presently, there are no tools that can reliably annotate *rps12* with accuracy, and there is no evaluation report on the hit rate and link accuracy of *rps12*.

Third, tools for quantitatively evaluating plastome annotation quality are still lacking. There are some efficient methods for assessing the completeness of nuclear genome assembly/annotation, such as BUSCO (Waterhouse et al., 2018) and CEGMA (Parra et al., 2007). In our previous study, we proposed three methods for quantitatively evaluating plastome annotations, i.e., gene number comparison, gene length difference comparison, and gene sequence similarity comparison (Qu et al., 2023). However, there is a lack of tools to implement these methods. In addition, there is still a lack of one-to-one methods for visual comparison of genes in evaluating the annotation quality of plastomes.

Fourth, the accurate extraction of coding and noncoding sequences from plastomes is a complicated task. The genic composition of plastomes is complex, including PCG genes and RNA (tRNA and rRNA) genes, intron-containing genes and genes without introns, cis-splicing genes and trans-splicing genes, non-overlapped and non-nested genes and overlapped and nested genes, single-copy genes and genes with two or three copies, as well as genes that span the start-end position and the start-end position within the noncoding (intergenic/intronic) regions (Qu et al., 2023; Raubeson and Jansen, 2005). Currently, there are no specific tools with multiple extraction modes to meet the diversified needs of users.

Fifth, the tools that can perform pre- and/or post-alignment processing are not integrated, and it is difficult to search customed tools that meet these scattered and personalized needs. Although most land plants have a conserved plastome structure, many lineages nevertheless have highly rearranged plastome structures, resulting in disordered gene order (Knox, 2014; Mower and Vickrey, 2018; Qu et al., 2022). In this case, determining how to use intergenic regions can be a major challenge (Shaw et al., 2007). In addition, for plant taxonomists with little or no programming experience, the following tasks can be prohibitively difficult: checking the codon status of PCGs, listing the missing taxa for each sequence, extracting each codon site in each PCG, extracting sub-alignments for each alignment matrix, generating alignment files formatted for PartitionFinder, and automatically screening markers by evaluating the degree of sequence variation in all alignment matrices. Therefore, the development of streamlined methods for processing pre- and/or post-alignment sequences is a critical need.

Sixth, tools for micro-synteny comparison of plastomes are lacking. Considering the limited number of genes in the plastome (Raubeson and Jansen, 2005), micro- synteny comparison of each gene is realistic and effective. It can not only display structural variations such as inversion, translocation, and duplication (Mower and Vickrey, 2018; Palmer, 1991), but also show the degree of annotation completeness such as gene loss/gain (Qu et al., 2023). Accordingly, pairwise visual comparison of each plastid gene can also be applied to evaluate the completeness of plastome annotations at the gene-level. However, at present, there is no specific tool for executing micro-synteny comparison of plastomes.

To address these significant gaps in analytical tools for plastomes, we developed online platform (PlastidHub) employing an interrelated task-based user interface design for comprehensive analysis of plastome sequences. All tools in PlastidHub possess batch processing capability, which is indispensable given the scale of datasets being routinely analyzed by botanists and genomic researchers.

## 2. Materials and Methods

### 2.1. Implementation

PlastidHub is a web application developed with Java, TypeScript, and Perl, which is convenient for users who prefer browser-driven applications or those who are not familiar with UNIX command-line tools. The front-end web interface was developed with vue3 (https://cn.vuejs.org/) and Element-Plus (https://element-plus.org/). PlastidHub has been tested on recently released web browsers, such as Google Chrome, Microsoft Edge, Mozilla Firefox, Opera, Apple Safari, 360 Speed Browser X, 360 Secure Browser, QQ Browser, and Huawei Browser. The back-end web framework uses SpringBoot v3.3.2 (https://docs.spring.io/spring-boot/index.html) with RabbitMQ (https://www.rabbitmq.com/) as message queue and Redis (https://redis.io/) storing result status, which can manage data and parameters submitted by users, and then relay the analysis results (generated files) to the front-end interface for users to manage (select all, clear selection, download, and delete). The back-end tools of PlastidHub were developed with Perl v5.26.1 programming language. All Perl scripts within PlastidHub can be executed in the command-line terminal of any computer systems (Windows, Linux, and Mac).

### 2.2. Analysis workflow

PlastidHub is an integrated plastome analysis platform including 10 tool kits: annotation, assessment, submission, visualization, extraction, pre-alignment, (post)- alignment, phylogeny, barcoding, genomics, each of which contains one or a few tools for performing specific tasks. The analytical workflows of PlastidHub are not fixed and can be customized by users. For examples, “annotation → assessment” is an analysis workflow; “annotation → assessment → submission” is another workflow; “annotation → assessment → visualization” is a third analysis workflow; “annotation → assessment → extraction → pre-alignment → (post)-alignment → phylogeny” is a fourth analysis workflow; “annotation → assessment → extraction → pre-alignment → (post)- alignment → barcoding” is a fifth analysis workflow; and “annotation → assessment → plastomics” is a sixth analysis workflow. Certainly, users can choose any tools without starting from early steps, such as annotation and/or assessment.

### 2.3. Supported file formats

PlastidHub supports GenBank-formatted (.gb/.gbf/.gbk), FASTA-formatted (.fasta/.fa/.fas/.fsa), and TEXT-formatted (.txt) input files, as indicated in the File Upload Panel of each tool. The format of output files is determined by the user, with possible types being GenBank, FASTA, BED (.bed), TABLE (.tbl), TAB-delimited (.txt), SVG (.svg), PNG (.png), TREE (.tre), and CONFIG (.conf).

### 2.4. File number and size limitation

The file number and size allowed by PlastidHub are shown in the static pop-up prompt box of each tool.

### 2.5. Special characters in filenames

Some functionalities in PlastidHub rely on third-party tools, so the use of certain special characters (∼!@#$%^&*()+=[]}{;:’"<>,/?|) is not recommended. Also, it is not recommended to use non-English characters in filenames. In particular, we recommend to using an underscore (_) or dot (.) instead of spaces () or dash line (-).

### 2.6. Parameter threshold range

The parameters that can be modified by users in PlastidHub have been set to a fixed range of thresholds, which prevents inappropriate parameter values from being selected.

### 2.7. Resource consumption

The run time of each tool is very short, and depends on the number of files. In theory, the processing time for each file should be less than 5 seconds. Because all tools do not need to handle large files, there is no need to consider the consumed computational resources.

### 2.8. Test data and tool evaluation

To evaluate the performance of some newly developed tools in PlastidHub, we downloaded 15,206 eukaryotic plastome sequences from NCBI RefSeq database (https://ftp.ncbi.nlm.nih.gov/refseq/release/plastid/, accessed 2024.5.16). First, we count the proportion of non-standardized quadripartite structure in 15,206 eukaryotic plastome sequences using the Quadripartition tool. Second, using 15,206 eukaryotic and 13,731 angiosperm plastome sequences, we evaluate the hit rate and link accuracy of *rps12* with the plastome annotation tool PGA v2.0. To evaluate plastome annotation consistency of PGA v2.0, the number of annotated genes in GenBank and PGA annotations were compared for the 13,731 angiosperm plastid sequences. Third, using *A. trichopoda* as reference, we applied the tools of “Assess Gene Number” and “Assess Gene Length” to evaluate the annotation quality of the target species *Rosa roxburghii*. Fourth, to evaluate the accuracy of the plastome Extraction tool, the number of extracted genes and PGA-annotated genes in the 13,731 angiosperm plastid sequences was compared. Fifth, to evaluate the effectiveness of the Barcoding tool in automatically screening molecular markers for plastomes, plastomes from Amaranthaceae species including *Amaranthus caudatus*, *A. tricolor*, *Atriplex centralasiatica*, *Chenopodium album*, *C. quinoa*, *Salicornia bigelovii*, *S. europaea*, *Spinacia oleracea*, *Suaeda glauca*, and *S. salsa* were used. Sixth, to evaluate the functionality (showing micro-synteny structural variations and evaluating annotation completeness) of the Gene Homology tool, we show plastome comparisons of *Amborella trichopoda* against other angiosperms (*Rosa roxburghii* and *Salicornia bigelovii*) and a gymnosperm species (*Calocedrus decurrens*), as well as a comparison of the gymnosperm species *Calocedrus decurrens* and *Calocedrus rupestris*.

## 3. Results and Discussion

### 3.1. Interrelated user interface design of PlastidHub

To create an easy-to-use web-based tool kit for researchers engaged in phylogenomic and comparative genomic analyses using plastome data, we generated numerous task-based interfaces where the users can easily choose a tool to accomplish their analysis goals (Fig. 1). The home page of PlastidHub contains three interfaces, i.e., “P-Utils” (tool interface), “Document” (help interface), and “My Files” (download interface).

**Fig. 1.**
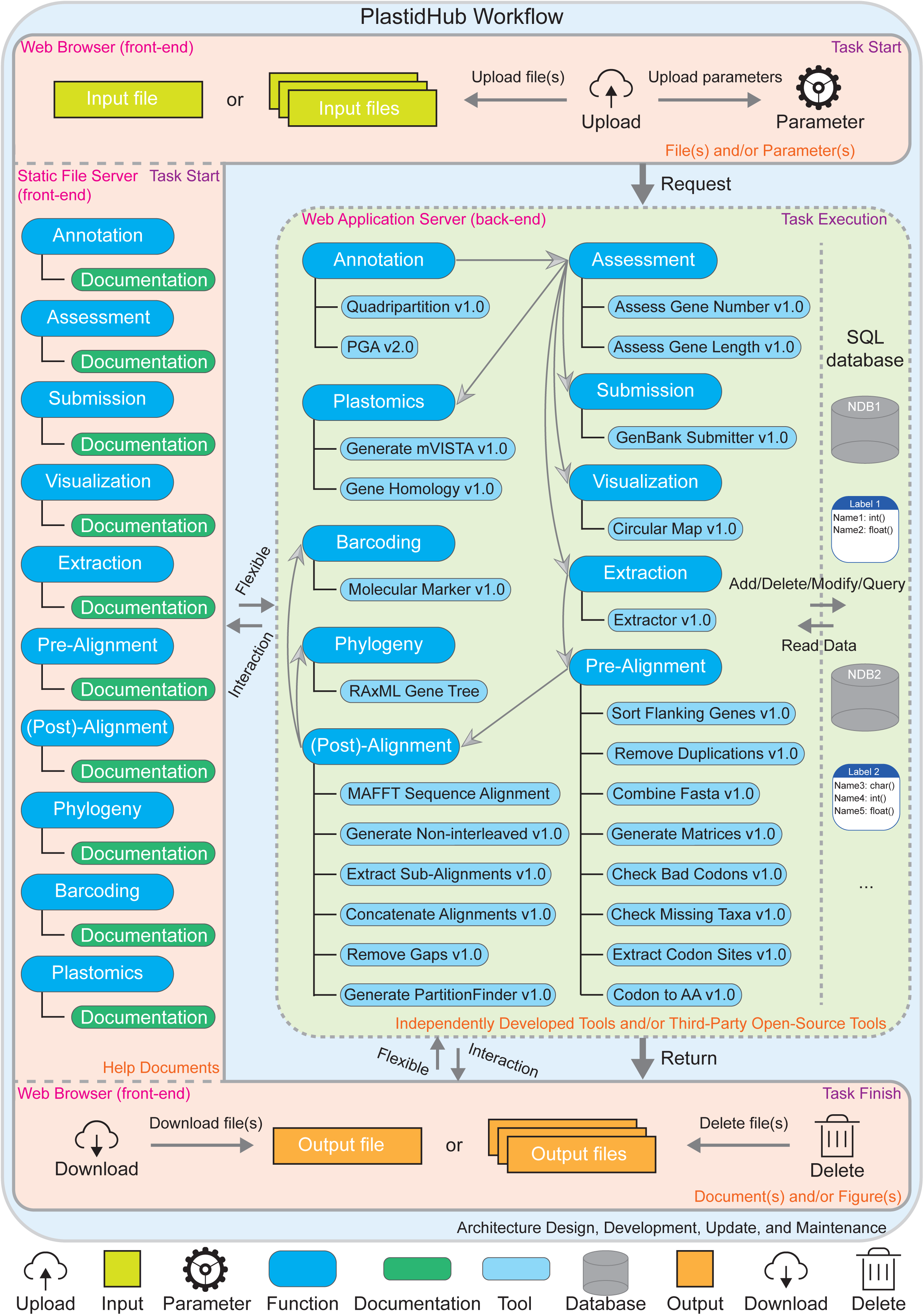
Overview of interrelated task-based user interface design in PlastidHub. Task Start includes front-end Web Browser for users to upload input Files and Parameters, and front-end Static File Server for users to look up Help Documents. Task Execution includes back-end Web Application Server for users to run tools. Task Finish includes front-end Web Browser for users to download and/or delete output Files (Documents and Figures). The C-word pink background indicates frond-end interface, and the light-green background indicates back-end interface. The Request and Return between front-end interface and back-end interface are shown. Left Flexible Interaction indicates the switch between tool interface and help interface, and bottom Flexible Interaction indicates the switch between download interface and tool inferface. Within the tool interface, data can be stored and read interactively between the tools and the database.

For the “P-Utils” tool interface, the left panel shows the links to task-running interfaces of 25 tools within 10 tool kits. The right panel of the “P-Utils” interface lists all 25 tools in PlastidHub, and users can directly access each specific tool. Each task- running tool has a brief introduction to help users make the right choice. For the separate task-running interface of each tool in the “P-Utils” interface, there are interactive options for uploading input files (required), selecting parameter values (optional), submitting (required) or resetting (optional) the task, and linking to the help document for reference. To avoiding submission errors, the maximum number of input files is limited, the formats (and allowed suffix) of input files are specified, and the parameter values are specified as an allowed range. In addition, if the input file(s) is/are not selected, the submit button will not become operable and a warning prompt such as “Please upload reference/target file first” will be displayed for a short duration (3s).

For the “Document” help interface, the left panel shows the links to help documents of 10 tool kits. The right panel of “Document” interface lists “Get Started”, “User Guide”, “Browser Compatibility”, “Tech Stack”, and “Content”. For the “Content” section, a hyperlink-based help document for each of 25 tools is listed. Within the help document of each tool, paragraph navigation links such as “General Introduction”, “Features”, “Command-line Tool”, and “Example” are set up to help users quickly locate the target paragraph.

An important customization feature of PlastidHub is the “My Files” download interface, allowing users to check the running status of the submitted tasks and download or delete the completed tasks at any time. Intermediate result documents are also provided to assist users in secondary processing. The validity period of the result files stored in the server database is one week. If the download button is clicked, a prompt such as “The result of output files will be stored for a week since task finished” will be displayed. Similarly, if the delete button is clicked, a warning prompt such as “Comfirm that you are deleting the result(s)” will be displayed. Each tool kit has a corresponding download interface, and the tool interface is located below the corresponding download interface, making it convenient for users to access directly. It indicates that the tool interface can be freely switched not only with the help interface, but also with the download interface. This design enhances the flexibility of running tasks in PlastidHub.

The middle panel of the home page contains user access and usage statistics, including “Task Handled”, “Page Views”, and “Unique Visitors”, providing real-time feedback on user visits and completed tasks. The bottom panel of the home page contains a “Links” section, which includes “GitHub” and “Change Log”, and a “Community” section, which includes “Feedback”, “Get Involved”, and “Contact”, as well as a “Policy” section, which includes “Terms of Use” and “Privacy Policy”.

### 3.2. Functionalities of 25 tools within 10 designed tool kits

#### 3.2.1. Annotation

This tool kit contains two tools that can complete standardization of quadripartite structure and plastome annotation.

##### 3.2.1.1. Quadripartition

This tool, which can operate on batch submissions, adjusts the quadripartite structure of plastomes from a non-standardized form to a standardized form (LSC-IRb-SSC-IRa). Because some plastome sequences generated in the laboratory or downloaded from NCBI are in non-standardized form or in opposite direction, it is necessary to adjust the structures and orientations of these plastome sequences for downstream analyses.

##### 3.2.1.2. Annotation

Plastome annotation leverages our previously developed tool PGA (Plastid Genome Annotator) (Qu et al., 2019). The main principle of plastome annotation remains unchanged, using the reference plastome as the query and unannotated target plastomes as the subject to locate genes, which we refer to as the reverse query-subject BLAST search approach. However, we have updated PGA v1.0 to a newer version, PGA v2.0, adding some new functions to annotate evolutionary remnants, and link *rps12*.

#### 3.2.2. Assessment

This tool kit contains two tools that can assess plastome annotation quality, serving as a benchmark for achieving high-quality genome annotation.

##### 3.2.2.1. Assess Gene Number

This tool assesses annotation completeness of plastomes as number of hitting, missing, and redundant genes. Specifically, annotation completeness of plastomes is assessed by comparing numbers and names between genes in the target plastome and corresponding genes in the reference plastome.

##### 3.2.2.2. Assess Gene Length

This tool assesses annotation accuracy of genes by comparing gene length difference. Specifically, annotation accuracy of plastomes can be assessed by comparing length differences between genes in the target plastome and their corresponding genes in the reference plastome.

#### 3.2.3. Submission

This tool kit contains a single tool that can convert batch submissions of multiple GenBank formatted flatfiles (.gb/.gbf/.gbk) to the 5-column feature table files (.tbl) and sequence files in FASTA format (.fsa/.fasta/.fas/.fa) with required information for submitting to GenBank.

#### 3.2.4. Visualization

This tool kit contains a single tool that can accept batch submissions and generate circular plastome maps with genes, GC content, quadripartite structure, and dispersed repeats.

#### 3.2.5. Extraction

This tool kit contains a single tool that can extract coding and noncoding sequences from batch submissions of multiple GenBank flatfiles of plastomes based on different extraction patterns.

#### 3.2.6. Pre-Alignment

This tool kit contains eight tools that can complete pre-alignment processing of sequence matrices.

##### 3.2.6.1. Sort Flanking Genes

This tool sorts flanking gene names of batch submissions and reverse complements their corresponding sequences for the intergenic regions in plastomes. Because some plastomes have obvious structural rearrangements such as inversions and translocations, it is necessary to adjust their sequences to the same direction before aligning intergenic regions for phylogenetic reconstruction.

##### 3.2.6.2. Remove Duplications

This tool removes duplicated sequences from FASTA files. It is useful for processing plastomes with quadripartite structures, because the genes within the two IR copies are identical, while there are few genes that can be duplicated in the LSC or SSCregions. For the genes with duplicated gene names but non-duplicated sequences, users are able to decide how to handle individual cases.

##### 3.2.6.3. Combine Fasta

This tool combines multiple FASTA files containing one or more species.

##### 3.2.6.4. Generate Matrices

This tool generates sequence matrices including multiple species for each coding and noncoding region of the plastome. The generated sequence matrices can be subsequently aligned.

##### 3.2.6.5. Check Bad Codons

This tool checks for bad codons in batch submissions of sequence matrices of PCGs.

##### 3.2.6.6. Check Missing Taxa

This tool checks for missing taxa from batch submissions of multiple-sequence matrices.

##### 3.2.6.7. Extract Codon Sites

This tool extracts codon sites from batch submissions of PCGs.

##### 3.2.6.8. Codon-to-AA

This tool translates codons to amino acids for batch submissions of PCGs. The generated amino acids sequence matrices can be aligned by alignment tools.

#### 3.2.7. (Post)-Alignment

This tool kit currently contains six tools that conduct alignment and post-alignment processing of sequence matrices.

##### 3.2.7.1. Multiple Sequence Alignment

This tool performs multiple sequence alignment on batch submissions of matrices using the mafft alignment tool.

##### 3.2.7.2. Generate Non-interleaved

This tool generates non-interleaved FASTA files from batch submissions of interleaved FASTA files.

##### 3.2.7.3. Extract Sub-alignments

This tool extracts sub-alignments from batch submissions of FASTA sequence alignments given a list of species names. Simultaneously, this tool removes gap sites from the sub-alignments.

##### 3.2.7.4. Concatenate Alignments

This tool concatenates supplied FASTA alignments.

##### 3.2.7.5. Remove Gaps

This tool removes gap sites from batch submissions of alignments.

##### 3.2.7.6. Generate PartitionFinder

This tool will generate files formatted for PartitionFinder from input FASTA or Phylip alignments.

#### 3.2.8. Phylogeny

This tool kit contains a single tool that performs maximum likelihood phylogenetic inference on batch submissions of aligned matrices.

#### 3.2.9. Barcoding

This tool calculates and extracts variable, invariable, and gap sites from batch submissions of alignments.

#### 3.2.10. Genomics

This tool kit contains two tools that perform comparative analysis of plastome sequences.

##### 3.2.10.1. Generate mVISTA

This tool generates input files for mVISTA alignment from batch submissions of GenBank flatfiles.

##### 3.2.10.2. Gene Homology

This tool generates circular views of two plastomes with homologous genes being connected and non-shared genes being marked. First, the homologous genes in the target plastome and their corresponding genes in the reference plastome are connected. Thus, the collinear state of these two plastomes can be judged by the connection status of homologous genes. Second, the genes that are not shared in the reference or target plastome are marked. Therefore, annotation completeness of plastomes can be assessed by comparing homologous genes and labeled non-shared genes.

### 3.3. Main function 1: High proportion of non-standardized quadripartite structure of plastomes in NCBI RefSeq database identified by the Quadripartition tool

The quadripartition tool in PlastidHub (Fig. 2A) is newly developed based on the optimization of the IR-copy-identification algorithm in our first release of PGA v1.0 (Qu et al., 2019). Flexibility is our main consideration for adjusting quadripartition structure. Therefore, we allow the IRb and IRa to be set as completely (100%) or almost (99%) identical, the minimum allowed IR length to be changed manually (e.g., > 1000 bp or > 100 bp), the orientation of the whole plastome sequences or the SSC sequences to be reverse complemented or not, and the standardized plastome sequences as well as coordinates and sequences of LSC, IRb, SSC, and IRa to be outputed.

**Fig. 2.**
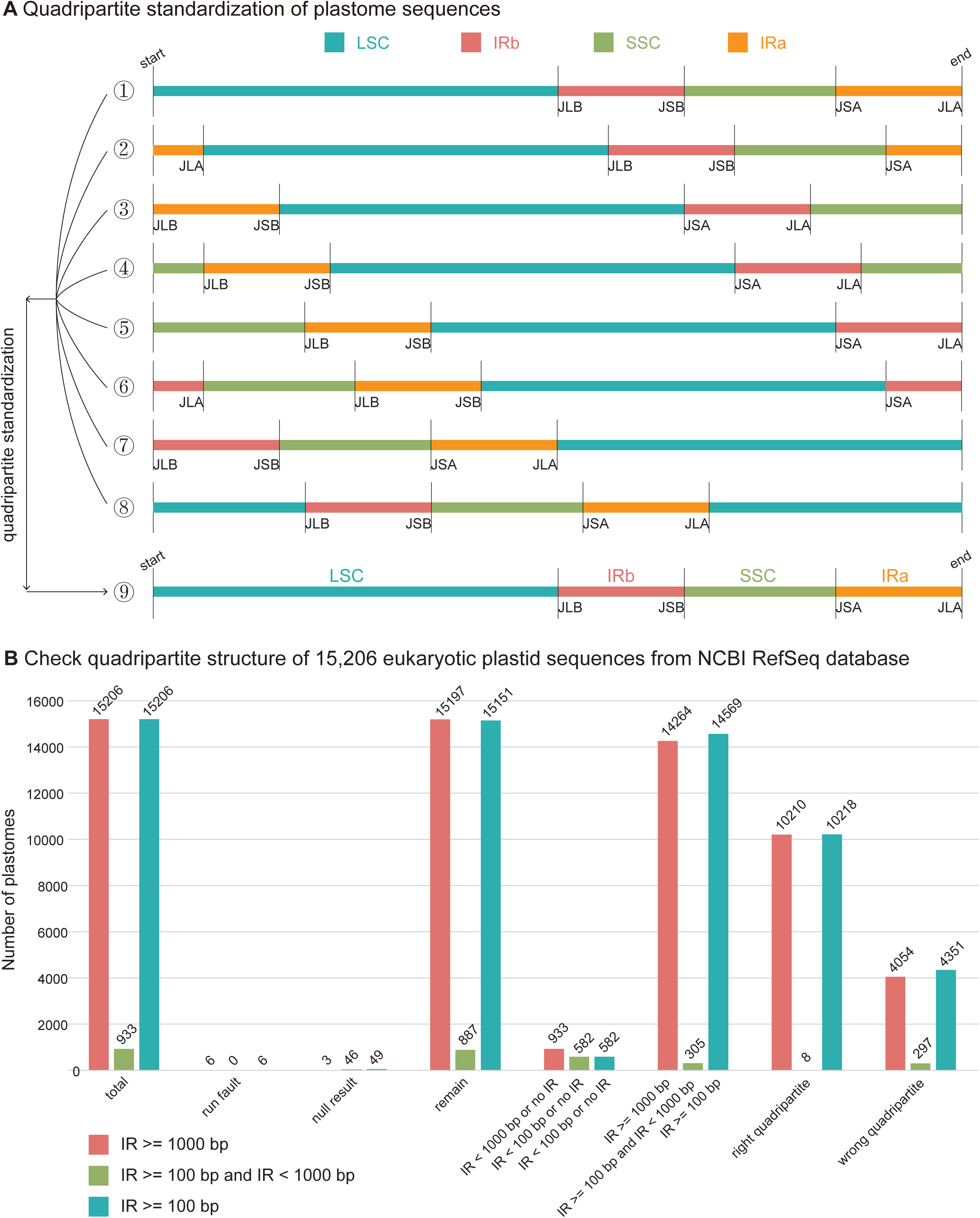
Standardization of quadripartite structure for plastome sequences. (A) Quadripartite standardization of plastome sequences. ①—⑧ indicate unknown quadripartite structure including non-standardized and possibly standardized forms. ⑨ indicate standardized quadripartite structure. (B) Check quadripartite structure of 15,206 eukaryotic plastid sequences from NCBI RefSeq database. Three scenarios are showed: IR >= 1000 bp, IR >= 100 bp and IR < 1000 bp, and IR >= 100 bp.

The proportion of non-standardized quadripartite structures were counted in 15,206 eukaryotic plastome sequences (Fig. 2B). If we set IR >= 1000 bp, except 9 abnormal sequences with run fault and null result, there are 933 sequences with IR < 1000 bp or no IR and 14,264 sequences (93.9% hit rate) with IR >= 1000 bp. For the 14,264 sequences, there are 10,210 sequences (71.6% accuracy rate) with correct quadripartite structure and 4054 sequences with incorrect quadripartite structures. If we set IR >= 100 bp and IR < 1000 bp, except 46 abnormal sequences with null result, there are 582 sequences with IR < 100 bp or no IR and 305 sequences (34.4% hit rate) with IR >= 100 bp and IR < 1000 bp. For the 305 sequences, there are 8 sequences (2.6% accuracy rate) with the correct quadripartite structure and 297 sequences with the incorrect quadripartite structure. In total, if we set IR >= 100 bp, except 54 abnormal sequences with run fault and null result, there are 582 sequences with IR < 100 bp or no IR and 14,569 sequences (96.2% hit rate) with IR >= 100 bp. For the 14,569 sequences, there are 10,218 sequences (70.1% accuracy rate) with the correct quadripartite structure and 4351 sequences with the incorrect quadripartite structure. These results indicate that the proportion of plastomes with non-standardized quadripartite structures in NCBI RefSeq database is very high, and the incorrect start-end position of circular plastomes is primarily caused by the offset of LSC-IRa junction (Turudić et al., 2022).

### 3.4. Main function 2: Flexibility and consistency of the Annotation tool PGA v2.0

We updated PGA v1.0 to PGA v2.0, providing flexible options to annotate or unannotate evolutionary remnants, namely pseudogenes and/or redundant genes (Fig. 3A). In some cases, it is difficult to distinguish pseudogenes from redundant genes, especially in highly divergent plastomes (Qu et al., 2023). Indeed, this is the case for PGA v2.0, which cannot distinguish redundant false-positive gene fragments from pseudogenes. Nonetheless, this function is still effective for searching pseudogene residues, which are important remnants for tracing the evolutionary history of these fast-evolving genes, especially in heterotrophic plants with highly degraded plastomes (Goodhead and Darby, 2015; Hsu et al., 2016; Xiao et al., 2016). If users want to annotate plastome sequences for regular phylogenetic analysis, we suggest avoiding the annotation of redundant genes and/or pseudogenes, avoiding potential alignment errors that could impact phylogenetic inference; if users want to annotate plastome sequences for comparative genomics including RNA-editing events, we suggest annotating redundant genes and/or pseudogenes, providing a chance to explore the degradation extent of functioanl genes due to the general lack of selective constraints.

**Fig. 3.**
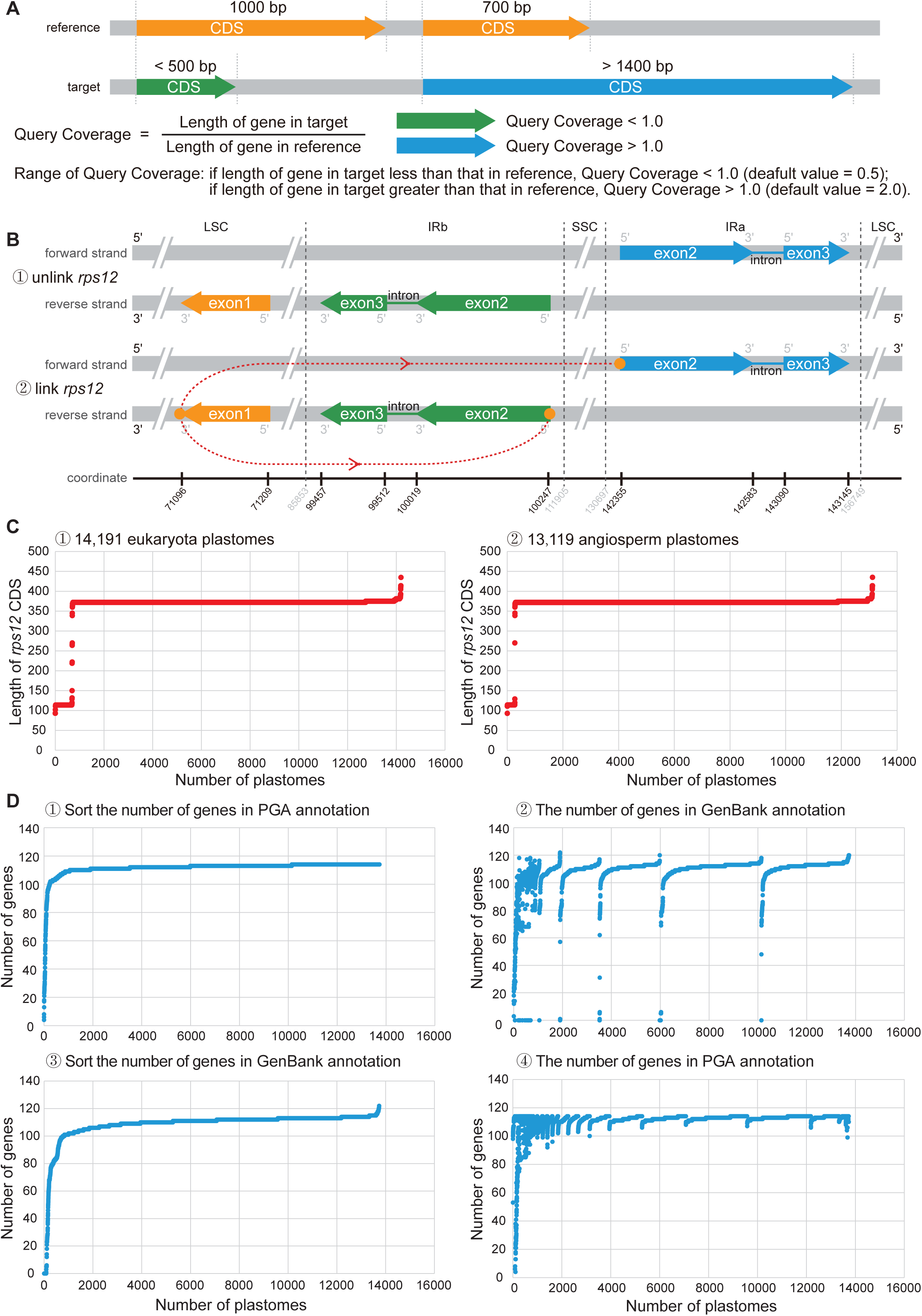
Improvement of annotation personalization for the plastome. (A) Annotation redundancy: annotate or unannotate evolutionary remains. The evolutionary remains (e.g., pseudogenes) can be unannotated by changing dafault parameter value (Y: annotate pseudogenes) in PGA v2.0. (B) Annotation link: ① unlink *rps12* or ② link *rps12*. The trans-splicing gene *rps12* can be unlinked by changing default parameter value (Y: link *rps12*) in PGA v2.0. (C) Evaluation of the annotation status of *rps12* by re-annotating 15,206 eukaryotic plastid sequences from NCBI RefSeq database with PGA v2.0. (D) Comparison of the number of annotated genes in GenBank and PGA using 13,731 angiosperm plastid sequences in NCBI RefSeq database.

We also provided a flexible parameter in PGA v2.0 to link or unlink *rps12*, which can meet the needs of different users when annotating plastomes using PGA. The *rps12* gene is a trans-splicing gene, containing three exons of two pre-mRNAs, including exon 1 in LSC, and exon 2 and exon 3 in IR (Fig. 3B). The *rps12* gene is often annotated incorrectly or incompletely, because the three exons are usually difficult to link manually (Qu et al., 2023; Shi et al., 2019; Tillich et al., 2017). The 15,206 eukaryotic plastid sequences from NCBI RefSeq database were used to evaluate the annotation and link status of *rps12* (Fig. 3C). Using *Amborella trichopoda* (angiosperm) as reference, 15,206 eukaryotic plastid sequences were re-annotated using PGA v2.0 with default parameter values. After running, 15,200 sequences were successfully annotated, while 6 sequences from Chlorophyta were not successfully annotated. For 15,200 eukaryotic sequences, 14,191 sequences could annotate *rps12* (93.4% hit rate), and the length of the annotated *rps12* ranged from 93 to 435 bp, while 1009 sequences were not annotated with *rps12*. For 14,191 eukaryotic sequences, 13,494 sequences were successfully linked for *rps12* (95.1% link accuracy), and the length of the annotated *rps12* ranged from 339 to 435 bp, while 697 sequences were only annotated with one exon (93 to 270 bp). For 13,731 angiosperm sequences, 13,119 sequences could annotate *rps12* (95.5% hit rate), and the length of the annotated *rps12* also ranged from 93 to 435 bp, while 612 sequences were not annotated with *rps12*. For 13,119 angiosperm sequences, 12,845 sequences were successfully linked for *rps12* (97.9% link accuracy), and the length of the annotated *rps12* also ranged from 339 to 435 bp, while 274 sequences were only annotated with one exon (93 to 270 bp). These results indicate that even if we use angiosperm *A. trichopoda* as a reference to annotate angiosperm or eukaryotic plastome sequences, the hit rate and link accuracy of *rps12* using PGA are all high.

To further evaluate the PGA v2.0 tool in PlastidHub, the number of annotated genes in GenBank and PGA were compared using the 13,731 angiosperm plastid sequences from the NCBI RefSeq database (Fig. 3D). The genes extracted from these 13,731 angiosperm plastid sequences represented GenBank annotation. Using *A. trichopoda* (angiosperm) as reference, the 13,731 angiosperm plastid sequences were re-annotated using PGA v2.0 with default parameter values, and the number of the extracted genes in these 13,731 angiosperm plastid sequences represented PGA annotation. The number of genes extracted from the GenBank and PGA annotations were then compared. The order of accession numbers for the angiosperm plastid sequences from PGA and GenBank was the same between ① and ② in Fig. 3D, as well as between ③ and ④ in Fig. 3D. Only 224 plastid sequences had gene numbers fewer than 100 for the PGA annotation (Fig. 3D ①), while there were 759 plastid sequences with gene numbers fewer than 100 for the GenBank annotation (Fig. 3D③). By comparing ② with ① in Fig. 3D, we found that the plastid sequences with gene numbers under 100 were not concentrated in the fixed range for the GenBank annotation, indicating significant differences in annotation quality of plastid sequences in GenBank annotations. In contrast, by comparing ④ with ③ in Fig. 3D, we found that the plastid sequences with gene numbers under100 were concentrated in the fixed range for the PGA annotation, indicating high consistency in annotation quality of plastid sequences in the PGA annotations. Importantly, the plastid sequences annotated by PGA with fewer than 100 genes were a subset of plastid sequences annotated in GenBank with fewer than 100 genes, indicating that the limited number of annotated genes in both the PGA and GenBank annotations was likely due to plastid sequence features. The NCBI website states that the organelle genomes with curated annotation information from the RefSeq project can be used as the standard. Given that plastome sequencing has been ongoing for several decades, employing various approaches and standards for data generation, assembly, and annotation, it is not surprising that there is considerable variation in data quality.

### 3.5. Main function 3: Evaluating the annotation quality of plastomes by the Assessment tool

To evaluate the annotation quality of plastomes, we proposed two quantitative methods: assessing gene number and assessing gene length (Qu et al., 2023). Using *A. trichopoda* as the reference species, we applied these two methods to evaluate the annotation quality of the target species *Rosa roxburghii* (Fig. 4). For gene number assessment, the annotation completeness of genes can be judged by number of hitting (nHG), missing (nMG), and redundant (nRG) genes; percentage of hitting (pHG), missing (pMG), and redundant (pRG) genes; number of hitting (nHP, nHT, nHR), missing (nMP, nMT, nMR), and redundant (nRP, nRT, nRR) PCGs, tRNAs, and rRNAs; and name of hitting, missing, and redundant PCGs, tRNAs, and rRNAs (Fig. 4A). Since plastomes do not contain many genes (∼100), it would be reasonable for users to manually check the annotation status (hitting, missing, redundant) of each gene, and users can determine why genes are missing (due to true loss or missing annotation for a gene that is actually present) or redundant (redundant false-positive genes or pseudogenes or missing in reference). For gene length assessment, annotation accuracy of genes can be assessed by comparing length differences between genes in the target plastome and their corresponding genes in the reference plastome (Fig. 4B). Gene length differences can then be sorted by magnitude, and those with the largest differences (or those larger than a specified threshold) can be checked. Gene lengths of tRNAs and rRNAs are more conserved than those of PCGs across plant species, with rare exceptions, so the length differences of PCGs, and tRNAs and rRNAs are separately compared. Together, the tools of “assess gene number” and “assess gene length” in PlastidHub are very practical for users who want to produce accurate annotations of gene numbers and gene boundaries.

**Fig. 4.**
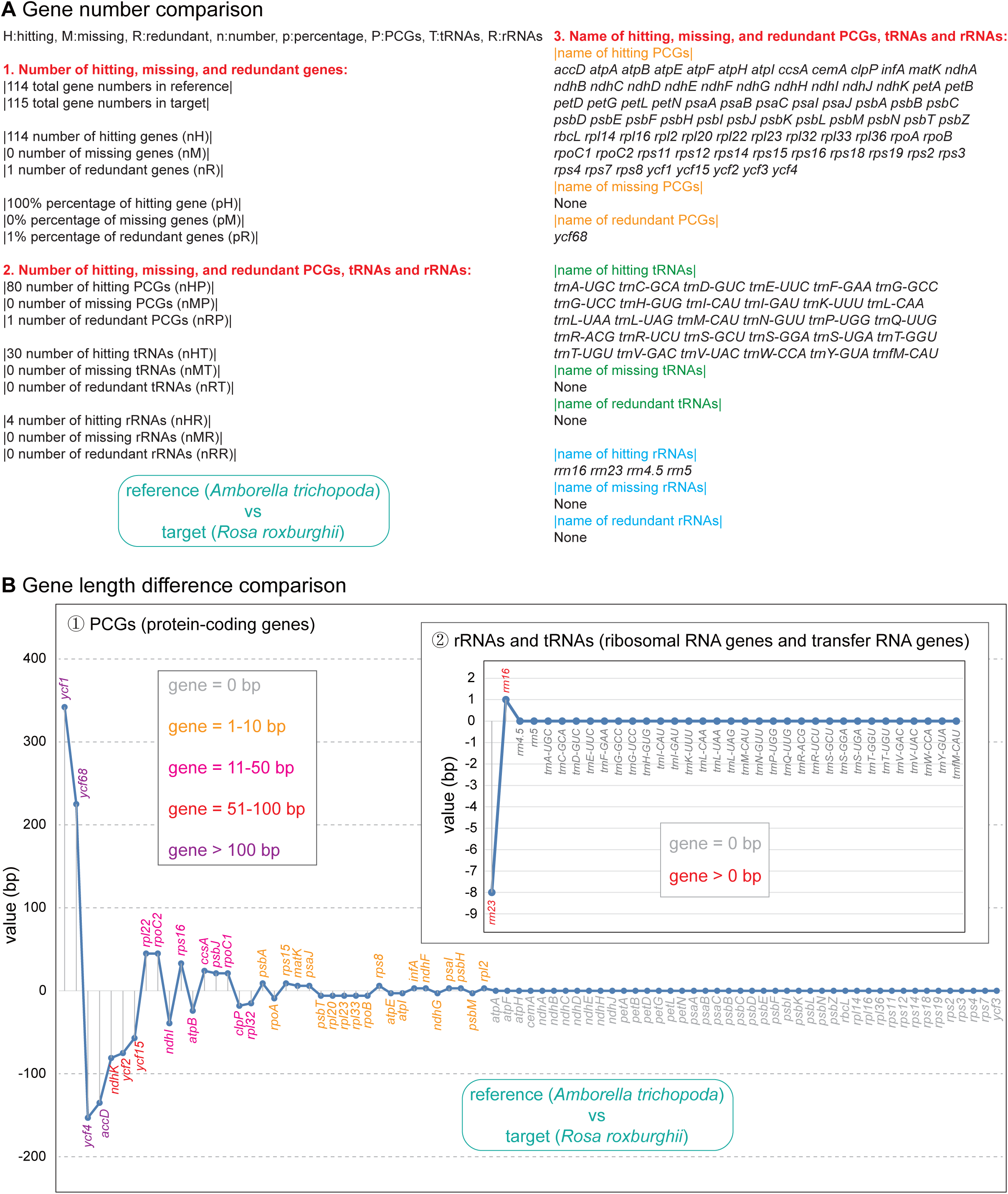
Standards for quantitatively evaluating the annotation quality of plastomes. (A) Gene number comparison between target and reference plastomes. Annotation completeness of plastomes is assessed by comparing numbers and names of hitting, missing, and redundant genes. (B) Gene length difference comparison between target and reference plastomes. Annotation accuracy of plastomes is assessed by comparing length differences between genes in the target plastome and their corresponding genes in the reference plastome. The length differences of ① PCGs and ② RNAs are separately compared. The gene length differences are sorted by magnitude, and those larger than a specified threshold (or those with the largest differences) are indicated as different color.

### 3.6. Main function 4: Accuracy of the plastome Extraction tool

Extraction of coding and noncoding sequences from multiple species is an important step before multiple sequence alignment and subsequent analysis of selection or phylogenetic reconstruction. Given the circular structure and diverse gene types of plastomes, accurate extraction of all target sequences is not trivial. Our newly developed extraction tool in PlastidHub considers all extraction patterns (Fig. 5), not only canonical scenarios such as genes with or without introns. This includes special scenarios such as nested genes in the plastome (Fig. 5A), trans-splicing genes in the plastome (Fig. 5B), genes that do not span the start-end position of circular plastome (Fig. 5C), and genes that do span the start-end position of circular plastome (Fig. 5D). In addition to PCGs with introns and sequences among PCGs, genes with introns and intergenic sequences are extracted. Furthermore, coding sequences with linked and unlinked CDSs and tRNAs are considered, as well as noncoding sequences. Finally, different extraction requirements can be implemented using the flexible set of parameters provided with this tool.

**Fig. 5.**
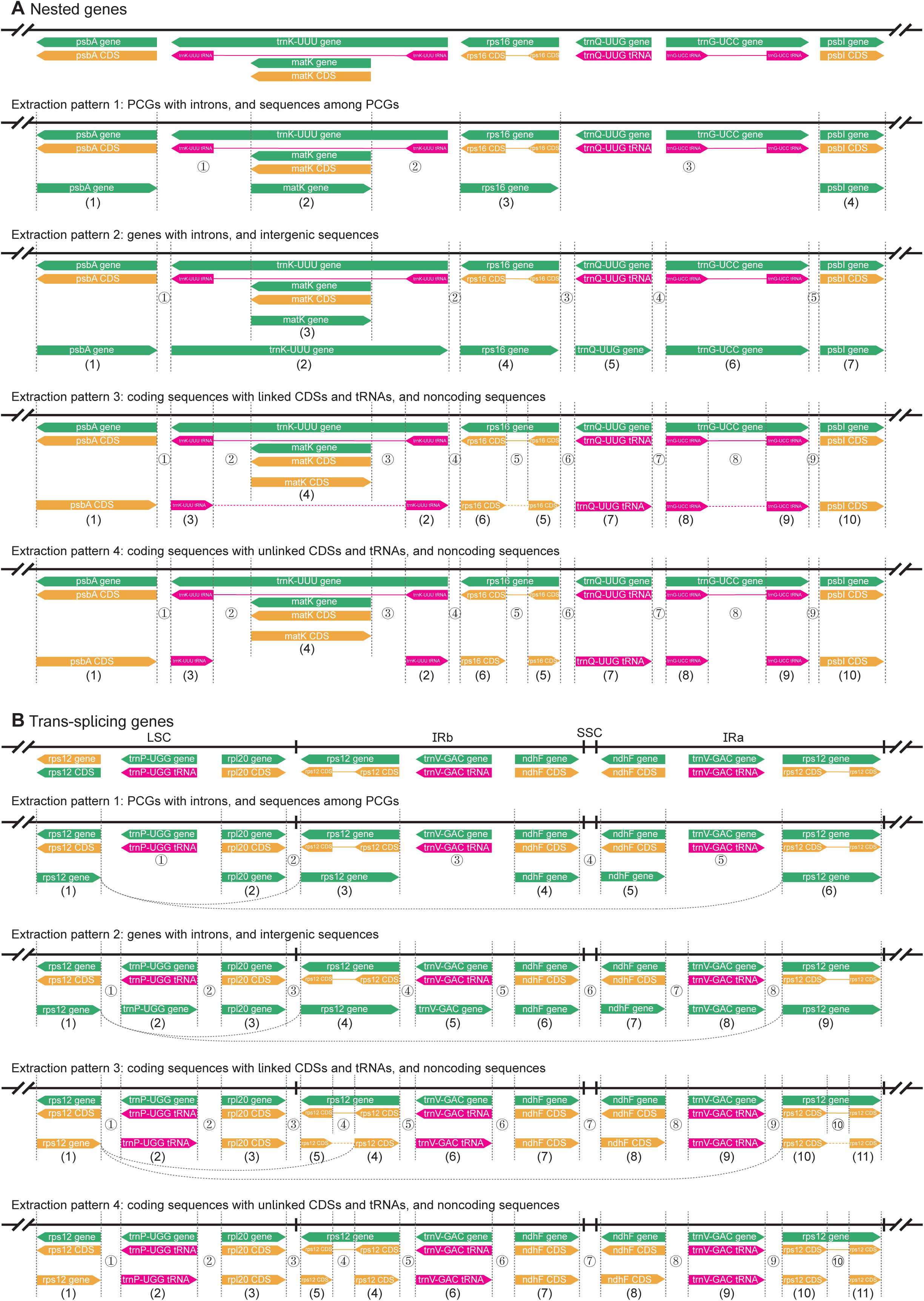
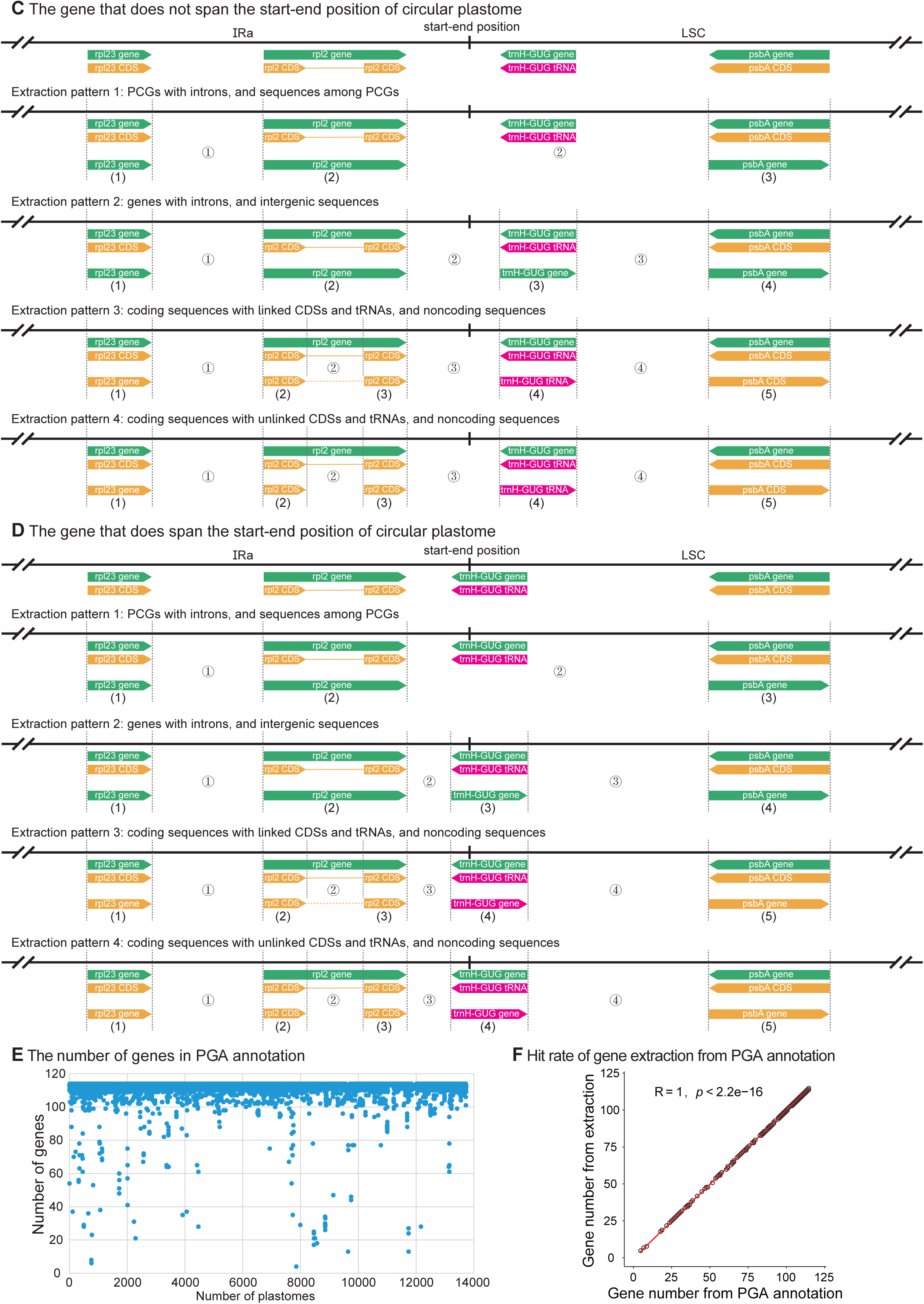
Four extraction patterns of canonical and special coding and noncoding sequences in plastomes. (A) Nested genes. Pattern 1: (1)—(4), PCGs; ①—③, sequences among PCGs. Pattern 2: (1)—(7), genes; ① — ⑤, intergenic sequences. Pattern 3: (1)—(10), coding sequences; ①—⑨, noncoding sequences. Exons of PCGs and tRNAs are connected with dash line. Pattern 4: (1)—(10), coding sequences; ①—⑨, noncoding sequences. Exons of PCGs and tRNAs are separately presented. (B) Trans-splicing genes. Pattern 1: (1)—(6), PCGs; ①—⑤, sequences among PCGs. Exon1 of *rps12* in LSC (1) and exon2, exon3, and intron of *rps12* in IRb (3) and Ira (6) are connected with curve line, respectively. Pattern 2: (1)—(9), genes; ①—⑧, intergenic sequences. Exon1 of *rps12* in LSC (1) and exon2, exon3, and intron of *rps12* in IRb (4) and IRa (9) are connected with curve line, respectively. Pattern 3: (1)—(11), coding sequences; ①—⑩, noncoding sequences. Exon1 of *rps12* in LSC (1) and exon2 (4) and exon3 (5) of *rps12* in IRb and exon2 (10) and exon3 (11) of *rps12* in IRa are connected with curve line, respectively. Pattern 4: (1)—(11), coding sequences; ① —⑩, noncoding sequences. Exon1 of *rps12* in LSC (1) and exon2 (4) and exon3 (5) of *rps12* in IRb and exon2 (10) and exon3 (11) of *rps12* in IRa are separately presented. (C) The gene that does not span the start-end position of circular plastome. Pattern 1: (1)—(3), PCGs; ① and ②, sequences among PCGs. Pattern 2: (1)—(4), genes; ① —③, intergenic sequences. Pattern 3: (1)—(5), coding sequences; ①—④, noncoding sequences. Exons of PCGs and tRNAs with introns are connected with dash line. Pattern 4: (1)—(5), coding sequences; ①—④, noncoding sequences. Exons of PCGs and tRNAs with introns are separately presented. (D) The gene that does span the start-end position of circular plastome. Pattern 1: (1)— (3), PCGs; ① and ②, sequences among PCGs. Pattern 2: (1)—(4), genes; ①—③, intergenic sequences. Pattern 3: (1)—(5), coding sequences; ① — ④, noncoding sequences. Exons of PCGs and tRNAs with introns are connected with dash line. Pattern 4: (1)—(5), coding sequences; ①—④, noncoding sequences. Exons of PCGs and tRNAs with introns are separately presented. (E) The number of genes in PGA annotation. The order of accession numbers was sorted from small to large. (F) Hit rate of gene extraction from PGA annotation. Gene number from extraction was significantly positively correlated with gene number from PGA annotation.

To evaluate the extraction tool, the gene numbers in PGA annotation were applied (Fig. 5E). Using *A. trichopoda* (angiosperm) as reference, the 13,731 angiosperm plastid sequences were re-annotated using PGA v2.0 with default parameter values, and the number of the annotated genes in these 13,731 angiosperm plastid sequences represented PGA annotation. The results indicated that the number of genes in PGA annotation were concentrated around 110, which was consistent with the common gene numbers in angiosperm plastomes (Jansen and Ruhlman, 2012; Wicke et al., 2011). Furthermore, the hit rate of gene extraction from PGA annotation was used to evaluate the accuracy of the extraction tool in PlastidHub (Fig. 5F). The gene number from extraction was significantly positively correlated with the gene number from PGA annotation (Pearson correlation: R=1, *p*<2.2e-16). Importantly, the difference in the number of extracted genes and PGA-annotated genes was minimal, showing at most one-gene difference. These results collectively indicated that the extraction tool in PlastidHub was accurate and effective in extracting genes from PGA annotations. Finally, it may be the state-of-the-art tool that was specifically designed for extracting canonical and special coding and noncoding sequences from plastome sequences, and can be applied to all users who need to extract any type of sequences from the plastomes.

### 3.7. Main function 5: Automatic screening of molecular markers using the Barcoding tool

DNA barcoding is an efficient method for species identification, molecular phylogenetics, population genetics, phylogeography, conservation genetics, and biodiversity studies. For plant barcoding efforts employing the plastome, screening molecular markers is always an indispensable step, but doing this in automated fashion can be challenging.

In this study, we developed a new tool that can be used to calculate and extract variable, invariable, and gap sites from batch submissions of sequence alignments (Fig. 6). Automatic screening of molecular markers by considering variable sites, parsimony informative sites, and total alignment length is an effective batch processing approach that does not rely on third-party tools, allowing users to more quickly move to downstream applications and analyses. Specifically, both the number and the percentage of variable sites/parsimony informative sites within each alignment are calculated. In addition, both the nucleotides of variable sites/parsimony informative sites and their position within each alignment are extracted. Most importantly, the alignments can be sorted according to percentage of variable sites or parsimony informative sites, which can help users more effectively identify informative markers. Variable sites or parsimony informative sites for each sequence in PCGs, RNAs, introns, and intergenic regions are separately displayed and sorted (Fig. 6A-6H). Total alignment length can be used to screen molecular markers (Fig. 6A-6H), because appropriate sequence length is an important prerequisite for designing primers (Ye et al., 2012). For both variable sites and parsimony informative sites, boxplots of PCGs, RNAs, tRNAs, rRNAs, introns, and intergenic regions showed that intergenic regions, introns, and PCGs are the three top choices, with more variation than RNAs, tRNAs, and rRNAs (Fig. 6I), which reflects differences in evolutionary rates among different genomic regions (Magee et al., 2010; Wolfe et al., 1987).

**Fig. 6.**
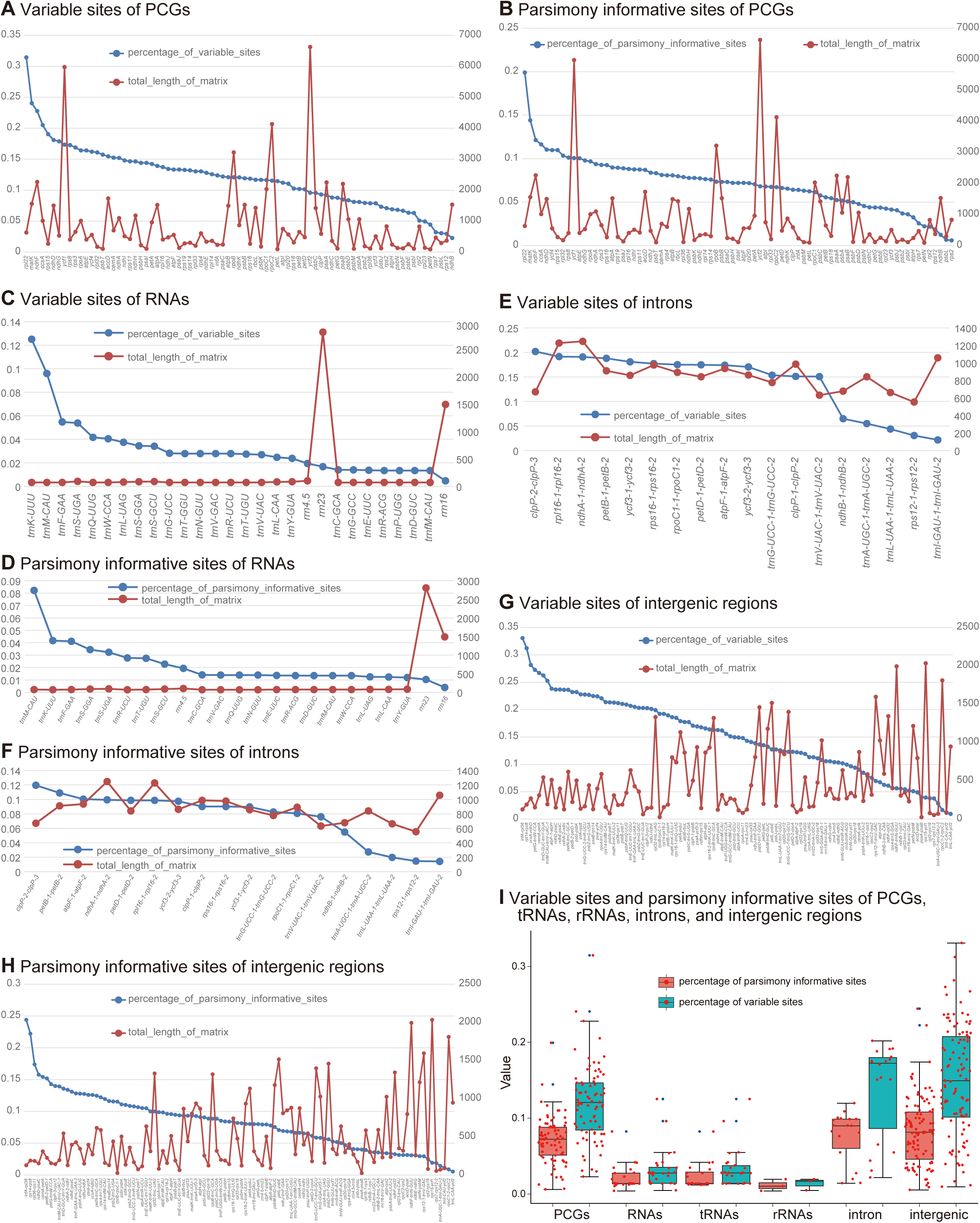
Automatically screening molecular markers for plastomes. (A, C, E, G) Variable sites of PCGs, RNAs, introns, and intergenic regions, respectively. The alignment matrices are sorted based on percentage of parsimony informative sites. Total lengths of alignment matrices are displayed synchronously. (B, D, F, H) Parsimony informative sites of PCGs, RNAs, introns, and intergenic regions, respectively. (I) Boxplots of variable sites and parsimony informative sites for PCGs, RNAs, tRNAs, rRNAs, introns, and intergenic regions. Red points are shake scatter points of all sequences, and blue points are outlier values of sequences.

### 3.8. Main function 6: Micro-synteny structural comparison using the Gene Homology tool

Genomic alignment at the whole plastome level is a common practice, and genome-scale alignments should allow visualization of regions with high vs. low levels of variation. VISTA (Frazer et al., 2004) is one of the most widely used tools for plastome comparison. However, it is not easy for non-experienced users to generate the input file format for mVISTA analysis. This was the motivation for developing the tool “generate mVISTA”, generates mVISTA alignments from GenBank flatfiles. There are other tools available for comparative analysis of plastomes, such as progressiveMAUVE (Darling et al., 2010), which are widely applied in plastomes with structural variations. However, tools that can perform micro-synteny comparison of plastomes are still lacking.

We developed a new tool called “gene homology” that can be used to generate a circular map of two plastomes with homologous genes being connected and non-shared genes being marked (Fig. 7). This tool has the function of pairwise visual comparison of each plastid gene, ultimately displaying micro-synteny structural variations, and evaluating annotation completeness at the gene-level. It can not only display structural variations such as inversions, translocations, and duplications, but also show the degree of annotation completeness considering gene losses/gains. Whether the reference and target plastomes have IR copies or not, and whether they have two IR copies or one IR copy, this tool considers all potential scenarios of plastome structure (Fig. 7A). To demonstrate the flexibility of this tool (Fig. 7B), we show plastome comparisons of *Amborella trichopoda* against other angiosperms (*Rosa roxburghii*, *Salicornia bigelovii*) and a gymnosperm species (*Calocedrus decurrens*), as well as a comparison of the gymnosperm species *Calocedrus decurrens* and *Calocedrus rupestris*. As far as we know, this is the first tool for pairwise visualization of plastomes at the gene level in order to show micro-synteny structural variations and evaluate annotation completeness.

**Fig. 7.**
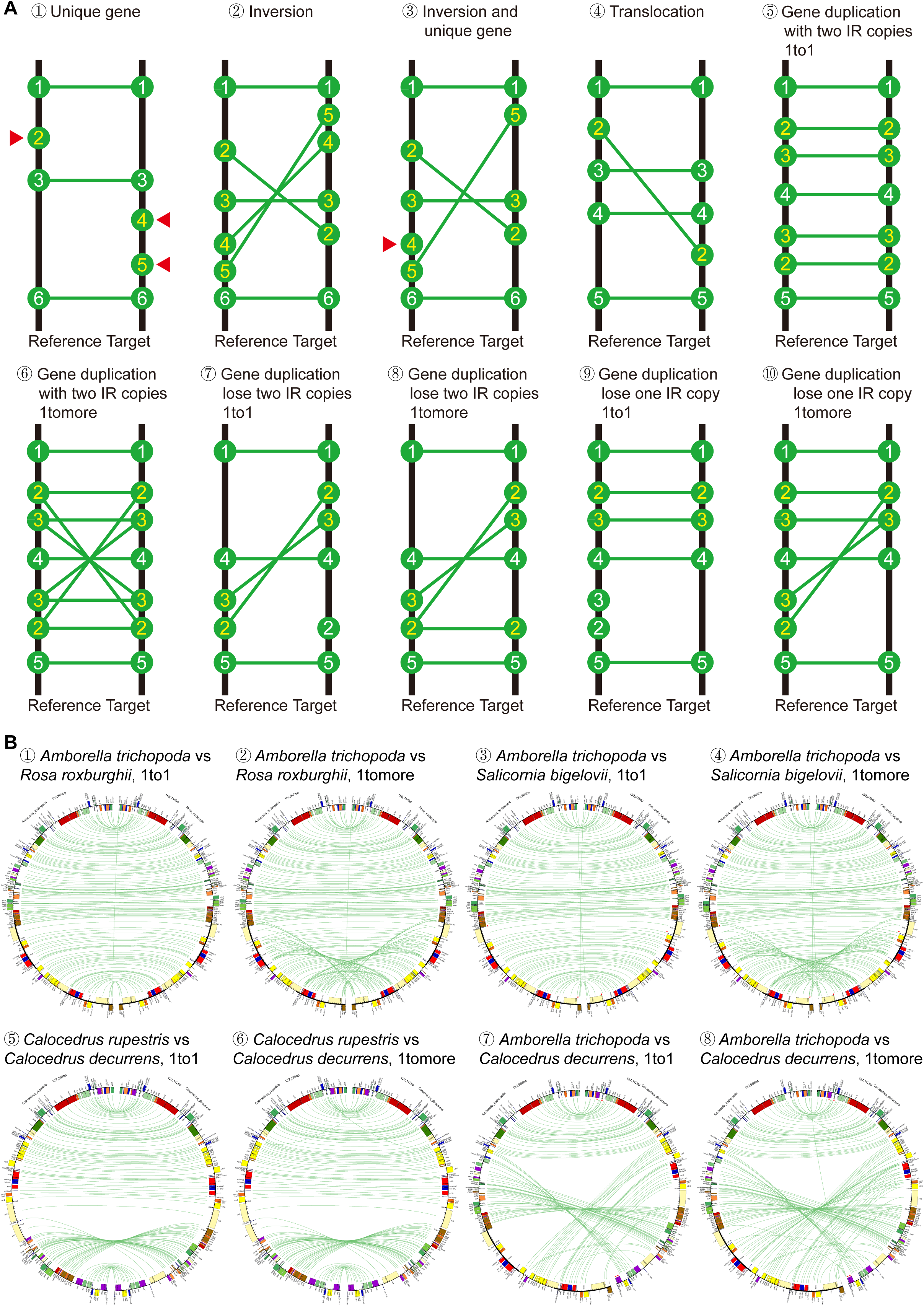
Micro-synteny comparision of each gene in target and reference plastomes. (A) Principle. ① Unique gene. Unique genes in target or reference plastome will be marked with red triangle symbol. ② Inversion. Inversion can be shown by connecting genes in reciprocally inverted segments. ③ Inversion and unique gene. Not all genes in inverted segments are shared between target and reference, so unique genes in inverted segments are marked with red triangle symbol. ④ Translocation. ⑤ Gene duplication, with two IR copies, 1to1. The tool only connects shared genes with same order. ⑥ Gene duplication, with two IR copies, 1tomore. ⑦ Gene duplication, lose two IR copies, 1to1. ⑧ Gene duplication, lose two IR copies, 1tomore. ⑨ Gene duplication, lose one IR copy, 1to1. ⑩ Gene duplication, lose one IR copy, 1tomore. For ⑥, ⑧, and ⑩, the tool connects shared genes with both same and opposite order. For ⑦ and ⑨, the tool only connects shared genes with same or opposite order. (B) Examples. ① *Amborella trichopoda* vs *Rosa roxburghii*, 1to1. ② *A. trichopoda* vs *R. roxburghii*, 1tomore. For ① and ②, no IR copies are lost from both species. ③ *A. trichopoda* vs *Salicornia bigelovii*, 1to1. ④ *A. trichopoda* vs *S. bigelovii*, 1tomore. For ③ and ④, no IR copies are lost from both species. ⑤ *Calocedrus rupestris* vs *C. decurrens*, 1to1. ⑥ *C. rupestris* vs *C. decurrens*, 1tomore. For ⑤ and ⑥, one IR copy is lost from both species. ⑦ *A. trichopoda* vs *C. decurrens*, 1to1. ⑧ *A. trichopoda* vs *C. decurrens*, 1tomore. For ⑦ and ⑧, one IR copy is lost from *C. decurrens*.

### 3.9. Similar functions with other tools: Submission and Visualization

As far as we know, there are two tools that can convert GenBank flatfiles to the tbl-formatted files required for NCBI/EMBL/DDBJ database submission: gbf2tbl.pl (NCBI) and GeSeq (Tillich et al., 2017). There are six tools that can perform plastome visualization: OGDRAW v1.3.1 (Greiner et al., 2019), CGView (Stothard et al., 2019), CPGAVAS2 (Shi et al., 2019), Chloroplot (Zheng et al., 2020), Chloe (https://chloe.plastid.org/annotate.html), and CPGView (Liu et al., 2023). GeSeq may be the only tool that can conduct both format conversion and gene visualization of annotated plastomes. Our new platform, PlastidHub, provides user-friendly tools for plastome submission and visualization. The visualization tool in PlastidHub also has a function to show the linked dispersed repeats including forward and inverted repeats, and different repeats that meet user-specified requirements can be displayed by adjusting parameters such as minimum allowed percent identity, minimum allowed length, and BLAST e-value.

### 3.10. Streamlining analysis with personalized new functions in three toolkits: Pre-Alignment, (Post)-Alignment, and Phylogeny

In this section, we describe five practical and flexible tools that can be used as part of a customized phylogenetic inference pipeline: “sort flanking genes”, “check bad codons”, “check missing taxa”, “extract sub-alignments”, and “RAxML gene tree”. The tool “sort flanking genes” was specifically developed for plastomes with obvious structural rearrangements, such as inversions and translocations. This is because it is essential to adjust the orientations of intergenic regions (to the same direction) before alignment and phylogenetic reconstruction. Thus, this tool will improve the alignment accuracy of intergenic regions in plastomes, and it can also be used to track the origin and evolution of gene remnants in intergenic regions, suitable for users with personalized analysis needs. The tool “check bad codons” can check for bad codons in batch submissions of PCG matrices, allowing users to remove unqualified sequences to improve alignment accuracy. We all know that correct PCG matrices do not include internal bad codons, so bad codons are signals that may contain errors in PCGs. The tool “check missing taxa” searches for missing taxa in batch submissions of multiple sequence matrices. This tool will output a two-dimensional table showing coverage of species and sequences, helping users evaluate the completeness of their dataset and potential false-positive problems involving particular species or plastid regions. The tool “extract sub-alignments” uses an input list of species names to generate sub-alignments from batch submissions of sequence alignments, while simultaneously deleting gap sites in the sub-alignments. This tool is important for users who want to test the impact of species sampling on phylogenetic inference. The tool “RAxML gene tree” conducts ML phylogenetic inference using RAxML on batch submissions of sequence alignments, which is useful for users who want to reconstruct gene and species trees to explore phylogenetic conflict.

### 3.11. Help documents and notes

To facilitate use of this new platform, we provide detailed user guides, test example files, and warning prompts when entering unreasonable parameter values. Help documents can be easily acquired from the document pages corresponding with each tool. Structured help documents can also be accessed in the supplemental information. Notes and warning prompts for each tool are shown in the tool interface. In general, use of unreasonable parameter values will prompt an error warning encouraging use of values within a specified range. Warnings will also appear in cases where files are incorrectly uploaded (e.g., due to wrong file format) or due to restrictions in file number or size. These educational resources will facilitate seamless and efficient use of PlastidHub, helping scientists execute their research goals.

## 4. Conclusions

Although some previously released tools, e.g., GeSeq (Tillich et al., 2017), possess similar functionalities as those in PlastidHub, none has nearly as many customizable functions. PlastidHub currently offers a set of 25 tools covering various aspects of plastome analyses. The majority of tools in PlastidHub (19/25) do not rely on third-party tools; only six of 25 tools in PlastidHub rely on third-party tools as BLAST+ v2.16.0 (Camacho et al., 2009), Circos v0.66 (Krzywinski et al., 2009), MAFFT v7.3 (Katoh and Standley, 2013), and RAxML v8.2.10 (Stamatakis, 2014). PlastidHub will be continuously updated, with new versions including additional customizable tools incorporating emerging methods and accommodating the diverse and expanding needs of the plastome research community.

PlastidHub is a cloud-based web application, meaning users do not need to install, update, or maintain software or invest in extensive computational resources in order to conduct analyses. As a web platform, improving user experience is the top priority of PlastidHub. The ability to switch between the tool interface and the help/download interface is a practical feature of PlastidHub, allowing users to switch between key webpages as their design and execute their workflows. User feedback and other developments in the field will be used to inform daily updates and the development of additional tools on the website.

## Data availability statement

All relevant data and code can be found within the article and on the PlastidHub website (https://www.plastidhub.cn) and GitHub (https://github.com/quxiaojian/PlastidHub).

## CRediT authorship contribution statement

**Na-Na Zhang:** Data curation; Formal analysis; Visualization; Writing—original draft. **Gregory W. Stull:** Investigation; Supervision; Writing—review & editing. **Xue-Jie Zhang:** Visualization; Writing—review & editing. **Shou-Jin Fan:** Conceptualization; Supervision; Writing—review & editing. **Ting-Shuang Yi:** Conceptualization; Supervision; Funding acquisition; Writing—original draft; Writing—review & editing. **Xiao-Jian Qu:** Conceptualization; Supervision; Funding acquisition; Formal analysis; Visualization; Writing—original draft; Writing—review & editing.

## Acknowledgements

We thank Dan Zou, Yu-Liang Chen, Zhen-Xiang Feng, Yuan-Ting Gu, and Rui-Yu Zhang for testing the PlastidHub. We thank the editors and reviewers for constructive comments and suggestions on improving this study. This study was funded by the Natural Science Foundation of Shandong Province (ZR2020QC022), the Science and Technology Basic Resources Investigation Program of China (no. 2019FY100900), and the open research project of “Cross Cooperative Team” of the Germplasm Bank of Wild Species, Kunming Institute of Botany, Chinese Academy of Sciences.

