## Supplementary files for "PlastidHub: an integrated analysis platform for plastid phylogenomics and comparative genomics"

### CONTENTS


**1. Annotation** ... 3

**1.1. Quadripartition** ... 3

**1.2. Annotation** ... 7

**2. Assessment** ... 12

**2.1. Assess Gene Number** ... 12

**2.2. Assess Gene Length** ... 15

**3. Submission** ... 18

**4. Visualization** ... 22

**5. Extraction** ... 30

**6. Pre-Alignment** ... 40

**6.1. Sort Genenames** ... 40

**6.2. Remove Duplications** ... 42

**6.3. Combine Fasta** ... 44

**6.4. Generate Matrices** ... 45

**6.5. Check Codons** ... 47

**6.6. Check Missings** ... 48

**6.7. Extract Sites** ... 49

**6.8. Codon-to-AA** ... 51

**7. (Post)-Alignment** ... 52

**7.1. Sequence Alignment** ... 52

**7.2. Generate Non-interleaved** ... 54

**7.3. Extract Sub-alignments** ... 55

**7.4. Concatenate Alignments** ... 58

**7.5. Remove Gaps** ... 60

**7.6. Generate PartitionFinder** ... 63

**8. Phylogeny** ... 64

**9. Barcoding** ... 66

**10. Plastomics** ... 71

**10.1. Generate mVISTA** ... 71

**10.2. Gene Homology** ... 73

**1. Annotation**

**Functionality**

This tool kit contains two tools that can complete standardization of quadripartite structure and plastome annotation, such as "1.1. Quadripartition and 1.2. Annotation".

**1.1. Quadripartition**

***Functionality***

This tool can batchly adjust the quadripartite structure of plastome from a non-standardized form to a standardized form (LSC-IRb-SSC-IRa). Due to the fact that some plastome sequences generated in the laboratory or downloaded from NCBI are always in non-standardized form or in opposite direction, it is necessary to adjust the structures and orientations of these plastome sequences for downstream analysis, including plastome annotation, map drawing, comparative genomics, etc.

***Features***

1. Standardization of quadripartite structures can be batchly adjusted for plastome sequences.
2. Plastome sequences in opposite direction can be batchly reverse complemented (-r).
3. The length and nucleotide of IRb and IRa can be set to be exactly identical or nearly identical (-s).
4. The minimum allowed IR length can be adjusted by the users (-l).
5. The result file Amborella_trichopoda.fasta contains plastome sequence with a standardized form (LSC-IRb-SSC-IRa) after adjustment. The result file Amborella_trichopoda_LSC_IRb_SSC_IRa.fasta contains each LSC, IRb, SSC, and IRa sequence after adjustment.
6. The result file quadripartite_structure_coordinate.txt contains coordinates of four fragments (LSC, IRb, SSC, and IRa) for pre-adjusted input files and post-adjusted output files.
7. The result file warning.txt contains all abnormal situations assisting users to check the nucleotides, length, and orientation of IRb and IRa. The result file screen.log contains screen output that can help developer debug.

***Command-line Tool***

perl quadripartite_standardization.pl -i -r -s -l -o

run:
perl quadripartite_standardization.pl -i input -r N -s N -l 1000 -o output
perl quadripartite_standardization.pl -i input -r N -s Y -l 1000 -o output
perl quadripartite_standardization.pl -i input -r Y -s N -l 1000 -o output
perl quadripartite_standardization.pl -i input -r Y -s Y -l 1000 -o output

parameter:
[-i -input] required: (default: input) input directory name containing FASTA files.
[-r -rc] required: (default: N) if N, entire sequence does not require reverse-complement; if Y, entire sequence requires reverse-complement.
[-s -same] required: (default: N) if N, length and nucleotide of IRb and IRa may not be

exactly the same, allowing for a certain degree of sequence difference; if Y, length and nucleotide of IRb and IRa must be exactly the same.

[-l -length]    required: (default: 1000) minimum allowed inverted-repeat (IR) length.

[-o -output]    required: (default: output) output directory name.

***Principles***

The adjustment of plastome quadripartite structure to a standardized form is shown in Figure 1.

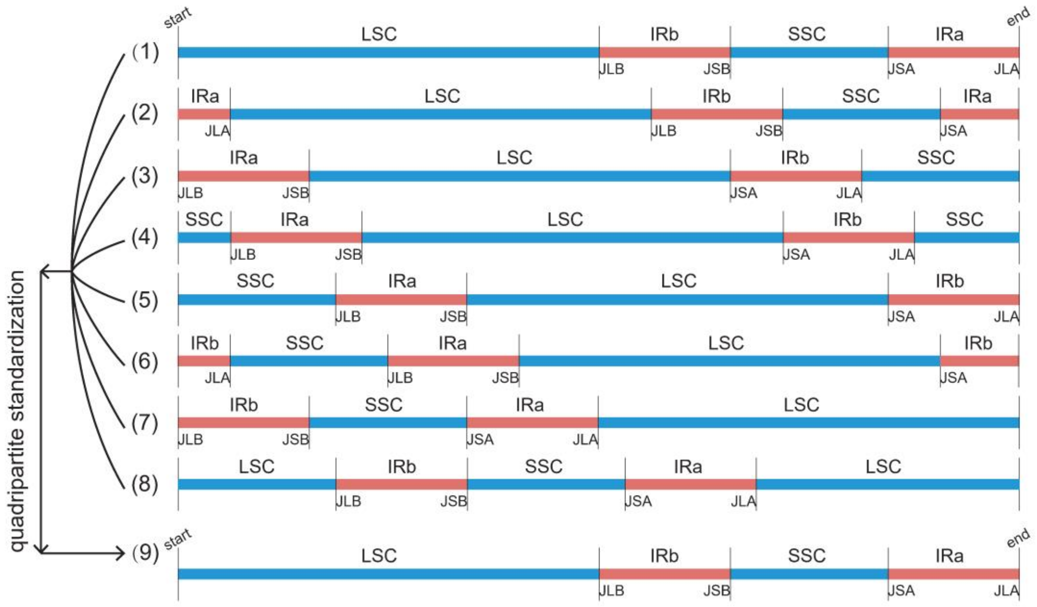


Figure 1. Standardization of quadripartite structure for plastome sequences (LSC-IRb-SSC-IRa).
(1)-(8) Plastome quadripartite structure before adjustment.
(1) Quadripartite structure is exactly the right one.
(2) Quadripartite structure is not a standardized form, due to start-end point in IRa.
(3) Quadripartite structure is not a standardized form, due to start-end point in junction of SSC-IRa.
(4) Quadripartite structure is not a standardized form, due to start-end point in SSC.
(5) Quadripartite structure is not a standardized form, due to start-end point in junction of IRb-SSC.
(6) Quadripartite structure is not a standardized form, due to start-end point in IRb.
(7) Quadripartite structure is not a standardized form, due to start-end point in junction of LSC-IRb.
(8) Quadripartite structure is not a standardized form, due to start-end point in LSC.
(9) Plastome quadripartite structure after adjustment. In addition, the sequence direction of plastomes can be reverse complemented; the length and nucleotide of two IR copies can be set to be exactly or nearly identical; and the minimum allowed IR length can be changed.

***Example***

1. Input -> Amborella_trichopoda.fasta, Rosa_roxburghii.fasta

Amborella_trichopoda.fasta

```
>Amborella_trichopoda
CCTTGATCCACTTGGCTACATCCGCCCATACCTATCTTTCTATTTTCTATAAATCTCTAGGAAAA
```

2. Output -> Amborella_trichopoda.fasta, Rosa_roxburghii.fasta

Amborella_trichopoda.fasta

```
>Amborella_trichopoda
GGGGATTGAACCCGCGCATGGTGGATTCACAATCCACTGCCTTGATCCACTTGGCTACATCCGCC
```

3. Output -> Amborella_trichopoda_LSC_IRb_SSC_IRa.fasta, Rosa_roxburghii _LSC_IRb_SSC_IRa.fasta

Amborella_trichopoda_LSC_IRb_SSC_IRa.fasta

```
>Amborella_trichopoda_LSC
GGGGATTGAACCCGCGCATGGTGGATTCACAATCCACTGCCTTGATCCACTTGGCTACATCCGCC
>Amborella_trichopoda_IRb
GTCGTTCGCCCGACTTGTTCTATTTACTTACGACGACGAAGAATGAAACTATCACTATATTTCTT
>Amborella_trichopoda_SSC
ATATAGATAGACTAAATATAGACTAAATAAACATAGACATAGACTCAAAAATACAAATAAAAATA
>Amborella_trichopoda_IRa
ATTTTGGGTAAGCATTCTTTTTCATCCCGATCATTGCATAATCTGACCCTTTTTTGGAGAACAAT
```

4. Output -> quadripartite_structure_coordinate.txt

```
FastaFileNames  AdjustmentForms LSC(JLA-JLB)    IRb(JLB-JSB)    SSC(JSB-JSA)    IRa(JSA-JLA)
Amborella_trichopoda    Pre-adjustment  162648-90911    90912-117572    117573-135986   135987-162647
Amborella_trichopoda    Post-adjustment 1-90950 90951-117611    117612-136025   136026-162686
Rosa_roxburghii Pre-adjustment  12666-98517     98518-124570    124571-143361   143362-12665
Rosa_roxburghii Post-adjustment 1-85852 85853-111905    111906-130696   130697-156749
```

5. Output -> warning.txt

```
Amborella_trichopoda need to be checked for following reason(s):
Nucleotide of IRb and IRa may not be exactly the same!
IRb and IRa may be forward repeat rather than inverted repeat!

Amborella_trichopoda need to be checked for following reason(s):
Nucleotide and length of IRb and IRa may not be exactly the same!
IRb and IRa may be forward repeat rather than inverted repeat!

Amborella_trichopoda need to be checked for following reason(s):
Nucleotide and length of IRb and IRa may not be exactly the same!

Amborella_trichopoda need to be checked for following reason(s):
Minimum allowed IR length 100000 may be larger than actual maximum IR length!
```

6. Output -> screen.log

```


Building a new DB, current time: 03/24/2024 11:33:55
New DB name:   C:\Users\Administrator\Desktop\input\%E%NMQy3%mHMfqZ%_Amborella_trichopoda_IR_temp
New DB title:  input/%E%NMQy3%mHMfqZ%_Amborella_trichopoda_IR_temp
Sequence type: Nucleotide
Keep MBits: T
Maximum file size: 1000000000B
Adding sequences from FASTA; added 1 sequences in 0.0041527 seconds.
```

##### 1.2. Annotation

***Functionality***

PGA (Plastid Genome Annotator), a standalone command line tool, can perform rapid, accurate, and flexible batch annotation of newly generated target plastomes based on well-annotated reference plastomes. In contrast to current existing tools, PGA uses reference plastomes as the query and unannotated target plastomes as the subject to locate genes, which we refer to as the reverse query-subject BLAST search approach. PGA accurately identifies gene and intron boundaries as well as intron loss. The program outputs GenBank-formatted files as well as a log file to assist users in verifying annotations.

Following six steps will be conducted to annotate plastomes: (1) Preparation of GenBank-formatted reference plastomes; (2) Preparation of FASTA-formatted target plastomes; (3) Reference database generation; (4) BLAST search; (5) Determining feature boundaries; (6) Generating GenBank and log files.

***New Features***

1. Link or unlink trans-splicing gene *rps12* can be selected by users.

2. Annotate or unannotate redundant genes, such as nested genes and residual genes with short fragments, can be selected by users.

***Updated Version***

Now, we have updated PGA v1.0 to a newer version, i.e., PGA v2.0.

***Recommended Suggestions***

1. Users should carefully check the GenBank-formatted reference plastome. PGA is packaged with several properly annotated plastomes, and it is thus possible for users to use PGA to re-annotate a plastome that is intended to be used as a reference, in order to correct possible inaccuracies.

2. It is important that users select a reference plastome that contains sufficient numbers of annotated genes for the target taxa. The number of genes in the reference plastome(s) should equal or exceed the number in the target plastome(s). If the number of genes in the target is uncertain, it may be best to use multiple reference plastomes. The *Amborella trichopoda* (AJ506156) and *Zamia furfuracea* (JX416857) plastomes included within PGA are examples of plastomes that contain the highest gene numbers among known angiosperms and gymnosperms, and as such it is recommended that they be included as references during PGA runs.

3. We do not recommend annotating highly incomplete plastomes using a complete reference plastome, because BLAST may annotate some genes redundantly (i.e., BLAST may return hits for genes that were not sequenced or are otherwise absent in the incomplete plastome, resulting in spurious annotations). To annotate highly incomplete plastomes or plastome segments, we recommend using progressiveMauve (as implemented in Mauve 2.4.0; Darling et al., 2010) to align the incomplete plastome to the reference plastome, followed by the use of the corresponding homologous block of the reference plastome as the reference for annotation in PGA.

4. We suggest that users carefully check highly divergent or otherwise unusual target plastomes for incorrect annotations. This is particularly important for plastomes with a high degree of gene loss, pseudogenization or sequence divergence.

***Command-line Tool***

perl PGA_v2.pl –r –t [–i –p –l –d –q –o –f –w]

run:

perl PGA_v2.pl –r reference –t target

perl PGA_v2.pl –r reference –t target –i 1000 –p 60 –l Y –d N –q 0.5,2.0 –o gb –f circular –w warning

parameter:

[-r -reference] required: (default: reference) input directory name containing GenBank-formatted file(s) that from the same or close families.

[-t -target] required: (default: target) input directory name containing FASTA-formatted file(s) that will be annotated.

[-i -ir] optional: (default: 1000) minimum allowed inverted-repeat (IR) length.

[-p -pidentity] optional: (default: 60) any PCGs with a TBLASTN percent identity less than this value will be listed in the log file and will not be annotated.

[-l -link] optional: (default: Y) if Y, the exons of the trans-splicing gene *rps12* will be linked; if N, the exons of the *rps12* gene will not be linked.

[-d -redundancy] optional: (default: N) if N, any PCGs with a query coverage per annotated PCG less or greater than each of the two values (<1,>1) in the following parameter will not be annotated; if Y, any PCGs with a query coverage per annotated PCG less or greater than each of the two values (<1,>1) in the following parameter will be annotated, which can be used to trace the evolutionary remains (pseudo-genes) of functional PCGs.

[-q -qcoverage] optional: (default: 0.5,2.0) any PCGs with a query coverage per annotated PCG less or greater than each of these two values (<1,>1) will be listed in the log file.

[-o -out] optional: (default: gb) output directory name.

[-f -form] optional: (default: circular) circular or linear form for FASTA-formatted file.

[-w -warning] optional: (default: warning) log file name containing warning information for annotated GenBank-formatted file(s).

***Principles***

Improvement of annotation personalization for the plastome is shown in Figure 1.

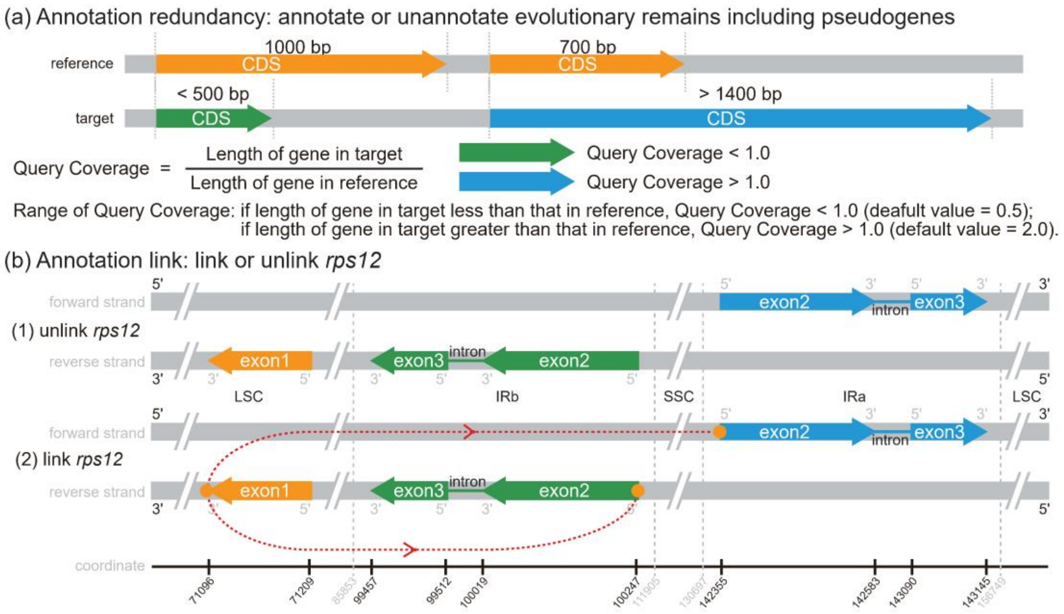


Figure 1. Improvement of annotation personalization for the plastome.

(a) Annotation redundancy: annotate or unannotate evolutionary remains including pseudogenes. The evolutionary remains (e.g., pseudogenes) can be annotated or unannotated by changing the dafault value (N: unannotate pseudogenes) of the parameter in PGA v2.0. Specifically, the detection threshold can be changed by adjusting another parameter with two flexibly changeable values (Query Coverage < 1.0, Query Coverage > 1.0). If a gene with a query coverage less or greater than each of these two manually set values (default value = 0.5, default value = 2.0), this gene will be preliminarily judged as a pseudogene or redundant false-positive sequence fragment, waiting for further confirmation by the user.

(b) Annotation link: link or unlink *rps12*. The trans-splicing gene *rps12* can be linked or unlinked by changing the default value (Y: link *rps12*) of the parameter in PGA v2.0.

***Example***

1. Input -> Amborella_trichopoda.gb

```
LOCUS       Amborella_trichopoda      162686 bp    DNA     circular UNA 08-JUN-2015
DEFINITION  Amborella trichopoda chloroplast genomic DNA, complete sequence.
ACCESSION   AJ506156
VERSION     AJ506156.2  GI:34481608
KEYWORDS    complete genome.
SOURCE      chloroplast Amborella trichopoda
  ORGANISM  Amborella trichopoda
            Eukaryota; Viridiplantae; Streptophyta; Embryophyta; Tracheophyta;
            Spermatophyta; Magnoliophyta; basal Magnoliophyta; Amborellales;
            Amborellaceae; Amborella.
FEATURES             Location/Qualifiers
     source          1..162686
                    /organism="Amborella trichopoda"
                    /mol_type="genomic DNA"
     repeat_region    90951..117611
                    /note="inverted repeat region B; IRB repeat region"
                    /rpt_type="inverted"
     rRNA           complement(139284..142097)
                    /gene="rrn23"
                    /product="23S ribosomal RNA"
     gene           complement(139284..142097)
                    /gene="rrn23"
     tRNA           join(complement(4472..4508), complement(1840..1874))
                    /gene="trnK-UUU"
                    /product="tRNA-Lys"
     gene           complement(1840..4508)
                    /gene="trnK-UUU"
     CDS            join(complement(16186..16330), complement(14506..14915))
                    /gene="atpF"
                    /codon_start=1
                    /transl_table=11
                    /product="ATPase I subunit"
                    /translation="MKNVTDSFVSLGHWPSAGSFGFNTDIFATNPINLSVVLGVLIFF
                    GKGVLSDLLDNRKQRILSTIRNSEELRGGAIEQLEKARARLRKVEIEADEFRVNGYSE
                    IEREKSNLINAAYENLERLENYKNESIHFEQQRAMNQVRQRVFQQALQGALETLNSYL
                    NSELHLRTISANIGMLGTMKNITD"
     gene           complement(14506..16330)
                    /gene="atpF"
```

2. Input -> Rosa_roxburghii.fasta

```
>Rosa_roxburghii
ATGGGCGAACGACGGGAATTGAACCCGCGCGTGGTGGATTCACAATCCACTGCCTTGATC
```

3. Output -> Rosa_roxburghii.gb

```
LOCUS       Rosa_roxburghii  156749 bp    DNA     circular PLN 25-DEC-2017
FEATURES             Location/Qualifiers
     source          1..156749
                     /organism="Rosa_roxburghii"
                     /mol_type="genomic DNA"
     gene            106222..109027
                     /gene="rrn23"
     rRNA            106222..109027
                     /gene="rrn23"
                     /product="23S ribosomal RNA"
     gene            complement(1704..4278)
                     /gene="trnK-UUU"
     tRNA            join(complement(4242..4278), complement(1704..1738))
                     /gene="trnK-UUU"
                     /product="tRNA-Lys"
     gene            complement(12213..12767)
                     /gene="atpF"
     CDS             complement(12213..12767)
                     /gene="atpF"
                     /codon_start=1
                     /transl_table=11
                     /product="ATP synthase CF0 subunit I"
```

4. Output -> warning.log

```
Rosa_roxburghii
Warning: atpF (negative one-intron PCG) lost intron!
Total number of genes in the reference plastome(s): 114.
Total number of genes annotated in the target plastome: 113.
All gene names from the reference plastome(s) that were not annotated in the target plastome:
infA
```

#### 2. Assessment

***Functionality***

This tool kit contains two tools that can assess plastome annotation quality, such as "2.1. Assess Gene Number, and 2.2. Assess Gene Length".

##### 2.1. Assess Gene Number

***Functionality***

This tool can assess annotation completeness of plastomes as number of hitting, missing, and redundant genes. That is to say, annotation completeness of plastomes can be assessed by comparing numbers and names between genes in the target plastome and their corresponding genes in the reference plastome.

***Features***

1. Only one reference plastome and one target plastome can be compared at a time.
2. Assessment results highly depend on the reference plastome selected by the user.
3. Number of hitting, missing, and redundant genes are listed. Number and name of hitting, missing, and redundant PCGs, tRNAs, and rRNAs are also listed.
4. Since plastomes do not contain many genes (~100), it would be reasonable for users to manually check the status (hitting vs. missing vs. redundant) for each gene. Users then can determine why genes are absent (true loss vs. missing annotation for a gene that is actually present) or redundant (true gain vs. wrong annotation for a gene that is actually absent).

***Command-line Tool***

perl assess_gene_number.pl –r –t –o

run:

perl assess_gene_number.pl –r reference –t target –o output

parameter:

[-r -reference] required: (default: reference) input reference directory containing one GenBank-formatted file.

[-t -target] required: (default: target) input target directory containing GenBank-formatted file(s).

[-o -output] required: (default: output) output directory containing TXT-format file(s) with statistic result.

***Example***

1. Input -> Amborella_trichopoda.gb

```
LOCUS       Amborella_trichopoda      162686 bp    DNA     circular UNA 08-JUN-2015
DEFINITION  Amborella trichopoda chloroplast genomic DNA, complete sequence.
ACCESSION   AJ506156
VERSION     AJ506156.2  GI:34481608
KEYWORDS    complete genome.
SOURCE      chloroplast Amborella trichopoda
  ORGANISM  Amborella trichopoda
            Eukaryota; Viridiplantae; Streptophyta; Embryophyta; Tracheophyta;
            Spermatophyta; Magnoliophyta; basal Magnoliophyta; Amborellales;
            Amborellaceae; Amborella.
FEATURES             Location/Qualifiers
     source          1..162686
                    /organism="Amborella trichopoda"
                    /mol_type="genomic DNA"
     repeat_region    90951..117611
                    /note="inverted repeat region B; IRB repeat region"
                    /rpt_type="inverted"
     rRNA          complement(139284..142097)
                    /gene="rrn23"
                    /product="23S ribosomal RNA"
     gene           complement(139284..142097)
                    /gene="rrn23"
     tRNA          join(complement(4472..4508), complement(1840..1874))
                    /gene="trnK-UUU"
                    /product="tRNA-Lys"
     gene           complement(1840..4508)
                    /gene="trnK-UUU"
     CDS           join(complement(16186..16330), complement(14506..14915))
                    /gene="atpF"
                    /codon_start=1
                    /transl_table=11
                    /product="ATPase I subunit"
                    /translation="MKNVTDSFVSLGHWPSAGSFGFNTDIFATNPINLSVVLGVLIFF
                    GKGVLSDLLDNRKQRILSTIRNSEELRGGAIEQLEKARARLRKVEIEADEFRVNGYSE
                    IEREKSNLINAAYENLERLENYKNESIHFEQQRAMNQVRQRVFQQALQGALETLNSYL
                    NSELHLRTISANIGMLGTMKNITD"
     gene           complement(14506..16330)
                    /gene="atpF"
```

2. Input -> Rosa_roxburghii.gb, Rosa_chinensis.gb, Rosa_rugosa.gb

Rosa_roxburghii.gb

```
LOCUS       Rosa_roxburghii  156749 bp    DNA     circular PLN 02-DEC-2022
FEATURES             Location/Qualifiers
     source          1..156749
                     /organism="Rosa_roxburghii"
                     /organelle="plastid:chloroplast"
                     /mol_type="genomic DNA"
                     /note="Annotation Method :: PGA-Plastid Genome Annotator"
     repeat_region   85853..111905
                     /note="inverted repeat B"
                     /rpt_type="inverted"
     repeat_region   complement(130697..156749)
                     /note="inverted repeat A"
                     /rpt_type="inverted"
     gene            58781..60244
                     /gene="accD"
     CDS             58781..60244
                     /gene="accD"
                     /codon_start=1
                     /transl_table=11
                     /product="acetyl-CoA carboxylase carboxyltransferase beta subunit"
     gene            complement(10635..12158)
                     /gene="atpA"
     CDS             complement(10635..12158)
                     /gene="atpA"
                     /codon_start=1
                     /transl_table=11
                     /product="ATP synthase CF1 alpha subunit"
```

3. Output -> Amborella_trichopoda_Rosa_roxburghii.txt, Amborella_trichopoda_Rosa_chinensis.txt, Amborella_trichopoda_Rosa_rugosa.txt

Amborella_trichopoda_Rosa_roxburghii.txt

```
H:hitting, M:missing, R:redundant, n:number, p:percentage, P:PCGs, T:tRNAs, R:rRNAs

1. Number of hitting and missing genes:
|114 total gene numbers in reference|
|115 total gene numbers in target|

|114 number of hitting genes (nH)|
|0 number of missing genes (nM)|
|1 number of redundant genes (nR)|

|100% percentage of hitting gene (pH)|
|0% percentage of missing genes (pM)|
|1% percentage of redundant genes (pR)|

2. Number of hitting and missing PCGs, tRNAs and rRNAs:
|80 number of hitting PCGs (nHP)|
|0 number of missing PCGs (nMP)|
|1 number of redundant PCGs (nRP)|

|30 number of hitting tRNAs (nHT)|
|0 number of missing tRNAs (nMT)|
|0 number of redundant tRNAs (nRT)|

|4 number of hitting rRNAs (nHR)|
|0 number of missing rRNAs (nMR)|
|0 number of redundant rRNAs (nRR)|

3. Name of hitting and missing PCGs, tRNAs and rRNAs:
|name of hitting PCGs|
```

##### 2.2. Assess Gene Length

***Functionality***

This tool can assess annotation accuracy of genes by comparing gene length difference. That is to say, annotation accuracy of plastomes can be assessed by comparing length differences between genes in the target plastome and their corresponding genes in the reference plastome.

***Features***

1. Only one reference plastome and one target plastome can be compared at a time.
2. Assessment results highly depend on the reference plastome selected by the user.
3. Gene name, gene length in reference plastome, gene length in target plastome, and gene length difference are listed in type of PCGs, tRNAs, and rRNAs.
4. Gene length differences have been sorted by magnitude, and those with the largest differences (or those larger than a specified threshold) can be checked, according to users' own standards.
5. Since plastomes do not contain many genes (~100), it would be reasonable for users to manually check the status for each gene. Users then can determine why genes have such significant length differences.

***Command-line Tool***

perl assess_gene_length.pl –r –t –o

run:

perl assess_gene_length.pl –r reference –t target –o output

parameter:

[-r -reference] required: (default: reference) input reference directory containing one GenBank-formatted file.

[-t -target] required: (default: target) input target directory containing GenBank-formatted file(s).

[-o -output] required: (default: output) output directory containing TXT-format file(s) with statistic result.

***Example***

1. Input -> Amborella_trichopoda.gb

```
LOCUS       Amborella_trichopoda      162686 bp    DNA     circular UNA 08-JUN-2015
DEFINITION  Amborella trichopoda chloroplast genomic DNA, complete sequence.
ACCESSION   AJ506156
VERSION     AJ506156.2  GI:34481608
KEYWORDS    complete genome.
SOURCE      chloroplast Amborella trichopoda
  ORGANISM  Amborella trichopoda
            Eukaryota; Viridiplantae; Streptophyta; Embryophyta; Tracheophyta;
            Spermatophyta; Magnoliophyta; basal Magnoliophyta; Amborellales;
            Amborellaceae; Amborella.
FEATURES             Location/Qualifiers
     source          1..162686
                    /organism="Amborella trichopoda"
                    /mol_type="genomic DNA"
     repeat_region    90951..117611
                    /note="inverted repeat region B; IRB repeat region"
                    /rpt_type="inverted"
     rRNA           complement(139284..142097)
                    /gene="rrn23"
                    /product="23S ribosomal RNA"
     gene           complement(139284..142097)
                    /gene="rrn23"
     tRNA           join(complement(4472..4508), complement(1840..1874))
                    /gene="trnK-UUU"
                    /product="tRNA-Lys"
     gene           complement(1840..4508)
                    /gene="trnK-UUU"
     CDS            join(complement(16186..16330), complement(14506..14915))
                    /gene="atpF"
                    /codon_start=1
                    /transl_table=11
                    /product="ATPase I subunit"
                    /translation="MKNVTDSFVSLGHWPSAGSFGFNTDIFATNPINLSVVLGVLIFF
                    GKGVLSDLLDNRKQRILSTIRNSEELRGGAIEQLEKARARLRKVEIEADEFRVNGYSE
                    IEREKSNLINAAYENLERLENYKNESIHFEQQRAMNQVRQRVFQQALQGALETLNSYL
                    NSELHLRTISANIGMLGTMKNITD"
     gene           complement(14506..16330)
                    /gene="atpF"
```

2. Input -> Rosa_roxburghii.gb, Rosa_chinensis.gb, Rosa_rugosa.gb

Rosa_roxburghii.gb

```
LOCUS       Rosa_roxburghii  156749 bp    DNA     circular PLN 02-DEC-2022
FEATURES             Location/Qualifiers
     source          1..156749
                     /organism="Rosa_roxburghii"
                     /organelle="plastid:chloroplast"
                     /mol_type="genomic DNA"
                     /note="Annotation Method :: PGA-Plastid Genome Annotator"
     repeat_region   85853..111905
                     /note="inverted repeat B"
                     /rpt_type="inverted"
     repeat_region   complement(130697..156749)
                     /note="inverted repeat A"
                     /rpt_type="inverted"
     gene            58781..60244
                     /gene="accD"
     CDS             58781..60244
                     /gene="accD"
                     /codon_start=1
                     /transl_table=11
                     /product="acetyl-CoA carboxylase carboxyltransferase beta subunit"
     gene            complement(10635..12158)
                     /gene="atpA"
     CDS             complement(10635..12158)
                     /gene="atpA"
                     /codon_start=1
                     /transl_table=11
                     /product="ATP synthase CF1 alpha subunit"
```

3. Output -> Amborella_trichopoda_Rosa_roxburghii.txt, Amborella_trichopoda_Rosa_chinensis.txt, Amborella_trichopoda_Rosa_rugosa.txt

Amborella_trichopoda_Rosa_roxburghii.txt

| type | gene_name | reference_length | target_length | length_difference |
|---|---|---|---|---|
| PCGs | ycf1 | 5385 | 5727 | 342 |
| PCGs | ycf4 | 708 | 555 | -153 |
| PCGs | accD | 1599 | 1464 | -135 |
| PCGs | ndhK | 762 | 681 | -81 |
| PCGs | ycf2 | 6915 | 6840 | -75 |
| PCGs | ycf15 | 147 | 90 | -57 |
| PCGs | rpl22 | 375 | 420 | 45 |
| PCGs | rpoC2 | 4110 | 4155 | 45 |
| PCGs | ndhI | 543 | 504 | -39 |
| PCGs | rps16 | 237 | 270 | 33 |
| PCGs | atpB | 1503 | 1479 | -24 |
| PCGs | ccsA | 942 | 966 | 24 |
| PCGs | psbJ | 123 | 144 | 21 |
| PCGs | rpoC1 | 2043 | 2064 | 21 |
| PCGs | clpP | 609 | 591 | -18 |
| PCGs | rpl32 | 174 | 159 | -15 |
| PCGs | psbA | 1053 | 1062 | 9 |
| PCGs | rpoA | 1005 | 996 | -9 |
| PCGs | rps15 | 264 | 273 | 9 |
| PCGs | matK | 1506 | 1512 | 6 |
| PCGs | psaJ | 129 | 135 | 6 |
| PCGs | psbT | 108 | 102 | -6 |
| PCGs | rpl20 | 360 | 354 | -6 |
| PCGs | rpl23 | 288 | 282 | -6 |
| PCGs | rpl33 | 207 | 201 | -6 |
| PCGs | rpoB | 3219 | 3213 | -6 |
| PCGs | rps8 | 399 | 405 | 6 |
| PCGs | atpE | 405 | 402 | -3 |

#### 3. Submission

***Functionality***

We re-scripted the gbf2tbl.pl Perl tool (provided by NCBI) that can batchly convert multiple GenBank flatfiles (.gb/.gbf/.gbk) to the 5-column feature table files (.tbl) and the sequence files in FASTA format (.fsa/.fasta/.fa/.fas) with required information for submitting to GenBank.

***Features***

1. Multiple GenBank files can be batchly converted to their corresponding 5-column feature table files and the sequence files in FASTA format.

2. Only essential information are retained in the 5-column feature table files. For protein-coding genes, only "gene" is retained in the $4^{th}$ column for "gene" in the $3^{rd}$ column, and "gene/codon_start/transl_table/product" are retained in the $4^{th}$ column for "CDS" in the $3^{rd}$ column; for tRNA genes, only "gene" is retained in the $4^{th}$ column for "tRNA" in the $3^{rd}$ column, and "gene/product" are retained in the $4^{th}$ column for "tRNA" in the $3^{rd}$ column; for rRNA genes, only "gene" is retained in the $4^{th}$ column for "gene" in the $3^{rd}$ column, and "gene/product" are retained in the $4^{th}$ column for "rRNA" in the $3^{rd}$ column.

***How to generate standardized GenBank annotation flatfiles for submission and downstream analysis?***

The storage form of annotation information is important for plastome annotation. The GenBank flatfile is a common template file for storing annotation information of genome sequences, and it can be easily recognized and manipulated using a variety of softwares. If the GenBank flatfile for plastome annotations is standardized, operations such as submission to GenBank and extraction of genes and relevant sequences will become much easier. However, errors are often present in GenBank flatfiles, even when GenBank flatfiles following the RefSeq standards are downloaded directly from GenBank. Therefore, we provide instructions on generating standardized GenBank flatfiles for submission and downstream analysis.

First, plastome sequences should be submitted to GenBank. Depositing annotated sequences to GenBank is an essential step before publication of research results, but this operation is generally inefficient, time-consuming, and tedious. Submission instructions and submission preparation tools (https://www.ncbi.nlm.nih.gov/genbank/submit/) are well documented by NCBI, which can assist with the submission of any type of sequence. For submission of plastome sequences with annotation information to GenBank, we suggest to using Banklt (https://www.ncbi.nlm.nih.gov/WebSub/), a web-based submission tool allowing automatic submission to GenBank. It is easy to fill out the submission template information including locus, definition, accession, organism, reference authors, reference title, and reference journal, all found in the head annotation section of the GenBank flatfiles (.gb/.gbf/.gbk). The GenBank flatfiles can be downloaded from GenBank, but these files are not allowed to be directly submitted to GenBank. So two files need to be prepared separately, including the sequence file in FASTA format (.fsa/.fasta/.fa/.fas) and the 5-column feature table file (.tbl). These two files can be generated by transforming the GenBank flatfiles using the gbf2tbl.pl Perl script provided by NCBI. As the input file of gbf2tbl tool, the incomplete GenBank flatfiles without the head annotation section (e.g., LOCUS, DEFINITION, ACCESSION, and ORGANISM.) are allowed, because this information is not required to generate the .fsa and the .tbl files. Furthermore, when using Banklt, you can generate the head annotation section of the GenBank flatfiles by filling in the corresponding location. That is to say, if you just want to submit the plastome sequences to GenBank, the GenBank flatfiles generated by plastome annotation tools can be incomplete and only retain the annotation and sequence content below the line “FEATURES Location/Qualifiers”. It is also fine if the start-line beginning with the word “LOCUS” is retained.

Second, GenBank flatfiles should be generated for downstream analysis. The head annotation section in the GenBank flatfiles is not necessary when you submit plastome sequences to GenBank, but this content provide some basic statistics useful for extracting information from GenBank flatfiles. The standard GenBank flatfiles of plastome sequences downloaded from GenBank include the head annotation section, which is generated by GenBank based on the information supplied during the Banklt submission process. Currently, some plastome annotation tools like GeSeq and CPGAVAS2 can generate complete GenBank flatfiles and allow users to modify the information included in the head annotation section. However, it is not easy to accurately modify the GenBank flatfiles for those who are not bioinformatics experts. Therefore, it is best to automatically generate head annotation sections that do not require major modifications. This online tool (https://submit.ncbi.nlm.nih.gov/genbank/template/submission/) can be used to prepare the submission template by yourself, and then the generated submission template file (.sbt) can be downloaded to personal computers. Standard GenBank flatfiles can then be generated by adding the content in the .sbt submission template file. When using table2asn (https://www.ncbi.nlm.nih.gov/genbank/table2asn/; table2asn is the replacement of the older now-obsolete tool tbl2asn), three types of files, including the .sbt submission template file, the .tbl feature table file, and the .fsa sequence file, are required to generate standard GenBank flatfiles. It should be noted that, after this transformation, amino acid sequences of each PCG that may not have existed in the initial GenBank flatfiles will then be present following the qualifier “/translation”.

***Future Suggestions***

We all know that the .gb GenBank flatfile and the .tbl feature table file are important for the display of plastome annotation information. If the .gb file contains errors, the .tbl file will also contain errors, and vice versa. Thus we propose to development of scripts or tools to check whether these two important files are complete and following established standards, and whether some essential annotation information in these files is incorrect or even missing.

***Command-line Tool***

perl gbf2tbl.pl –i –o

run:
perl gbf2tbl.pl –i input –o output

parameter:
[-i -input]　　required: (default: input) input directory containing .gb file(s).
[-o -output]　required: (default: output) output directory containing .tbl and .fsa files.

***Example***

1. Input -> Rosa_roxburghii.gb

```
LOCUS       Rosa_roxburghii  156749 bp    DNA     circular PLN 02-DEC-2022
FEATURES             Location/Qualifiers
     source          1..156749
                     /organism="Rosa_roxburghii"
                     /organelle="plastid:chloroplast"
                     /mol_type="genomic DNA"
                     /note="Annotation Method :: PGA-Plastid Genome Annotator"
     repeat_region   85853..111905
                     /note="inverted repeat B"
                     /rpt_type="inverted"
     repeat_region   complement(130697..156749)
                     /note="inverted repeat A"
                     /rpt_type="inverted"
     gene            58781..60244
                     /gene="accD"
     CDS             58781..60244
                     /gene="accD"
                     /codon_start=1
                     /transl_table=11
                     /product="acetyl-CoA carboxylase carboxyltransferase beta subunit"
     gene            complement(10635..12158)
                     /gene="atpA"
     CDS             complement(10635..12158)
                     /gene="atpA"
                     /codon_start=1
                     /transl_table=11
                     /product="ATP synthase CF1 alpha subunit"
```

2. Output -> Rosa_roxburghii.tbl

```
>Feature Rosa_roxburghii
85853	111905	repeat_region
			note	inverted repeat B
			rpt_type	inverted
156749	130697	repeat_region
			note	inverted repeat A
			rpt_type	inverted
58781	60244	gene
			gene	accD
58781	60244	CDS
			gene	accD
			codon_start	1
			transl_table	11
			product	acetyl-CoA carboxylase carboxyltransferase beta subunit
12158	10635	gene
			gene	atpA
12158	10635	CDS
			gene	atpA
			codon_start	1
			transl_table	11
			product	ATP synthase CF1 alpha subunit
55911	54433	gene
			gene	atpB
55911	54433	CDS
			gene	atpB
			codon_start	1
			transl_table	11
			product	ATP synthase CF1 beta subunit
```

3. Output -> Rosa_roxburghii.fsa

```
>Rosa_roxburghii [organism=Rosa_roxburghii] [organelle=plastid:chloroplast]
[mol_type=genomic DNA] [note=Annotation Method :: PGA-Plastid Genome Annotator]
[topology=circular] [gcode=11]
ATGGGCGAACGACGGGAATTGAACCCGCGCGTGGTGGATTCACAATCCACTGCCTTGATC
CACTTGGCTACATCCGCCCCTTGTACTATTTAAATAAAAATAGAAATTACAAATATTTCC
ACCATTTATCATTACTTGTAAAAGAGAATACAACATAAATGGAAACCCTTTTATTTTAAT
CTTTTAGTAAAAAAAAAAAAAGAGTACTAAAATACTAATTTAATTATAGTAATAATTTTA
ATTAATAATTATTTTAATTCATAATTAAAGTAATATGGTAAAGGAGCAACACCAAACCTC
TTGATATAACAAGAATTTTAGTATTGCTCCTTTACTTTCAAAACTTTCACGAACTCATAT
ATACTAAGACCAAAGTCTTATCCATTGATAGATGGAACTTCAACAGCAGCTAGGTCTAGA
GGGAAATTATGAGCATTACGTTCATGCATAACTTCCATACCAAGGTTAGCGCGGTTAATA
ATATCAGCCCAAGTATTAATTACACGACCTTGACTATCAACTACAGATTGATTGAAATTA
AATCCATTTAAGTTGAAAGCCATAGTGCTGACACCTAAAGCGGTGAACCAGATACCTACT
ACAGGCCAAGCAGCTAAGAAGAAGTGTAAAGAACGAGAATTGTTGAAACTAGCATATTGG
AAGATCAATCGGCCAAAATAACCATGAGCGGCTACGATATTATAAGTTTCTTCCTCTTGA
CCGAATCTGTAACCTTCATTAGCAGATTCATTTTCTGTGGTTTCCCTGATCAAACTAGAG
GTTACCAAAGAACCGTGCATAGCACTGAATAGGGAGCCGCCGAATACACCAGCCACACCT
AGCATGTGAAATGGGTGCATAAGGATGTTGTGCTCAGCTTGGAATACAATCATGAAGTTG
AAAGTACCAGAGATTCCTAGAGGCATACCGTCAGAAAAGCTTCCTTGACCAATTGGGTAG
ATCAAGAAAACAGCAGTAGCAGCTGCAACAGGAGCTGAATATGCAACAGCAATCCAAGGG
CGCATACCCAGACGGAAACTAAGTTCCCACTCACGACCCATGTAGCAAGCTACACCAAGT
AAGAAGTGTAGAACAATTAGCTCATAAGGACCACCGTTGTATAACCACTCATCAACGGAA
```

4. Output -> Rosa_roxburghii.err

If there are errors or warnings in the annotation, this file will be generated.

**4. Visualization**

***Functionality***

This tool can batchly generate the circular plastome maps with genes, GC content, quadripartite structure, and dispersed repeats.

***Features***

1. The circular views of plastome maps, with genes, GC content, quadripartite structure, and dispersed repeats, can be batchly generated based on the Circos tool. If you use it, please cite following paper.

Krzywinski M, Schein J, Birol I, et al. Circos: an information aesthetic for comparative genomics. Genome Research, 2009, 19(9): 1639–1645.

2. Both SVG and PNG format figures are exported based on the Circos tool.

3. The color type of gene blocks follows the OGDRAW tool, and the color block illustration figure (color_block_illustration.pdf) can be individually downloaded.

4. GC content are calculated with window size of 100 nucleotides and step size of one nucleotide.

5. The longest inverted-repeat (IR) is generated by blast search the complete plastome against itself, and the minimum allowed length (with default value of 1000 bp) can be changed by users.

6. The dispersed repeats are produced by blast search the complete plastome against itself, and the minimum allowed percent identity for blast search of dispersed repeats (with default value of 90 bp), the minimum allowed length for dispersed repeats (with default value of 15 bp), and the evalue for blast search of dispersed repeats (with default value of 10) can be changed by users. The forward and inverted repeats are shown with black and red color lines, respectively.

7. All intermediate documents for generating these circular plastome maps are also provided for downloading.

***Recommended Suggestions***

1. The standard GenBank format flatfiles should contain information of correct quadripartite structure. That is to say, from 0 bp to 156749 bp (for the whole length of Rosa roxburghii plastome), the quadripartite structure must be LSC-IRb-SSC-IRa.

2. Whether the repeat region (i.e., IRb and IRa) of plastome are annotated in the GenBank flatfiles or not does not affect the display of quadripartite structure (i.e., LSC, SSC, IRb, and IRa).

***Command-line tool***

perl visualization.pl –i [–m –s –l –e] –o

run:

perl scripts\visualization.pl –i visualization\input –m 1000 –s 90 –l 15 –e 10 –o visualization\output

parameter:

[-i -input] required: (default: input) input directory name containing GenBank-formatted file(s).

[-m -minir] optional: (default: 1000) minimum allowed length for the longest inverted-repeat (IR).

[-s -similarity] optional: (default: 90) minimum allowed percent identity for blast search of dispersed repeats.

[-l -length] optional: (default: 15) minimum allowed length for dispersed repeats.

[-e -evalue] optional: (default: 10) evalue for blast search of dispersed repeats.

[-o -output] required: (default: output) output directory name.

***Color block illustration***

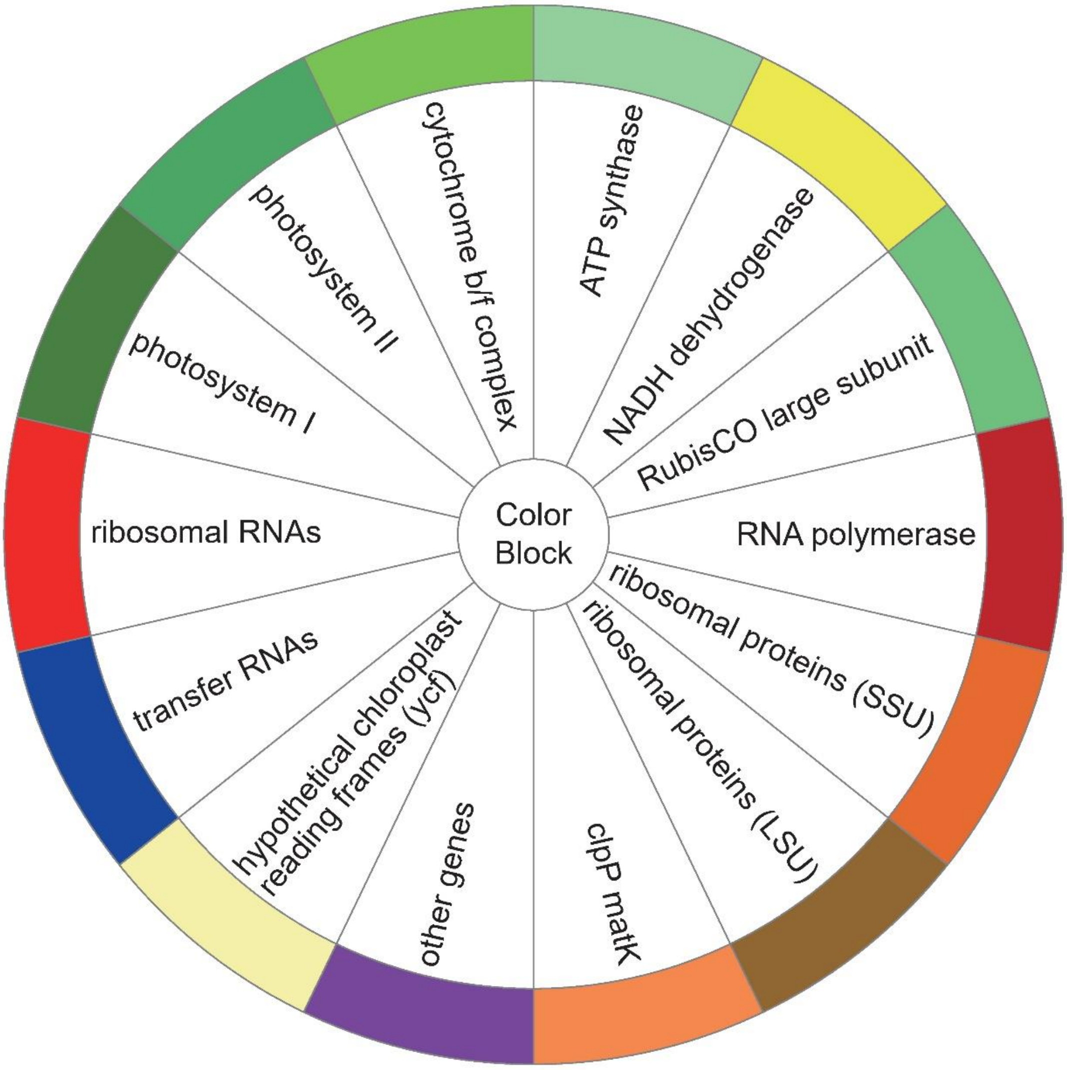

***Example***

Input:

Rosa_roxburghii.gb

Output:

Rosa_roxburghii_GCcontent_background.hist.txt
Rosa_roxburghii_GCcontent_foreground.hist.txt
Rosa_roxburghii_ideogram.conf
Rosa_roxburghii_innergenes.label.txt
Rosa_roxburghii_innergenes.position.txt
Rosa_roxburghii_karyotype.txt
Rosa_roxburghii_outergenes.label.txt
Rosa_roxburghii_outergenes.position.txt
Rosa_roxburghii_quadripartite.highlight.txt
Rosa_roxburghii_quadripartite.IRb.IRa.txt
Rosa_roxburghii_quadripartite.junction.txt
Rosa_roxburghii_quadripartite.label.txt
Rosa_roxburghii_repeatsFR.hist.txt
Rosa_roxburghii_repeatsFR.link.txt
Rosa_roxburghii_repeatsIR.hist.txt
Rosa_roxburghii_repeatsIR.link.txt
Rosa_roxburghii_species_length.txt
Rosa_roxburghii_ticks.conf
Rosa_roxburghii_visualization.conf
Rosa_roxburghii_visualization.png
Rosa_roxburghii_visualization.svg

1. Input -> Rosa_roxburghii.gb

```
LOCUS       Rosa_roxburghii  156749 bp    DNA     circular PLN 02-DEC-2022
FEATURES             Location/Qualifiers
     source          1..156749
                     /organism="Rosa_roxburghii"
                     /organelle="plastid:chloroplast"
                     /mol_type="genomic DNA"
                     /note="Annotation Method :: PGA-Plastid Genome Annotator"
     repeat_region   85853..111905
                     /note="inverted repeat B"
                     /rpt_type="inverted"
     repeat_region   complement(130697..156749)
                     /note="inverted repeat A"
                     /rpt_type="inverted"
     gene            58781..60244
                     /gene="accD"
     CDS             58781..60244
                     /gene="accD"
                     /codon_start=1
                     /transl_table=11
                     /product="acetyl-CoA carboxylase carboxyltransferase beta subunit"
     gene            complement(10635..12158)
                     /gene="atpA"
     CDS             complement(10635..12158)
                     /gene="atpA"
                     /codon_start=1
                     /transl_table=11
                     /product="ATP synthase CF1 alpha subunit"
```

2. Output -> Rosa_roxburghii_visualization.png

3. Output -> Rosa_roxburghii_visualization.svg

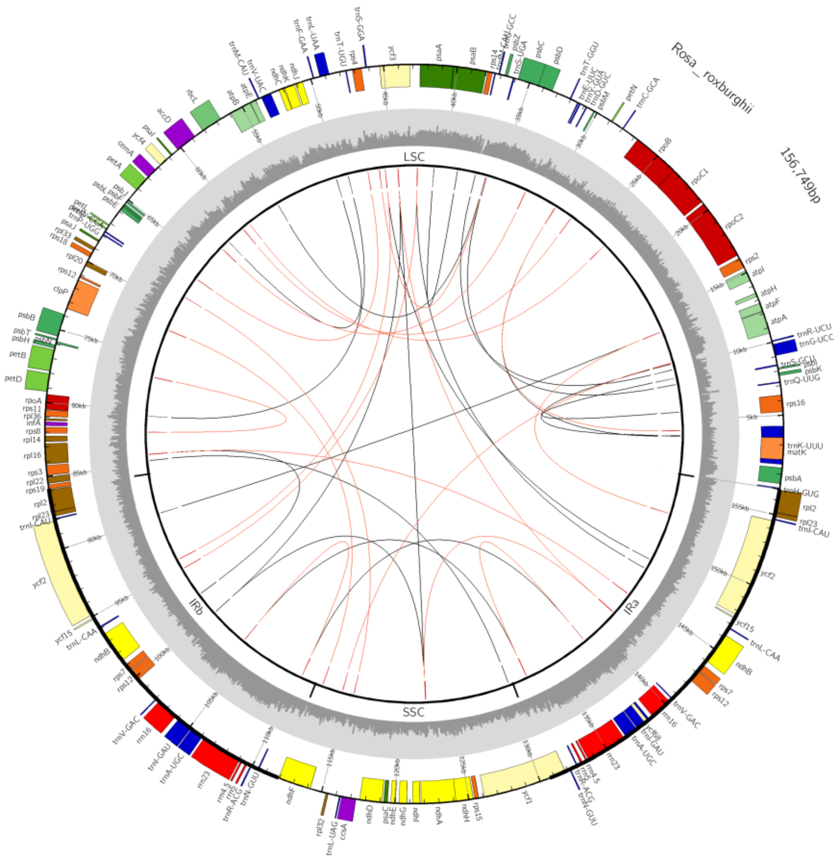

4. Output -> Rosa_roxburghii_GCcontent_background.hist.txt

```
Rosa_roxburghii 1       2       0
Rosa_roxburghii 1       100     1
Rosa_roxburghii 101     200     1
Rosa_roxburghii 201     300     1
Rosa_roxburghii 301     400     1
Rosa_roxburghii 401     500     1
Rosa_roxburghii 501     600     1
Rosa_roxburghii 601     700     1
Rosa_roxburghii 701     800     1
Rosa_roxburghii 801     900     1
Rosa_roxburghii 901     1000    1
```

5. Output -> Rosa_roxburghii_GCcontent_foreground.hist.txt

```
Rosa_roxburghii 1       2       1
Rosa_roxburghii 1       100     0.49
Rosa_roxburghii 101     200     0.2
Rosa_roxburghii 201     300     0.19
Rosa_roxburghii 301     400     0.29
Rosa_roxburghii 401     500     0.4
Rosa_roxburghii 501     600     0.38
Rosa_roxburghii 601     700     0.41
Rosa_roxburghii 701     800     0.39
Rosa_roxburghii 801     900     0.48
Rosa_roxburghii 901     1000    0.45
```

6. Output -> Rosa_roxburghii_ideogram.conf

```
<ideogram>

<spacing>
default = 0r
break   = 0
</spacing>


radius           = 0.8r
thickness        = 2p
fill             = yes

fill_color       = gpos100
stroke_thickness = 3
stroke_color     = black

</ideogram>
```

7. Output -> Rosa_roxburghii_innergenes.label.txt

```
Rosa_roxburghii 4       77      trnH-GUG
Rosa_roxburghii 378     1439    psbA
Rosa_roxburghii 1704    4278    trnK-UUU
Rosa_roxburghii 1998    3509    matK
Rosa_roxburghii 5247    6381    rps16
Rosa_roxburghii 7213    7284    trnQ-UUG
Rosa_roxburghii 8379    8466    trnS-GCU
Rosa_roxburghii 10635   12158   atpA
Rosa_roxburghii 12213   12767   atpF
Rosa_roxburghii 13249   13494   atpH
```

8. Output -> Rosa_roxburghii_innergenes.position.txt

```
Rosa_roxburghii 4       77      fill_color=chr14
Rosa_roxburghii 378     1439    fill_color=dgreen
Rosa_roxburghii 1704    4278    fill_color=chr14
Rosa_roxburghii 1998    3509    fill_color=orange
Rosa_roxburghii 5247    6381    fill_color=dorange
Rosa_roxburghii 7213    7284    fill_color=chr14
Rosa_roxburghii 8379    8466    fill_color=chr14
Rosa_roxburghii 10635   12158   fill_color=lgreen
Rosa_roxburghii 12213   12767   fill_color=lgreen
Rosa_roxburghii 13249   13494   fill_color=lgreen
```

9. Output -> Rosa_roxburghii_karyotype.txt

```
chr - Rosa_roxburghii Rosa_roxburghii 0 156749 black
```

10. Output -> Rosa_roxburghii_outergenes.label.txt

```
Rosa_roxburghii 7704    7889    psbK
Rosa_roxburghii 8126    8236    psbI
Rosa_roxburghii 9069    9834    trnG-UCC
Rosa_roxburghii 10027   10098   trnR-UCU
Rosa_roxburghii 27932   28002   trnC-GCA
Rosa_roxburghii 28813   28902   petN
Rosa_roxburghii 32018   32089   trnT-GGU
Rosa_roxburghii 33399   34460   psbD
Rosa_roxburghii 34408   35829   psbC
Rosa_roxburghii 36549   36737   psbZ
```

11. Output -> Rosa_roxburghii_outergenes.position.txt

```
Rosa_roxburghii 7704    7889    fill_color=dgreen
Rosa_roxburghii 8126    8236    fill_color=dgreen
Rosa_roxburghii 9069    9834    fill_color=chr14
Rosa_roxburghii 10027   10098   fill_color=chr14
Rosa_roxburghii 27932   28002   fill_color=chr14
Rosa_roxburghii 28813   28902   fill_color=chrun
Rosa_roxburghii 32018   32089   fill_color=chr14
Rosa_roxburghii 33399   34460   fill_color=dgreen
Rosa_roxburghii 34408   35829   fill_color=dgreen
Rosa_roxburghii 36549   36737   fill_color=dgreen
```

12. Output -> Rosa_roxburghii_quadripartite.highlight.txt

```
Rosa_roxburghii 1       85852   fill_color=black,z=0
Rosa_roxburghii 85853   111905  fill_color=black,z=0
Rosa_roxburghii 111906  130696  fill_color=black,z=0
Rosa_roxburghii 130697  156749  fill_color=black,z=0
```

13. Output -> Rosa_roxburghii_quadripartite.IRb.IRa.txt

```
Rosa_roxburghii 85853   111905  fill_color=black,z=0
Rosa_roxburghii 130697  156749  fill_color=black,z=0
```

14. Output -> Rosa_roxburghii_quadripartite.junction.txt

```
Rosa_roxburghii 1       2       0
Rosa_roxburghii 85851   85852   10
Rosa_roxburghii 111904  111905  10
Rosa_roxburghii 130695  130696  10
Rosa_roxburghii 156748  156749  10
```

15. Output -> Rosa_roxburghii_quadripartite.label.txt

```
Rosa_roxburghii 1       85852   LSC
Rosa_roxburghii 85853   111905  IRb
Rosa_roxburghii 111906  130696  SSC
Rosa_roxburghii 130697  156749  IRa
```

16. Output -> Rosa_roxburghii_repeatsFR.hist.txt

```
Rosa_roxburghii 1       2       0
Rosa_roxburghii 3504    3523    10
Rosa_roxburghii 29017   29036   10
Rosa_roxburghii 110448  110465  10
Rosa_roxburghii 138495  138512  10
Rosa_roxburghii 38857   38874   10
Rosa_roxburghii 41072   41089   10
Rosa_roxburghii 84591   84611   10
Rosa_roxburghii 130310  130330  10
Rosa_roxburghii 5798    5814    10
```

17. Output -> Rosa_roxburghii_repeatsFR.link.txt

```
Rosa_roxburghii 3504    3523    Rosa_roxburghii 29017   29036
Rosa_roxburghii 110448  110465  Rosa_roxburghii 138495  138512
Rosa_roxburghii 38857   38874   Rosa_roxburghii 41072   41089
Rosa_roxburghii 84591   84611   Rosa_roxburghii 130310  130330
Rosa_roxburghii 5798    5814    Rosa_roxburghii 8991    9007
Rosa_roxburghii 8376    8407    Rosa_roxburghii 36067   36098
Rosa_roxburghii 45657   45673   Rosa_roxburghii 148285  148301
Rosa_roxburghii 3594    3613    Rosa_roxburghii 10355   10374
Rosa_roxburghii 44442   44482   Rosa_roxburghii 122358  122398
Rosa_roxburghii 36172   36194   Rosa_roxburghii 56309   56331
```

18. Output -> Rosa_roxburghii_repeatsIR.hist.txt

```
Rosa_roxburghii 1       2       0
Rosa_roxburghii 84010   84031   10
Rosa_roxburghii 144496  144473  10
Rosa_roxburghii 10431   10450   10
Rosa_roxburghii 114530  114511  10
Rosa_roxburghii 45657   45673   10
Rosa_roxburghii 94317   94301   10
Rosa_roxburghii 132137  132154  10
Rosa_roxburghii 138512  138495  10
Rosa_roxburghii 122359  122398  10
```

19. Output -> Rosa_roxburghii_repeatsIR.link.txt

```
Rosa_roxburghii 84010   84031   Rosa_roxburghii 144496  144473
Rosa_roxburghii 10431   10450   Rosa_roxburghii 114530  114511
Rosa_roxburghii 45657   45673   Rosa_roxburghii 94317   94301
Rosa_roxburghii 132137  132154  Rosa_roxburghii 138512  138495
Rosa_roxburghii 122359  122398  Rosa_roxburghii 142314  142275
Rosa_roxburghii 76780   76796   Rosa_roxburghii 81952   81936
Rosa_roxburghii 44445   44486   Rosa_roxburghii 142312  142271
Rosa_roxburghii 27238   27258   Rosa_roxburghii 60301   60281
Rosa_roxburghii 36071   36098   Rosa_roxburghii 46233   46206
Rosa_roxburghii 104090  104107  Rosa_roxburghii 110465  110448
```

20. Output -> Rosa_roxburghii_species_length.txt

```
Rosa_roxburghii 15994   20899   156,749bp
Rosa_roxburghii 23051   27990   Rosa_roxburghii
```

21. Output -> Rosa_roxburghii_ticks.conf

```
show_ticks          = yes
show_tick_labels    = yes
show_grid           = yes

<ticks>
skip_first_label     = no
skip_last_label      = no
radius               = dims(ideogram,radius_outer)
tick_separation      = 2p
min_label_distance_to_edge = 10p
label_separation = 5p
label_offset     = 5p
multiplier       = 1e-3
color            = black
```

22. Output -> Rosa_roxburghii_visualization.conf

```
<<include etc/colors_fonts_patterns.conf>>
<<include etc/housekeeping.conf>>

<<include Rosa_roxburghii_ideogram.conf>>
<<include Rosa_roxburghii_ticks.conf>>

<image>
<<include etc/image.conf>>
#angle_orientation* = clockwise
angle_orientation* = counterclockwise
#background* = transparent
#radius* = 1500p
angle_offset* = 8.6
</image>

karyotype   = visualization\output/Rosa_roxburghii_karyotype.txt
chromosomes_units           = 1000
chromosomes_display_default = yes
```

#### 5. Extraction

***Functionality***

This tool can batchly extract coding and noncoding sequences from multiple GenBank flatfiles of plastomes based on different extraction patterns.

***Features***

1. Coding and noncoding sequences in plastomes can be batchly extracted from multiple GenBank flatfiles.

2. Different extraction patterns are developed for plastome sequences, not only PCGs with introns and sequences among PCGs are extracted, but also genes with introns and intergenic sequences are extracted, not only coding sequences with linked CDSs and tRNAs, and noncoding sequences are considered, but also coding sequences with unlinked CDSs and tRNAs, and noncoding sequences are considered.

3. Different extraction patterns can be combined, i.e., extraction pattern 5 = extraction pattern 1 + extraction pattern 2, extraction pattern 6 = extraction pattern 3 + extraction pattern 4, and extraction pattern 7 = extraction pattern 1 + extraction pattern 2 + extraction pattern 3 + extraction pattern 4.

***Command-line Tool***

```
perl extraction.pl –i –p –o

run:
perl extraction.pl –i input –p 3 –o output

parameter:
[-i -input]     required: (default: input) input directory containing GenBank-formatted files.
[-p -pattern]   required: (default: 3) 1/2/3/4/5/6/7 extraction pattern,
                1=PCGs with introns, and sequences among PCGs,
                2=genes with introns, and intergenic sequences,
                3=coding sequences with linked CDSs and tRNAs, and noncoding sequences,
                4=coding sequences with unlinked CDSs and tRNAs, and noncoding sequences,
                5=1+2,
                6=3+4,
                7=1+2+3+4.
[-o -output]    required: (default: output) output directory.
```

***Principles***

The gene types in plastomes and four extraction patterns are shown in Figures 1, 2, and 3.

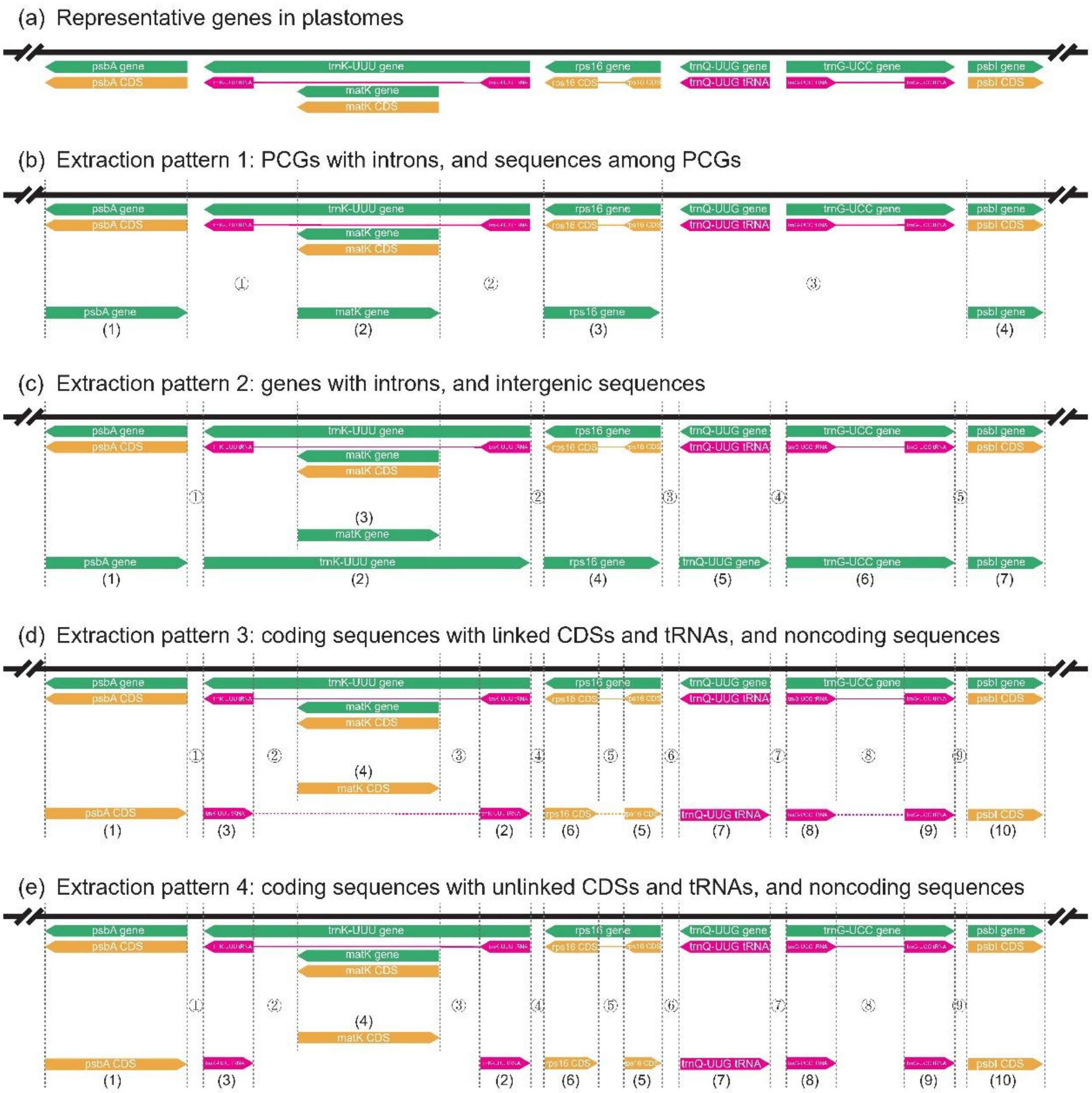


Figure 1. The canonical gene types such as protein-coding genes and tRNA genes without and with introns in plastomes and four extraction patterns are shown.

(a) Representative gene types of plastome. Canonical gene types are shown.

(b) Extraction pattern 1: PCGs with introns, and sequences among PCGs. The labels “(1), (2), (3), (4)” indicate PCGs. The labels “①, ②, ③” indicate sequences among PCGs, including intergenic sequences, tRNAs or rRNAs within these sequences.

(c) Extraction pattern 2: genes with introns, and intergenic sequences. The labels “(1), (2), (3), (4), (5), (6), (7)” indicate genes. The labels “①, ②, ③, ④, ⑤” indicate intergenic sequences.

(d) Extraction pattern 3: coding sequences with linked CDSs and tRNAs, and noncoding sequences. The labels “(1), (2), (3), (4), (5), (6), (7), (8), (9), (10)” indicate coding sequences. The labels “①, ②, ③, ④, ⑤, ⑥, ⑦, ⑧, ⑨” indicate noncoding sequences, including intergenic sequences and introns. The exons of the PCGs and tRNAs with introns are connected with dash line. For examples, exon1 “(2)” and exon2 “(3)”of *trnK-UUU* are connected with dash line, exon1 “(5)” and exon2 “(6)” of rps16 are connected with dash line, exon1 “(8)” and exon2 “(9)” of *trnG-UCC* are connected with dash line.

(e) Extraction pattern 4: coding sequences with unlinked CDSs and tRNAs, and noncoding sequences. The labels “(1), (2), (3), (4), (5), (6), (7), (8), (9), (10)” indicate coding sequences. The labels "①, ②, ③, ④, ⑤, ⑥, ⑦, ⑧, ⑨" indicate noncoding sequences, including intergenic sequences and introns. The exons of the PCGs and tRNAs with introns are separately presented. For examples, exon1 "(2)" and exon2 "(3)"of *trnK-UUU* are separately presented, exon1 "(5)" and exon2 "(6)" of rps16 are separately presented, exon1 "(8)" and exon2 "(9)" of *trnG-UCC* are separately presented.

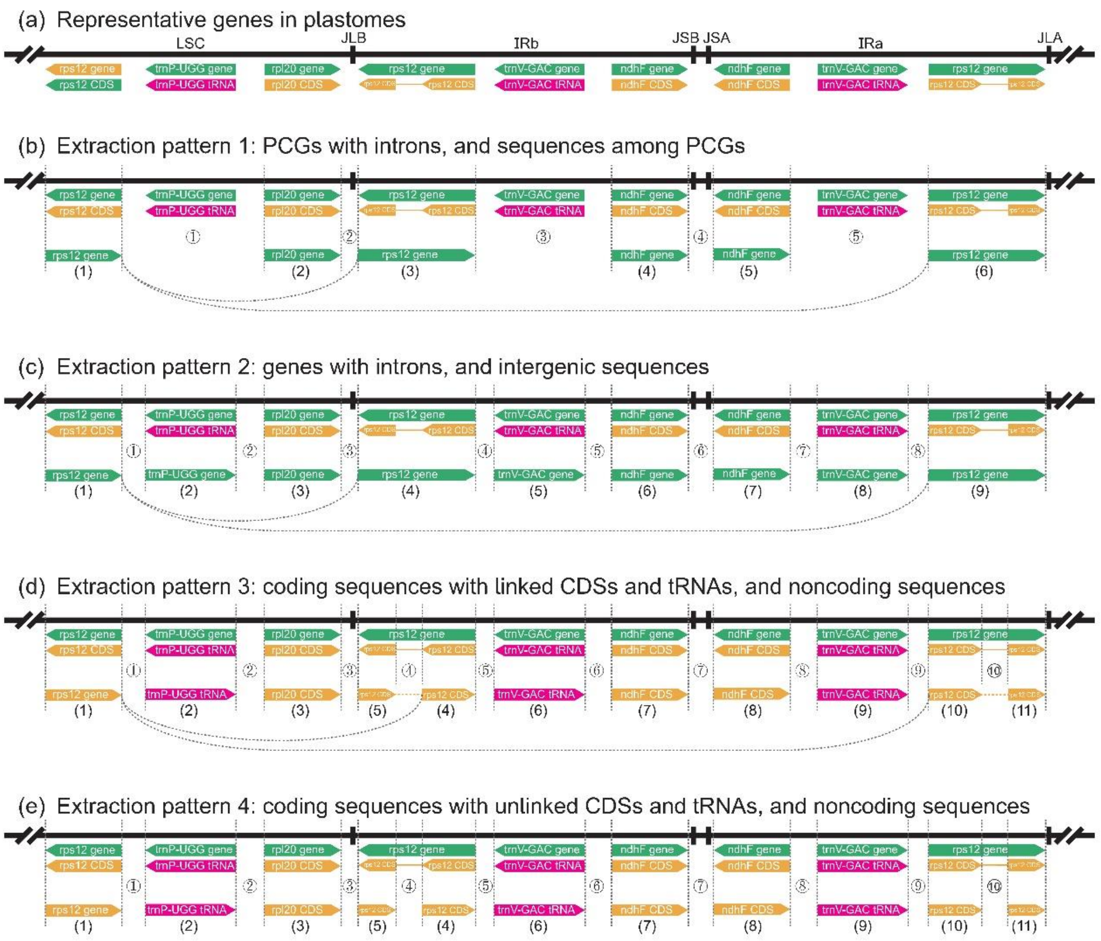


Figure 2. The special gene type like trans-splicing gene *rps12* in plastomes and four extraction patterns are shown.

(a) Representative gene types of plastome. The trans-splicing gene *rps12* are shown.

(b) Extraction pattern 1: PCGs with introns, and sequences among PCGs. The labels "(1), (2), (3), (4), (5), (6)" indicate PCGs. The labels "①, ②, ③, ④, ⑤" indicate sequences among PCGs, including intergenic sequences, tRNAs or rRNAs within these sequences. The exon1 of *rps12* in the LSC region "(1)" and the exon2, exon3 and intron of *rps12* in the IRb region "(3)" are connected with curve line, and the exon1 of *rps12* in the LSC region "(1)" and the exon2, exon3 and intron of *rps12* in the IRa region "(6)" are also connected with curve line.

(c) Extraction pattern 2: genes with introns, and intergenic sequences. The labels "(1), (2), (3), (4), (5), (6), (7), (8), (9)" indicate genes. The labels "①, ②, ③, ④, ⑤, ⑥, ⑦, ⑧" indicate intergenic sequences. The exon1 of *rps12* in the LSC region "(1)" and the exon2, exon3 and intron of *rps12* in the IRb region "(4)" are connected with curve line, and the exon1 of *rps12* in the LSC region "(1)" and the exon2, exon3 and intron of *rps12* in the IRa region "(9)" are also connected with curve line.

(d) Extraction pattern 3: coding sequences with linked CDSs and tRNAs, and noncoding sequences. The labels "(1), (2), (3), (4), (5), (6), (7), (8), (9), (10), (11)" indicate coding sequences. The labels "①, ②, ③, ④, ⑤, ⑥, ⑦, ⑧, ⑨, ⑩" indicate noncoding sequences, including intergenic sequences and introns. The exon1 of *rps12* in the LSC region "(1)" and the exon2 "(4)" and exon3 "(5)" of *rps12* in the IRb region are connected with curve line, and the exon1 of *rps12* in the LSC region "(1)" and the exon2 "(10)" and exon3 "(11)" of *rps12* in the IRa region are also connected with curve line.

(e) Extraction pattern 4: coding sequences with unlinked CDSs and tRNAs, and noncoding sequences. The labels "(1), (2), (3), (4), (5), (6), (7), (8), (9), (10), (11)" indicate coding sequences. The labels "①, ②, ③, ④, ⑤, ⑥, ⑦, ⑧, ⑨, ⑩" indicate noncoding sequences, including intergenic sequences and introns. The exon1 of *rps12* in the LSC region "(1)" and the exon2 "(4)" and exon3 "(5)" of *rps12* in the IRb region are separately presented, and the exon1 of *rps12* in the LSC region "(1)" and the exon2 "(10)" and exon3 "(11)" of *rps12* in the IRa region are also separately presented.

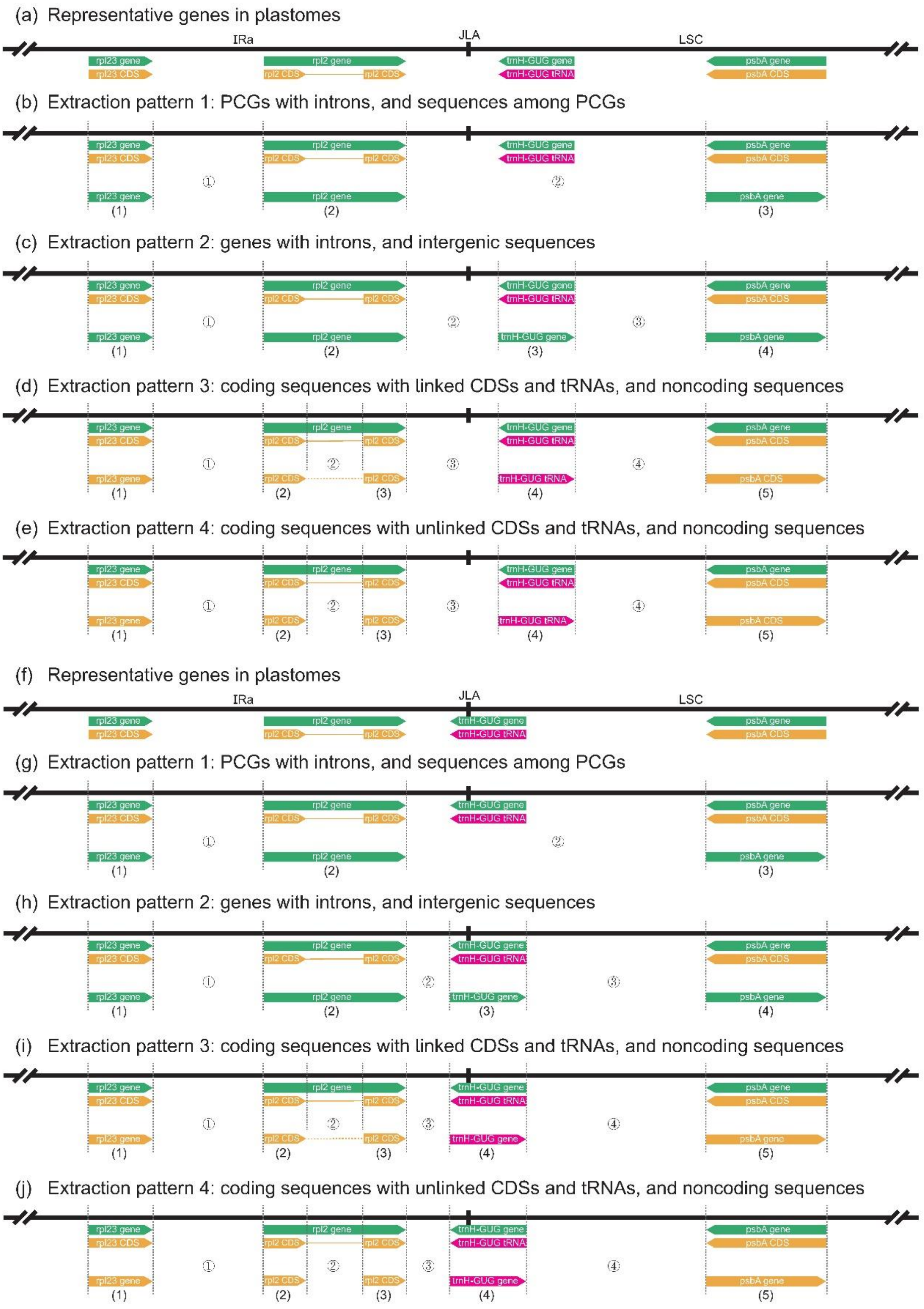


Figure 3. The gene that does not span or spans the start-end position (JLA junction of IRa-LSC) in circular plastomes and four extraction patterns are shown.

(a) Representative gene types of plastome. The tRNA gene *trnH-GUG* does not span the start-end position (JLA junction of IRa-LSC).

(b) Extraction pattern 1: PCGs with introns, and sequences among PCGs. The labels “(1), (2), (3)” indicate PCGs. The labels “①, ②” indicate sequences among PCGs, including intergenic sequences, tRNAs or rRNAs within these sequences. For the sequences among PCGs that span the start-end position (JLA junction of IRa-LSC), i.e. "②", the extracted sequence is *rpl2-psbA*, which indicates that the sequence *rpl2*-end (or *rpl2*-JLA) and the sequence start-*psbA* (or JLA-*psbA*) are connected.

(c) Extraction pattern 2: genes with introns, and intergenic sequences. The labels "(1), (2), (3), (4)" indicate genes. The labels "①, ②, ③" indicate intergenic sequences. For the intergenic sequences that span the start-end position (JLA junction of IRa-LSC), i.e. "②", the extracted sequence is *rpl2-trnH-GUG*, which indicates that the sequence *rpl2*-end (or *rpl2*-JLA) and the sequence start-*trnH-GUG* (or JLA-*trnH-GUG*) are connected.

(d) Extraction pattern 3: coding sequences with linked CDSs and tRNAs, and noncoding sequences. The labels "(1), (2), (3), (4), (5)" indicate coding sequences. The labels "①, ②, ③, ④" indicate noncoding sequences, including intergenic sequences and introns. The exons of the PCGs and tRNAs with introns are connected with dash line. For examples, exon1 "(2)" and exon2 "(3)"of *rpl2* are connected with dash line. For the noncoding sequences that span the start-end position (JLA junction of IRa-LSC), i.e. "③", the extracted sequence is *rpl2*-2-*trnH-GUG*, which indicates that the sequence *rpl2*-2-end (or *rpl2*-2-JLA) and the sequence start-*trnH-GUG* (or JLA-*trnH-GUG*) are connected.

(e) Extraction pattern 4: coding sequences with unlinked CDSs and tRNAs, and noncoding sequences. The labels "(1), (2), (3), (4), (5)" indicate coding sequences. The labels "①, ②, ③, ④" indicate noncoding sequences, including intergenic sequences and introns. The exons of the PCGs and tRNAs with introns are separately presented. For examples, exon1 "(2)" and exon2 "(3)"of *rpl2* are separately presented. For the noncoding sequences that span the start-end position (JLA junction of IRa-LSC), i.e. "③", the extracted sequence is *rpl2*-2-*trnH-GUG*, which indicates that the sequence *rpl2*-2-end (or *rpl2*-2-JLA) and the sequence start-*trnH-GUG* (or JLA-*trnH-GUG*) are connected.

(f) Representative gene types of plastome. The tRNA gene *trnH-GUG* spans the start-end position (JLA junction of IRa-LSC).

(g) Extraction pattern 1: PCGs with introns, and sequences among PCGs. The labels "(1), (2), (3)" indicate PCGs. The labels "①, ②" indicate sequences among PCGs, including intergenic sequences, tRNAs or rRNAs within these sequences. For the sequences among PCGs that span the start-end position (JLA junction of IRa-LSC), i.e. "②", the extracted sequence is *rpl2-psbA*, which indicates that the sequence *rpl2*-end (or *rpl2*-JLA) and the sequence start-*psbA* (or JLA-*psbA*) are connected.

(h) Extraction pattern 2: genes with introns, and intergenic sequences. The labels "(1), (2), (3), (4)" indicate genes. The labels "①, ②, ③" indicate intergenic sequences. For the gene that span the start-end position (JLA junction of IRa-LSC), i.e. "(3)", the extracted sequence is *trnH-GUG*, which indicates that the 5'-end sequence of *trnH-GUG* (or 5'-end-*trnH-GUG*-JLA) and the 3'-end sequence of *trnH-GUG* (or JLA-3'-end-*trnH-GUG*) are connected.

(i) Extraction pattern 3: coding sequences with linked CDSs and tRNAs, and noncoding sequences. The labels "(1), (2), (3), (4), (5)" indicate coding sequences. The labels "①, ②, ③, ④" indicate noncoding sequences, including intergenic sequences and introns. The exons of the PCGs and tRNAs with introns are connected with dash line. For examples, exon1 "(2)" and exon2 "(3)"of *rpl2* are connected with dash line. For the coding sequences that span the start-end position (JLA junction of IRa-LSC), i.e. "(4)", the extracted sequence is *trnH-GUG*, which indicates that the 5'-end sequence of *trnH-GUG* (or 5’-end-*trnH-GUG*-JLA) and the 3’-end sequence of *trnH-GUG* (or JLA-3’-end-*trnH-GUG*) are connected.

(j) Extraction pattern 4: coding sequences with unlinked CDSs and tRNAs, and noncoding sequences. The labels “(1), (2), (3), (4), (5)” indicate coding sequences. The labels “①, ②, ③, ④” indicate noncoding sequences, including intergenic sequences and introns. The exons of the PCGs and tRNAs with introns are separately presented. For examples, exon1 “(2)” and exon2 “(3)”of *rpl2* are separately presented. For the coding sequences that span the start-end position (JLA junction of IRa-LSC), i.e. “(4)”, the extracted sequence is *trnH-GUG*, which indicates that the 5’-end sequence of *trnH-GUG* (or 5’-end-*trnH-GUG*-JLA) and the 3’-end sequence of *trnH-GUG* (or JLA-3’-end-*trnH-GUG*) are connected.

***Example***

1. Input -> Rosa_roxburghii.gb

```
LOCUS       Rosa_roxburghii  156749 bp    DNA     circular PLN 02-DEC-2022
FEATURES             Location/Qualifiers
     source          1..156749
                     /organism="Rosa_roxburghii"
                     /organelle="plastid:chloroplast"
                     /mol_type="genomic DNA"
                     /note="Annotation Method :: PGA-Plastid Genome Annotator"
     repeat_region   85853..111905
                     /note="inverted repeat B"
                     /rpt_type="inverted"
     repeat_region   complement(130697..156749)
                     /note="inverted repeat A"
                     /rpt_type="inverted"
     gene            58781..60244
                     /gene="accD"
     CDS             58781..60244
                     /gene="accD"
                     /codon_start=1
                     /transl_table=11
                     /product="acetyl-CoA carboxylase carboxyltransferase beta subunit"
     gene            complement(10635..12158)
                     /gene="atpA"
     CDS             complement(10635..12158)
                     /gene="atpA"
                     /codon_start=1
                     /transl_table=11
                     /product="ATP synthase CF1 alpha subunit"
```

2. Output -> Rosa_roxburghii_p3_linked_CDS_RNA.bed

```
Rosa_roxburghii trnH-GUG       -       4       77      tRNA
Rosa_roxburghii psbA    -       378     1439    CDS
Rosa_roxburghii trnK-UUU-2     -       1704    1738    tRNA
Rosa_roxburghii matK    -       1998    3509    CDS
Rosa_roxburghii trnK-UUU-1     -       4242    4278    tRNA
Rosa_roxburghii rps16-2 -       5247    5476    CDS
Rosa_roxburghii rps16-1 -       6342    6381    CDS
Rosa_roxburghii trnQ-UUG       -       7213    7284    tRNA
Rosa_roxburghii psbK    +       7704    7889    CDS
Rosa_roxburghii psbI    +       8126    8236    CDS
Rosa_roxburghii trnS-GCU       -       8379    8466    tRNA
Rosa_roxburghii trnG-UCC-1     +       9069    9091    tRNA
Rosa_roxburghii trnG-UCC-2     +       9787    9834    tRNA
```

3. Output -> Rosa_roxburghii_p3_linked_CDS_RNA.fasta

```
>trnH-GUG_Rosa_roxburghii
gcggatgtagccaagtggatcaaggcagtggattgtgaatccaccacgcgcgggttcaattcccgtcgttcgcc
>psbA_Rosa_roxburghii
atgactgcaattttagagagacgtgaaagcgaaagcctatggggtcgcttttgtaattggataaccagcactgaaaaccgtctttacattggatggtttg
>trnK-UUU_Rosa_roxburghii
gggttgctaactcaacggtagagtactcggcttttaaccgactagttccgggttcgagtcccgggcaaccca
>matK_Rosa_roxburghii
atggaagaatttcaaggatatttagaattatatagatctcagcaacatgacttcctatacccacttatctttcgggagtatatttatgcacttgctcatg
>rps16_Rosa_roxburghii
atggtaaaacttcgtttaaaacgatgtggtagaaagcaacgagccatttatcgaatcgttgcaattgatgttcgatcccgacgagagggaagagatcttc
>trnQ-UUG_Rosa_roxburghii
tggggcgtggccaagtggtaaggcaacgggttttggtcccgctactcggaggttcgaatccttccgtcccag
>psbK_Rosa_roxburghii
atgcttaatattttaagtttgatctgtatctaccttaattctgctctttattccagtagtttttcgtcgccaaattgcccgaagcctacgctttttga
>psbI_Rosa_roxburghii
atgcttactctcaaactatttgtttacacggtagtgatattctttgtttctctctttatcttcggattcctgtctaatgatccaggacgtaatcctggac
>trnS-GCU_Rosa_roxburghii
ggagagatggctgagtggactaaagcgtcggattgctaatccgttgtacgagtttttcgtaccgagggttcgaatccctctctttccg
>trnG-UCC_Rosa_roxburghii
gcgggtatagtttagtggtaaaaccctagccttccaagctaacgatgcgggttcgattcccgctacccgct
```

4. Output -> Rosa_roxburghii_p3_regions_among_linked_CDS_RNA.fasta

```
>trnH-GUG-psbA_Rosa_roxburghii
cccttgtactatttaaataaaaatagaaattacaaatatttccaccatttatcattacttgtaaaagagaatacaacataaatggaaacccttttatttt
>psbA-trnK-UUU-2_Rosa_roxburghii
ggtaaaatcttggtttatttaatcattaatcatcagggactcccaagcacacaaattctctataaatagataaatagaaataatagataattgagggctt
>trnK-UUU-2-matK_Rosa_roxburghii
atggagtagataatttccttgttagaaaaaaaataggtaaaaaaaccctccccaaaccgtgcttgcatttttcattgcacacggctttccctatggatac
>matK-trnK-UUU-1_Rosa_roxburghii
ttaaaattcaatttgaatcagaaaagaaataggtgatttattgggttatcaaatgatacatagtacgatacagtcaaaacaaggtatttatattattatt
>trnK-UUU-1-rps16-2_Rosa_roxburghii
aaataaacaaattagaatgtgtagatacaatcagaatcccaataaataacgaaattgaatggtacaacacaatcaaaccattaaaaaaaaaaaaaaaatg
>rps16-2-rps16-1_Rosa_roxburghii
attgggatagatgaagaataccccccctagaaacgtataagaagttttcccctcgtacggctcgagaaaattcgaaattatgtctatgtatagaattaca
>rps16-1-trnQ-UUG_Rosa_roxburghii
aacattccttcagtttggaacccatatgtaattgattcaattatggaatcatgaataatcattggttcggatagagattaatagaatttaatattggaaa
>trnQ-UUG-psbK_Rosa_roxburghii
tttatttctattcgacactaataaacactaatattggtacgggttattcgtcaactccagtctaaatatctattatttctaaaatggattactagaattt
>psbK-psbI_Rosa_roxburghii
tatttttaataaagtcccagacaaattcatgatttattcgaaaaaaattcgtggaattgataagatcagatacgtcttacactctcaattcaaatattga
>psbI-trnS-GCU_Rosa_roxburghii
aaaataaggttttctttgcttgattttaaactgttattagtaagattttatctattccacttctttaactattaatttttaactattaataaaaaagagc
```

#### 6. Pre-Alignment

This tool kit contains eight tools that can complete pre-alignment processing for the sequence matrices, such as "6.1. Sort Genenames, 6.2. Remove Duplications, 6.3. Combine Fasta, 6.4. Generate Matrices, 6.5. Check Codons, 6.6. Check Missings, 6.7. Extract Sites, and 6.8. Codon-to-AA".

##### 6.1. Sort Genenames

***Functionality***

This tool can batchly sort flanking genenames and reverse complement their corresponding sequences for the intergenic regions in plastomes. Because some plastomes have obvious structural rearrangements such as inversions and translocations, it is necessary to adjust their sequences to the same direction before aligning intergenic regions for phylogenetic reconstruction. This processing will improve the alignment accuracy of intergenic regions in plastomes.

***Features***

1. Flanking genenames and their corresponding sequences of the intergenic regions can be batchly sorted for different sequence files.
2. The flanking genenames in the header of fasta format files will be sorted according to ASCII order.
3. The sequences with sorted flanking genenames will be reverse complemented.

***Command-line Tool***

perl sort_genenames.pl –i –o

run:
perl sort_genenames.pl –i input –o output

parameter:
[-i -input]  required: (default: input) input directory containing fasta files.
[-o -output]  required: (default: output) output directory.

***Example***

1. Input -> Amaranthus_caudatus_intergenic.fasta, Amaranthus_tricolor_intergenic.fasta

| | | | |
|---|---|---|---|
| Amaranthus_caudatus_intergenic.fasta | 2022/12/11 11:45 | FASTA 文件 | 66 KB |
| Amaranthus_tricolor_intergenic.fasta | 2022/12/11 11:45 | FASTA 文件 | 74 KB |

Amaranthus_caudatus_intergenic.fasta

```
>start-trnH-GUG_Amaranthus_caudatus
ttgaatttggatg
>psbA-trnH-GUG_Amaranthus_caudatus
aatttcgtttttagtttagtatagatgagttattgaaagttaaaggagcaatgccgttttcttgttttgtcaagaaattggttattgctcctttattagt
>psbA-trnK-UUU-2_Amaranthus_caudatus
ggtaaaatcttggtttatttaatttaatcatcagggactcccaagcacattaattcttctcaaaatagagaattgaaagcttgttattcaacagtataac
>matK-trnK-UUU-2_Amaranthus_caudatus
ttttggttataagactatctaaatggaatggaaattatacctaaatgaatggaggtataagataaaaaatattattcattcatttctatactgaaatgtt
>matK-trnK-UUU-1_Amaranthus_caudatus
ttcaatttagaccgaaaacaaaagtaaaaatagagggtttcttggattatcaaatgatacatagtgcgataccgtcaaaacaaggcattttattacgaaa
>rps16-2-trnK-UUU-1_Amaranthus_caudatus
tgaactaaaaaacaaagataatgtggtggtagtttggtagaatttttttattccttattctcgtcctgtctattttttatgtagtgccaatccaacacaa
>rps16-1-rps16-2_Amaranthus_caudatus
gtgcgacttgaaggacatgatctggtgtggattcttatatccatcattctattctataagaaaggatgctcttgactcgacatcctttgctctgttccac
>rps16-1-trnQ-UUG_Amaranthus_caudatus
aacattcctcaaatttggaaccggtatgcggttgattcaattatggaatcatgaatagtcattggttcagtcaggacagtcaatacatatatattatatg
>psbK-trnQ-UUG_Amaranthus_caudatus
aacagacatttttttttattgcattttggttgagttattgatataaaaaaaaaatgaagaaaaaaatcgttcttttttttaacttactcactcaatcaaa
>psbI-psbK_Amaranthus_caudatus
tacacaatctccaagattattttttttggaaaaaagagaatagattctctatttttataccatatatcctataatataaaatataatttgaatttgaaat
>psbI-trnS-GCU_Amaranthus_caudatus
gactaaaaaggggttatttgctttatttttcatctatcttttttttttcaaaatagaattatatataattctattttgaaaaaaaaaaagatagaataa
>trnG-UCC-1-trnS-GCU_Amaranthus_caudatus
tacagttggattattgtatcgaaataaaacttttgtcgaacactgaaattggatgtaatcacgacaggattcgaaaagaatcatgaaatttagagtattg
```

2. Output -> Amaranthus_caudatus_intergenic.fasta, Amaranthus_tricolor_intergenic.fasta

| | | | |
|---|---|---|---|
| Amaranthus_caudatus_intergenic.fasta | 2023/4/11 19:19 | FASTA 文件 | 66 KB |
| Amaranthus_tricolor_intergenic.fasta | 2023/4/11 19:19 | FASTA 文件 | 74 KB |

Amaranthus_caudatus_intergenic.fasta

```
>start-trnH-GUG_Amaranthus_caudatus
ttgaatttggatg
>trnH-GUG-psbA_Amaranthus_caudatus
cccctacctaaaattcggaatttatttctaagttcagtattcagaacttgcctttgattaaatcttgaatccattttgaattaaaaaaaaaactgactag
>psbA-trnK-UUU-2_Amaranthus_caudatus
ggtaaaatcttggtttatttaatttaatcatcagggactcccaagcacattaattcttctcaaaatagagaattgaaagcttgttattcaacagtataac
>trnK-UUU-2-matK_Amaranthus_caudatus
atggagtagataatgttcttgttaaaaaagaaaaaacccctccccaaaccgtgcttgcatttttcattgcacacggctttccctatgtatacatttcaaa
>matK-trnK-UUU-1_Amaranthus_caudatus
ttcaatttagaccgaaaacaaaagtaaaaatagagggtttcttggattatcaaatgatacatagtgcgataccgtcaaaacaaggcattttattacgaaa
>trnK-UUU-1-rps16-2_Amaranthus_caudatus
gaaaaagagatatgttatgtagatacaatcgagataaaaataaaatgaattggaatgaaaactttgaattagcaagaaatcgaaacaaaagttgctaatt
>rps16-2-rps16-1_Amaranthus_caudatus
attgggatagattagatcaacaatacccccccctggaaacgtataggaggttttctcctcgtacggctcgagaaaaatgattcaaggtttaggtatataaa
>rps16-1-trnQ-UUG_Amaranthus_caudatus
aacattcctcaaatttggaaccggtatgcggttgattcaattatggaatcatgaatagtcattggttcagtcaggacagtcaatacatatatattatatg
>trnQ-UUG-psbK_Amaranthus_caudatus
attcgatttctattcaacattaataaacactaatattgttattgattgttcgtcaattccagcctaactatcaaaaaaaatcaacttgattctgttgtta
>psbK-psbI_Amaranthus_caudatus
gatctttcatcttgtcctaaaaaaatgaatgatttttcgagaaaaattattctaatattagaataataaataagaaataattgataaaatcagacaagt
>psbI-trnS-GCU_Amaranthus_caudatus
gactaaaaaggggttatttgctttatttttcatctatcttttttttttcaaaatagaattatatataattctattttgaaaaaaaaaaagatagaataa
>trnS-GCU-trnG-UCC-1_Amaranthus_caudatus
ccgttgaaaatagtaatagtcaaatattaaataataaatttcgaaaaatttgaaaattatgtaaaaaatatgtaataaactcaaattaattagactaaaa
```

#### 6.2. Remove Duplications

***Functionality***

Duplicated sequences can be removed from the same fasta format file(s). It is useful for processing the plastome with quadripartite structure, because the genes within the two IR (inverted repeat) copies are completely identical, and there are few genes that can be duplicated in the LSC (large single-copy) or SSC (small single-copy) regions. For the genes with duplicated genenames but non-duplicated sequences, it depends on users to judge how to process these potential duplicated sequences.

***Features***

1. Duplicated sequences from the same file can be batchly removed for multiple fasta files.
2. The genes with duplicated genenames and sequences will be automatically removed from the fasta files without prompting information.
3. The genes with duplicated genenames but non-duplicated sequences will not be removed, while prompting information will be shown in the warning.txt file.
4. The genes with non-duplicated genenames but duplicated sequences will not be removed, while prompting information will be shown in the warning.txt file.

***Command-line Tool***

perl remove_duplications.pl –i –o

run:

perl remove_duplications.pl –i input –o output

parameter:

[-i -input]   required: (default: input) input directory name containing fasta files for each species.

[-o -output]   required: (default: output) output directory name.

***Example***

1. Input -> Amaranthus_caudatus_CDS_RNA.fasta, Amaranthus_caudatus_intergenic.fasta, Amaranthus_tricolor_CDS_RNA.fasta, Amaranthus_tricolor_intergenic.fasta

| | | | |
|---|---|---|---|
| Amaranthus_caudatus_CDS_RNA.fasta | 2022/12/11 11:45 | FASTA 文件 | 92 KB |
| Amaranthus_caudatus_intergenic.fasta | 2022/12/11 21:43 | FASTA 文件 | 66 KB |
| Amaranthus_tricolor_CDS_RNA.fasta | 2022/12/11 11:45 | FASTA 文件 | 83 KB |
| Amaranthus_tricolor_intergenic.fasta | 2022/12/11 21:43 | FASTA 文件 | 74 KB |

Amaranthus_caudatus_CDS_RNA.fasta

```
>ndhG_Amaranthus_caudatus
atggatttacctggaccaatacatgattttcttttagtttttctggggtcgggccttatattaggtagtctaggcgtagtattatttaccaatacaattt
>ndhI_Amaranthus_caudatus
atgttccctatggtaactgggttcatcaattatggtcaacaaacaatacgtgcggcaaggtacattggtcaaggttttatgattactttatcccacgcta
>ndhA_Amaranthus_caudatus
atgataattgatataacaaaagtacaagctatcaattctttttccagattggaatccttaaaagaggtttatgggactatatggctgcttttccctattt
>ndhH_Amaranthus_caudatus
atgactgtaccaacttccagaaaagacctcatgatagtcaatatgggccctcaacatccatcaatgcatggcgttctccgactcatcgttactttagacg
>rps15_Amaranthus_caudatus
atgaaacaaaattcattcatatctgttatttcgaatgaaaaaaaagaagaaaatagggggtctgttgaatttcaagtattgtgttttaccaataagatac
>ycf1_Amaranthus_caudatus
atgattttcaatcttttctactaggtaatctagtatccttaggcatgaagataatcaattcggtcgttgtggtcggactctattatggatttctgacca
>trnN-GUU_Amaranthus_caudatus
tcctcagtagctcagtggtagagcggtcggctgttaaccgattggtcgtaggttcgaatcctacttggggag
>trnR-ACG_Amaranthus_caudatus
gggcctgtagctcagaggattagagcacgtggctacgaaccacggtgtcgggggttcgaatccctcctcgccca
>rrn5_Amaranthus_caudatus
tattctggtgtcctaggcgtagaggaaccacaccaatccatcccgaacttggtggttaaactctactgcggtgacgatactgtaggggaggtcctgcgga
>rrn4.5_Amaranthus_caudatus
gaaggtcacggcgagacgagccgtttatcattacgataggtgtcaagtggaagtgcagtgatgtatgcagctgaggcatcctaacagaccgatagacttg
>rrn23_Amaranthus_caudatus
ttcaaacgaggaaaggcttacggtggatacctaggcacccagagacgaggaagggcgtattaatcgacgaaatgcttcggggagttgaaaataagccgag
>trnA-UGC_Amaranthus_caudatus
ggggatatagctcagttggtagagcgccgctcttgcaaggcggatgtcagcggttcgagtccgcttatctcca
>trnI-GAU_Amaranthus_caudatus
gggctattagctcagtggtagagcgcgcccctgataagggcgaggtctctggttcaagtccaggatggccca
>rrn16_Amaranthus_caudatus
tctcatggagagttcgatcctggctcaggatgaacgctggcggcatgcttaacacatgcaagtcggacgggaagtggtgtttccagtggcggacgggtga
```

2. Output -> Amaranthus_caudatus_CDS_RNA.fasta, Amaranthus_caudatus_intergenic.fasta, Amaranthus_tricolor_CDS_RNA.fasta, Amaranthus_tricolor_intergenic.fasta, warning.txt

| | | | |
|---|---|---|---|
| Amaranthus_caudatus_CDS_RNA.fasta | 2023/4/11 22:36 | FASTA 文件 | 77 KB |
| Amaranthus_caudatus_intergenic.fasta | 2023/4/11 22:36 | FASTA 文件 | 58 KB |
| Amaranthus_tricolor_CDS_RNA.fasta | 2023/4/11 22:36 | FASTA 文件 | 74 KB |
| Amaranthus_tricolor_intergenic.fasta | 2023/4/11 22:36 | FASTA 文件 | 67 KB |
| warning.txt | 2023/4/11 22:36 | TXT 文件 | 1 KB |

Amaranthus_caudatus_CDS_RNA.fasta

```
>ndhG_Amaranthus_caudatus
atggatttacctggaccaatacatgattttcttttagtttttctggggtcgggccttatattaggtagtctaggcgtagtattatttaccaatacaattt
>ndhI_Amaranthus_caudatus
atgttccctatggtaactgggttcatcaattatggtcaacaaacaatacgtgcggcaaggtacattggtcaaggttttatgattactttatcccacgcta
>ndhA_Amaranthus_caudatus
atgataattgatataacaaaagtacaagctatcaattctttttccagattggaatccttaaaagaggtttatgggactatatggctgcttttccctattt
>ndhH_Amaranthus_caudatus
atgactgtaccaacttccagaaaagacctcatgatagtcaatatgggccctcaacatccatcaatgcatggcgttctccgactcatcgttactttagacg
>rps15_Amaranthus_caudatus
atgaaacaaaattcattcatatctgttatttcgaatgaaaaaaaagaagaaaatagggggtctgttgaatttcaagtattgtgttttaccaataagatac
>ycf1_Amaranthus_caudatus
atgattttcaatcttttctactaggtaatctagtatccttaggcatgaagataatcaattcggtcgttgtggtcggactctattatggatttctgacca
```

warning.txt

```
Not removed! Because in Amaranthus_caudatus_CDS_RNA.fasta, header >rps12_Amaranthus_caudatus is duplicated, but sequence is not duplicated.
Not removed! Because in Amaranthus_tricolor_CDS_RNA.fasta, header >rps12_Amaranthus_tricolor is duplicated, but sequence is not duplicated.
Not removed! Because in Amaranthus_tricolor_intergenic.fasta, header >trnN-GUU-trnR-ACG_Amaranthus_tricolor is duplicated, but sequence is not duplicated.
```

#### 6.3. Combine Fasta

***Functionality***

This tool can combine FASTA-formatted sequence files of multiple species.

***Features***

1. Any FASTA-formatted files will be combined together.

***Command-line Tool***

perl combine_fasta.pl –i –o

run:

perl combine_fasta.pl –i input –o output

parameter:

[-i -input] required: (default: input) input directory with multiple FASTA-formatted files.

[-o -output] required: (default: output) output directory with one combined file in FASTA-format.

***Example***

1. Input -> Amaranthus_caudatus_CDS_RNA.fasta, Amaranthus_tricolor_CDS_RNA.fasta

Amaranthus_caudatus_CDS_RNA.fasta

```
>ndhG_Amaranthus_caudatus
atggatttacctggaccaatacatgattttcttttagtttttctggggtcgggccttatattaggtagtctaggcgtagtattatttaccaatacaattt
>ndhI_Amaranthus_caudatus
atgttccctatggtaactgggttcatcaattatggtcaacaaacaatacgtgcggcaaggtacattggtcaaggttttatgattactttatcccacgcta
>ndhA_Amaranthus_caudatus
atgataattgatataacaaaagtacaagctatcaattctttttccagattggaatccttaaaagaggtttatgggactatatggctgcttttccctattt
>ndhH_Amaranthus_caudatus
atgactgtaccaacttccagaaaagacctcatgatagtcaatatgggccctcaacatccatcaatgcatggcgttctccgactcatcgttactttagacg
>rps15_Amaranthus_caudatus
atgaaacaaaattcattcatatctgttatttcgaatgaaaaaaaagaagaaaataggggtctgttgaatttcaagtattgtgttttaccaataagatac
>ycf1_Amaranthus_caudatus
atgattttcaatcttttctactaggtaatctagtatccttaggcatgaagataatcaattcggtcgttgtggtcggactctattatggatttctgacca
```

Amaranthus_tricolor_CDS_RNA.fasta

```
>ndhG_Amaranthus_tricolor
atggatttacctggaccaatacatgattttcttttagtttttctggggtcgggccttatattaggtggtctaggagtagtattatttaccaatacaatt
>ndhI_Amaranthus_tricolor
atgttccctatggtaactgggttcatcaattatggtcaacaaacaatacgtgcggcaaggtacattggtcaaggttttatgattactttatcccacgct
>ndhA_Amaranthus_tricolor
atgataattgatataacaaaagtacaagctatcaattctttttccagattggaatccttaaaagaggtttatgggactatatggctgcttttccctatt
>ndhH_Amaranthus_tricolor
atgactgtaccaacttccagaaaagacctcatgatagtcaatatgggccctcaccatccatcaatgcatggcgttctccgactcatcgttactttagac
>rps15_Amaranthus_tricolor
atgaaacaaaattcattcatatctgttatttcgaatgaaaaaaaagaagaaaataggggtctgttgaatttcaagtattgtgttttaccaataagata
```

2. Output -> all.fasta

```
>ndhG_Amaranthus_caudatus
atggatttacctggaccaatacatgattttcttttagtttttctggggtcgggccttatattaggtagtctaggcgtagtattatttaccaatacaattt
>ndhI_Amaranthus_caudatus
atgttccctatggtaactgggttcatcaattatggtcaacaaacaatacgtgcggcaaggtacattggtcaaggttttatgattactttatcccacgcta
>ndhA_Amaranthus_caudatus
atgataattgatataacaaaagtacaagctatcaattctttttccagattggaatccttaaaagaggtttatgggactatatggctgcttttccctattt
>ndhH_Amaranthus_caudatus
atgactgtaccaacttccagaaaagacctcatgatagtcaatatgggccctcaacatccatcaatgcatggcgttctccgactcatcgttactttagacg
>rps15_Amaranthus_caudatus
atgaaacaaaattcattcatatctgttatttcgaatgaaaaaaaagaagaaaataggggtctgttgaatttcaagtattgtgttttaccaataagatac
>ycf1_Amaranthus_caudatus
atgattttcaatcttttctactaggtaatctagtatccttaggcatgaagataatcaattcggtcgttgtggtcggactctattatggatttctgacca
>ndhG_Amaranthus_tricolor
atggatttacctggaccaatacatgattttcttttagtttttctggggtcgggccttatattaggtggtctaggagtagtattatttaccaatacaatt
>ndhI_Amaranthus_tricolor
atgttccctatggtaactgggttcatcaattatggtcaacaaacaatacgtgcggcaaggtacattggtcaaggttttatgattactttatcccacgct
>ndhA_Amaranthus_tricolor
atgataattgatataacaaaagtacaagctatcaattctttttccagattggaatccttaaaagaggtttatgggactatatggctgcttttccctatt
>ndhH_Amaranthus_tricolor
atgactgtaccaacttccagaaaagacctcatgatagtcaatatgggccctcaccatccatcaatgcatggcgttctccgactcatcgttactttagac
>rps15_Amaranthus_tricolor
atgaaacaaaattcattcatatctgttatttcgaatgaaaaaaaagaagaaaataggggtctgttgaatttcaagtattgtgttttaccaataagata
```

##### 6.4. Generate Matrices

***Functionality***

This tool can generate sequence matrices for each coding region and noncoding region of plastomes for multiple species. The generated sequence matrices will be easy to be aligned by alignment tools.

***Features***

1. The first file (.fasta format) containing multiple sequences from multiple species must be uploaded.

2. The second file (.txt format) containing sequence names can be optionally provided. If the second file is not provided, the number of sequence matrices in the output directory will be euqal to the number of sequence names in the first file. If the second file is provided, the number of sequence matrices in the output directory will be equal to or less than the number of sequence names in the second file. That is to say, if the sequence names in the second file are all included in the first file, the number of sequence matrices in the output directory will be equal to the number of sequence names in the second file; if the sequence names in the second file are partially included in the first file, the number of sequence matrices in the output directory will be less than the number of sequence names in the second file, and only the sequence names shared by the second file and the first file are occurred in the output directory.

3. The header name in the first file (.fasta format) must be like “>*psbA*_Amaranthus_caudatus”, i.e., “>sequence name_species name”. The sequence name in the second file (.txt format) must be like “*psbA*”.

***Command-line Tool***

perl generate_matrices.pl –i [–n] –o

run:
perl generate_matrices.pl –i all.fasta –n codingname.txt –o output
perl generate_matrices.pl –i all.fasta –o output

parameter:
[-i -input] required: (default: input) input filename containing sequences from multiple species.
[-n -name] optional: (default: codingname.txt/noncodingname.txt) input filename containing sequence names or not assign this file.
[-o -output] required: (default: output) output directory.

***Example***

1. Input -> all.fasta

```
>trnH-GUG_Amaranthus_caudatus
gcggatgtagccaagtggatcaaggcagtggattgtgaatccaccacgcgcgggttcaattcccgtcgttcgcc
>psbA_Amaranthus_caudatus
atgactgcaattttagagagacgcgaaagcgaaagcctatggggtcgtttctgtaactggataaccagcactgaaaaccgtctttacatcggatggtttggtgttttgatg
>trnK-UUU_Amaranthus_caudatus
gggttgctaactcaatggtagagtactcggcttttaaccgatcggttccgggttcgagtcccgggcaaccca
>matK_Amaranthus_caudatus
atggaaaaattacaaggacatagagaactagataggtcttggcaacataactttttctatccacttatctttcaggaatatatttatgtatttgcatatgatcatgcttta
>rps16_Amaranthus_caudatus
atggtgaaacttcgtttgaaacgatgtggtagaaagcaacgagccgtctatcgaattgttgcaattgatgttcggtcccgaagagagggaagagatttgcagaaagtgggt
>trnQ-UUG_Amaranthus_caudatus
tggggcgtggccaagtggtaaggcatcgggttttggtcccgctattcggaggttcgaatccttccgtcccag
>psbK_Amaranthus_caudatus
atgcttaatatctttagtttgatctgtcttaattcggccctttattcgagtagtttttctttgccaaattacctgaggcctacgctttttgagtccaattgtagatttt
>psbI_Amaranthus_caudatus
atgcttactctcaaactctttgtttacacagtagtaatatttttgtttctctcttcattttcggattcctatctaatgatccaggacgtaatcctgggcgcgaagaataa
>trnS-GCU_Amaranthus_caudatus
ggagagatggctgagtggacgaaagcgtcggattgctaatccgttgtacgagttattcgtaccgagggttcgaatccctctctttccg
>trnG-UCC_Amaranthus_caudatus
gcgggtatagtttagtggtaaaaccctagccttccaagctaacgatgcgggttcgattcccgctacccgct
```

```
>trnH-GUG_Amaranthus_tricolor
gcggatgtagccaagtggatcaaggcagtggattgtgaatccaccacgcgcgggttcaattcccgtcgttcgcc
>psbA_Amaranthus_tricolor
atgactgcaattttagagagacgcgaaagcgaaagcctatggggtcgtttctgtaactggataaccagcactgaaaaccgtctttacatcggatggtttggtgttttgatg
>trnK-UUU_Amaranthus_tricolor
gggttgctaactcaatggtagagtactcggcttttaaccgatcggttccgggttcgagtcccgggcaaccca
>matK_Amaranthus_tricolor
atggaaaaattacaaggacatagagaactagataggtcttggcaacataactttttctatccacttatctttcaggaatatatttatgtatttgcatatgatcatgcttta
>rps16_Amaranthus_tricolor
atggtgaaacttcgtttgaaacgatgtggtagaaagcaacgagccgtctatcgaattgttgcaattgatgttcggtcccgaagagagggaagagatttgcagaaagtgggt
>trnQ-UUG_Amaranthus_tricolor
tggggcgtggccaagtggtaaggcatcgggttttggtcccgctattcggaggttcgaatccttccgtcccag
>psbK_Amaranthus_tricolor
atgcttaatatctttagtttgatctgtcttaattcggccctttattcgagtagtttttctttgccaaattaccggaggcctacgctttttgagtccaattgtagatttt
>psbI_Amaranthus_tricolor
atgcttactctcaaacttttgtttacacagtagtaatatttttgtttctcttttcattttcggattcctatctaatgatccaggacgtaatcctgggcgcgaagaataa
>trnS-GCU_Amaranthus_tricolor
ggagagatggctgagtggacgaaagcgtcggattgctaatccgttgtacgagttattcgtaccgagggttcgaatccctctctttccg
>trnG-UCC_Amaranthus_tricolor
gcgggtatagtttagtggtaaaaccctagccttccaagctaacgatgcgggttcgattcccgctacccgct
```

2. Input -> codingname.txt

```
accD
atpA
atpB
atpE
atpF
atpH
atpI
ccsA
cemA
clpP
```

3. Output -> accD.fasta, atpA.fasta, …, clpP.fasta

| | | | |
|---|---|---|---|
| accD.fasta | 2023/4/12 9:51 | FASTA 文件 | 4 KB |
| atpA.fasta | 2023/4/12 9:51 | FASTA 文件 | 4 KB |
| atpB.fasta | 2023/4/12 9:51 | FASTA 文件 | 3 KB |
| atpE.fasta | 2023/4/12 9:51 | FASTA 文件 | 1 KB |
| atpF.fasta | 2023/4/12 9:51 | FASTA 文件 | 2 KB |
| atpH.fasta | 2023/4/12 9:51 | FASTA 文件 | 1 KB |
| atpI.fasta | 2023/4/12 9:51 | FASTA 文件 | 2 KB |
| ccsA.fasta | 2023/4/12 9:51 | FASTA 文件 | 2 KB |
| cemA.fasta | 2023/4/12 9:51 | FASTA 文件 | 2 KB |
| clpP.fasta | 2023/4/12 9:51 | FASTA 文件 | 2 KB |

accD.fasta

```
>Amaranthus_caudatus
atgcttattcggacaagaaatgttcaaattgattttcatcgaatgactattcatctattttattttcatgcgaatagggggcaagaaagctctatgaaaa
>Amaranthus_tricolor
atgcttattctgacaagaaatgttcaaattgattttcatcgaatgactattcatctattttattttcatgcgaatagggggcaagaaagctctatgaaaa
```

##### 6.5. Check Codons

***Functionality***

This tool can batchly check the bad codons in the sequence matrices of PCGs (protein-coding genes).

***Features***

1. The bad codons can be batchly checked for multiple PCG matrices.
2. Only PCG genes (i.e., linked exons and without introns) can be checked. That is to say, the intron sequences are not allowed in the sequences.

***Command-line Tool***

perl check_codons.pl –i –o

run:

perl check_codons.pl –i input –o output

parameter:

| | |
|---|---|
| [-i -input] | required: (default: input) input directory containing .fasta PCG matrices. |
| [-o -output] | required: (default: output) output directory. |

***Example***

1. Input -> atpA.fasta, atpE.fasta

| | | | |
|---|---|---|---|
| atpA.fasta | 2022/12/11 17:49 | FASTA 文件 | 4 KB |
| atpE.fasta | 2022/12/11 16:46 | FASTA 文件 | 1 KB |

atpA.fasta

```
>Amaranthus_caudatus
atggtaaccattcgagcagacgaaattagcaatattatccgtgaacgtattgaacaataaaatcgagaagtaaaggttgtaaataccggtaccgtacttc
>Amaranthus_tricolor
atggtaaccattcgagcagacgaaattagcaatattatccgtgaacgtattgaacaatataatcgagaagtaaaggttgtaaataccggtaccgtacttc
```

2. Output -> bad_codons.txt

```
Bad codon TAA in position 58 of atpA in species Amaranthus_caudatus!
```

##### 6.6. Check Missings

***Functionality***

This tool can batchly check the missing taxa from multiple sequence matrices.

***Features***

1. The missing taxa can be batchly checked for multiple sequence matrices.
2. The missing taxa of each sequence matrix will be listed in the missing_taxa.txt file.

***Command-line Tool***

perl check_missings.pl –i –o

run:
perl check_missings.pl –i input –o output

parameter:

| | |
|---|---|
| [-i -input] | required: (default: input) input directory containing .fasta sequence matrices. |
| [-o -output] | required: (default: output) output directory. |

***Example***

1. Input -> accD.fasta, atpA.fasta, atpB.fasta

| | | | |
|---|---|---|---|
| accD.fasta | 2022/12/11 17:23 | FASTA 文件 | 2 KB |
| atpA.fasta | 2022/12/11 17:49 | FASTA 文件 | 4 KB |
| atpB.fasta | 2022/12/11 17:23 | FASTA 文件 | 2 KB |

accD.fasta

```
>Amaranthus_caudatus
atgcttattcggacaagaaatgttcaaattgattttcatcgaatgactattcatctattttattttcatgcgaatagggggcaagaaagctctatgaaaa

```

atpA.fasta

```
>Amaranthus_caudatus
atggtaaccattcgagcagacgaaattagcaatattatccgtgaacgtattgaacaataaaatcgagaagtaaaggttgtaaataccggtaccgtacttc
>Amaranthus_tricolor
atggtaaccattcgagcagacgaaattagcaatattatccgtgaacgtattgaacaatataatcgagaagtaaaggttgtaaataccggtaccgtacttc

```

atpB.fasta

```
>Amaranthus_tricolor
atgacaatcaatcctactacttctggttctggggtttccacgcttgaaaaaaaaaacctggggcgtatcgctcaaattattggtccggtcctggacgtag

```

2. Output -> missing_taxa.txt

```
accD
Amaranthus_tricolor
atpA

atpB
Amaranthus_caudatus
```

##### 6.7. Extract Sites

***Functionality***

This tool can batchly extract each codon site or codon sites of PCGs (protein-coding genes).

***Features***

1. The codon sites can be batchly extracted for multiple PCG matrices.

2. Only PCG genes (i.e., linked exons and without introns) can be extracted. That is to say, intron sequences are not allowed in the sequences.

3. Three extraction patterns of codon sites are developed. If the extraction pattern 1 (site1 + site2 + site3) is selected, it will generate three fasta format files, i.e., "sequence name_site1.fasta", "sequence name_site2.fasta", and "sequence name_site3.fasta"; if the extraction pattern 2 (site12 + site3) is selected, it will generate two fasta format files, i.e., "sequence name_site12.fasta" and "sequence name_site3.fasta"; if the extraction pattern 3 (site1 + site23) is selected, it will generate two fasta format files, i.e., "sequence name_site1.fasta" and "sequence name_site23.fasta".

4. In the mean time, the bad codons can be batchly checked for multiple PCG matrices. And the bad codons of each sequence will be listed in the bad_codons.txt file.

***Command-line Tool***

perl extract_sites.pl –i –s –o

run:
perl extract_sites.pl –i input –s 1 –o output

parameter:
[-i -input]    required: (default: input) input directory.
[-s -site]    required: (default: 1) 1/2/3, extraction pattern of codon sites, 1=site1 + site2 + site3, 2=site12 + site3, 3=site1 + site23.
[-o -output]    required: (default: output) output directory.

***Example***

1. Input -> atpA.fasta, atpE.fasta

| | | | |
|---|---|---|---|
| atpA.fasta | 2022/12/11 17:49 | FASTA 文件 | 4 KB |
| atpE.fasta | 2022/12/11 16:46 | FASTA 文件 | 1 KB |

atpA.fasta

```
>Amaranthus_caudatus
atggtaaccattcgagcagacgaaattagcaatattatccgtgaacgtattgaacaataaaatcgagaagtaaaggttgtaaataccggtaccgtacttc
>Amaranthus_tricolor
atggtaaccattcgagcagacgaaattagcaatattatccgtgaacgtattgaacaatataatcgagaagtaaaggttgtaaataccggtaccgtacttc
```

2. Output -> atpA_site1.fasta, atpA_site2.fasta, atpA_site3.fasta, …, bad_codons.txt

| | | | |
|---|---|---|---|
| atpA_site1.fasta | 2023/4/12 11:17 | FASTA 文件 | 2 KB |
| atpA_site2.fasta | 2023/4/12 11:17 | FASTA 文件 | 2 KB |
| atpA_site3.fasta | 2023/4/12 11:17 | FASTA 文件 | 2 KB |
| atpE_site1.fasta | 2023/4/12 11:17 | FASTA 文件 | 1 KB |
| atpE_site2.fasta | 2023/4/12 11:17 | FASTA 文件 | 1 KB |
| atpE_site3.fasta | 2023/4/12 11:17 | FASTA 文件 | 1 KB |
| bad_codons.txt | 2023/4/12 11:17 | TXT 文件 | 1 KB |

atpA_site1.fasta

```
>Amaranthus_caudatus
agaacgggaaaaacgcagctacggaggaagagccggggagcacgcgggagggtggtgggaagagcatgtaaggggtagggtcacggatgagagaagcacg
>Amaranthus_tricolor
agaacgggaaaaacgcagctacggaggaagagccggggagcacgcgggagggtggtcggaagagtatgtaaggggtagggtcacggatgagagaagcacg
```

atpA_site2.fasta

```
>Amaranthus_caudatus
ttctgcaatgattgagtaaaagatattacgcttatgagtcgtagtaattcgattataagctgtctatacaatgttttgagtttaaggctaccggtcatct
>Amaranthus_tricolor
ttctgcaatgattgagtaaaagatattacgcttatgagtcgtagtaattcgattataagctgtctatacaatgttttgagtttaaggctaccggtcatct
```

atpA_site3.fasta

```
>Amaranthus_caudatus
gactaacatcttctattaaataaagtatctcataatccttttctttaagataaaataggaacttgtgactttttaagtttggaaaattaaaaaattgatg
>Amaranthus_tricolor
gactaacatcttctattaattaaagtatctcataatccttttctttaagataaaataggaacttgtgactttttaagtttggaaaattaaaaaattgatg
```

bad_codons.txt

```
Bad codon TAA in position 58 of atpA in species >Amaranthus_caudatus!
```

##### 6.8. Codon-to-AA

***Functionality***

This tool can batchly translate codons to amino acids for PCGs (protein-coding genes). The generated amino acids sequence matrices can be aligned by alignment tools.

***Features***

1. Codons can be batchly translated to amino acids for multiple PCGs.
2. Only PCG genes (i.e., linked exons and without introns) can be translated. That is to say, intron sequences are not allowed in the sequences.
3. In the mean time, the bad codons can be batchly checked for multiple PCG matrices. And the bad codons of each sequence will be listed in the bad_codons.txt file.

***Command-line Tool***

perl codon_to_aa.pl –i –o

run:

perl codon_to_aa.pl –i input –o output

parameter:

| | |
|---|---|
| [-i -input] | required: (default: input) input directory name containing .fasta files. |
| [-o -output] | required: (default: output) output directory name. |

***Example***

1. Input -> atpA.fasta, atpE.fasta

| | | | |
|---|---|---|---|
| atpA.fasta | 2022/12/11 17:49 | FASTA 文件 | 4 KB |
| atpE.fasta | 2022/12/11 16:46 | FASTA 文件 | 1 KB |

atpA.fasta

```
>Amaranthus_caudatus
atggtaaccattcgagcagacgaaattagcaatattatccgtgaacgtattgaacaataaaatcgagaagtaaaggttgtaaataccggtaccgtacttc
>Amaranthus_tricolor
atggtaaccattcgagcagacgaaattagcaatattatccgtgaacgtattgaacaatataatcgagaagtaaaggttgtaaataccggtaccgtacttc
```

2. Output - > atpA.fasta, atpE.fasta, bad_codons.txt

| | | | |
|---|---|---|---|
| atpA.fasta | 2022/12/13 12:12 | FASTA 文件 | 2 KB |
| atpE.fasta | 2022/12/13 12:12 | FASTA 文件 | 1 KB |
| bad_codons.txt | 2022/12/13 12:12 | TXT 文件 | 1 KB |

atpA.fasta

```
>Amaranthus_caudatus
MVTIRADEISNIIRERIEQ*NREVKVVNTGTVLQVGDGIARIHGLDEVMAGELVEFEEGTIGIALNLESNNVGVVLMGDGLLIQEGSSVKATGRIAQIPV
>Amaranthus_tricolor
MVTIRADEISNIIRERIEQYNREVKVVNTGTVLQVGDGIARIHGLDEVMAGELVEFQEGTIGIALNLESNNVGVVLMGDGLLIQEGSSVKATGRIAQIPV
```

bad_codons.txt

```
Bad codon TAA in position 58 of atpA in species Amaranthus_caudatus!
```

#### 7. (Post)-Alignment

This tool kit contains six tools that can complete alignment and post-alignment processing for the sequence matrices, such as "7.1. Sequence Alignment, 7.2. Generate Non-interleaved, 7.3. Concatenate Alignments, 7.4. Extract Sub-alignments, 7.5. Remove Gaps, and 7.6. Generate PartitionFinder".

##### 7.1. Sequence Alignment

***Functionality***

This tool can batchly do alignment for multiple sequence matrices by the frequently used mafft alignment tool.

***Features***

1. Multiple sequence matrices can be batchly aligned by the frequently used built-in alignment tool–mafft. If you use it, please cite following paper.

Katoh, K., & Standley, D. M. (2013). MAFFT multiple sequence alignment software version 7: improvements in performance and usability. Molecular Biology and Evolution, 30(4), 772-780. https://doi.org/10.1093/molbev/mst010.

2. The alignment command of mafft will be set to this mode "mafft --localpair --maxiterate 16 --inputorder input > output".

3. If duplicated species are existed within the same sequence matrix file, mafft alignment will be normally run, and the warning information like "Duplicated species name Arabidopsis_thaliana for the gene atpA!" will be outputed to the "warning.txt" file.

4. If the sequence matrix with species number <= 50 and sequence length <= 10,000, mafft alignment will be normally run; if the sequence matrix with species number > 50 or if the sequence matrix with sequence length > 10,000, mafft alignment will not be run, and the warning information like "Species number larger than 50 for the gene atpA!" or "Sequence length longer than 10 kb for the gene atpA!" will be outputed to the "warning.txt" file.

***Command-line Tool***

```
perl sequence_alignment.pl –i –o

run:
perl sequence_alignment.pl –i input –o output

parameter:
[-i -input]     required: (default: input) input directory containing .fasta sequence matrices.
[-o -output]    required: (default: output) output directory.
```

***Example***

1. Input -> accD.fasta, accD_aa.fasta, atpA.fasta, atpA_aa.fasta

| | | | |
|---|---|---|---|
| accD.fasta | 2022/12/14 10:21 | FASTA 文件 | 10 KB |
| accD_aa.fasta | 2022/12/14 10:21 | FASTA 文件 | 4 KB |
| atpA.fasta | 2022/12/14 10:21 | FASTA 文件 | 10 KB |
| atpA_aa.fasta | 2022/12/14 10:21 | FASTA 文件 | 4 KB |

accD.fasta

```
>Amaranthus_caudatus
atgcttattcggacaagaaatgttcaaattgattttcatcgaatgactattcatctattttattttcatgcgaatagggggcaagaaagctctatgaaaa
>Amaranthus_tricolor
atgcttattctgacaagaaatgttcaaattgattttcatcgaatgactattcatctattttattttcatgcgaatagggggcaagaaagctctatgaaaa
>Atriplex_centralasiatica
atgaaaaaggattggttcaattcgatcttgtctaaaaaagagttagaacgcagatacgggcgtggcttaagtaaatcaatggacagtcttggtcctactg
>Chenopodium_album
atgaaaaagggttggttcaattcgatcttgtctaaaaaagagttagaacacagatacgggtgtggattaagtaaatcaatggacagtcttggtcctactg
>Chenopodium_quinoa
atgaaaaagggttggttcaattcgatcttgtctaaaaaagagttagaacacagatacgggtgtggattaagtaaatcaatggacagtcttggtcctactg
>Salicornia_bigelovii
atgaaaaaagggtggttcaattcgctgttgtctaagaaagaattagaacacagatgtgcattaagtaaatcactggatagtcttggtcctattgaaaata
```

2. Output -> accD.fasta, accD_aa.fasta, atpA.fasta, atpA_aa.fasta

| | | | |
|---|---|---|---|
| accD.fasta | 2022/12/14 10:22 | FASTA 文件 | 11 KB |
| accD_aa.fasta | 2022/12/14 10:22 | FASTA 文件 | 4 KB |
| atpA.fasta | 2022/12/14 10:22 | FASTA 文件 | 10 KB |
| atpA_aa.fasta | 2022/12/14 10:23 | FASTA 文件 | 4 KB |

accD.fasta

```
>Amaranthus_caudatus
atgcttattcggacaagaaatgttcaaattgattttcatcgaatgactattcatctattt
tattttcatgcgaatagggggcaagaaagctctatgaaaaaatggtggttcaattcgatg
ttgtctaagaaagggttcgaacata------agtgtggattaagtaaatcaatggacagt
cttggtcctattgaaaataccagtagaaaggaagatccgagtctaaatgatacagaaaaa
agcattcatagttggagaaatggtgacaattctagttacagcagtaatattgataatttc
tttcgtgtcatg------gacattccgaatttcatctctgatgatactttttttacttata
gatcgtagggggggacagttattccatatattttgatattgaaaatcaaatttttgagatt
gataacgatcagtcttttctgagtgaactacaaagttctttttcgaattattggacttca
agttatctgaactcgaaatctaagagtgacgatacttataacgatcgttccggatattat
acta---aagatagttggaataatcacattaataattgcattgacagttatcttcattct
caaatcagtattgggagttccatcctaagtggtagtgacaattccagtgacagttacatt
ttgagttacatttttaatgaaagtggaaat---aggagtgaaattttcagt---------
------------------------aaaagaacgagcacaaatgatagtaatttacctaga
atagaaagttctaataatcttgatgtaactcaaaaatataaacatttgtgggttcaatgt
gaaaattgttatgcattaaattataagaaattgcttaagtcaaaaatgggtatttgtgaa
caatgtggatatcatttgaaaattaatagttcagatagaatcgaacttctgattgatccg
ggtacttggaatcctatggatgaagatatggtctctacggatcctattgaatttcattca
gaggaggaagcttataaagatcgtattgattcttatcaaacaaagacaggtttaactgaa
gctgttcaaacaggtataggtcaaataaatggtatttctatagcaattggggttatggat
```

3. Output -> warning.txt

Duplicated species name Amaranthus_caudatus for the gene atpA!

Species number larger than 50 for the gene atpA!

Sequence length longer than 10kb for the gene atpB!

#### 7.2. Generate Non-interleaved

***Functionality***

This tool will batchly generate non-interleaved fasta format files from interleaved fasta format files.

***Features***

1. Interleaved fasta format files can be batchly converted to non-interleaved fasta format files.
2. Whether there are nucleotides or amino acids in the fasta format files, this tool can generated non-interleaved fasta format files.

***Command-line Tool***

perl generate_non-interleaved.pl –i –o

run:

perl generate_non-interleaved.pl –i input –o output

parameter:

[-i -input]     required: (default: input) input directory containing interleaved .fasta file.

[-o -output]     required: (default: output) output directory.

***Example***

1. Input -> atpE.fasta

```
>Amaranthus_caudatus
atgaccttaaatctttgtgtactgactcctaatcgaagtgtttggaattcacaagtaaaa
gcaatcattttatctactaatagtggccaaatcggcgtattaaaagaccatgccgctact
gcgacagccgtagatataggtgttttaagaatgttaggccttgacgaccaatggtcaaca
atggctctgatgggcggttttgctagaataagcaataatcagattactattttagtaaat
gatgctgagaagggtagtgacattgatccacaagaagctcaggaaactcttgaaatagca
gaagctaacttgaagaaggcggaaggaaagagacaattaattgaagcaaatctagctctc
agacgagctaggacacgagtagaggctcgcaatacaatttcgtaa
>Amaranthus_tricolor
atgaccttaaatctttgtgtactgactcctaatcgaagtgtttggaattcacaagtaaaa
gcaatcattttatctactaatagtggccaaatcggcgtattaaaagaccatgccgctact
gcgacagccgtagatataggtgttttaagaatgttaggccttgacgaccaatggtcaaca
atggctctgatgggcggttttgctagaataagcaataatcagattactattttagtaaat
gatgctgagaagggtagtgacattgatccacaagaagctcaggaaactcttgaaatagca
gaagctaacttgaagaaggcggaaggaaagagacaattaattgaagcaaatctagctctc
agacgagctaggacacgagtagaggctcgcaatacaatttcgtaa
```

2. Output -> atpE.fasta

```
>Amaranthus_caudatus
atgaccttaaatctttgtgtactgactcctaatcgaagtgtttggaattcacaagtaaaagcaatcattttatctactaatagtggccaaatcggcgtatt
>Amaranthus_tricolor
atgaccttaaatctttgtgtactgactcctaatcgaagtgtttggaattcacaagtaaaagcaatcattttatctactaatagtggccaaatcggcgtatt
```

##### 7.3. Extract Sub-alignments

***Functionality***

This tool can batchly extract sub-alignments from fasta format sequence alignment matrices based on assigned species names. In the mean time, this tool can batchly remove gap sites in the sub-alignments.

***Features***

1. The sub-alignments can be batchly extracted from the alignment matrices.
2. Non-aligned sequence matrices cannot be allowed as input files.
3. Nucleotides, amino acids, upper case letters, lower case letters, gap (–), and special letters such as “N”, “n”, “X”, “x” (except 4 types of nucleotides, i.e., A, T, G, C, and 20 types of amino acids, i.e., A, C, D, E, F, G, H, I, K, L, M, N, P, Q, R, S, T, V, W, Y), are allowed in the alignment matrices.
4. A “list.txt” file containing assigned species names must be provided.
5. The gap sites in the sub-alignments can be simultaneously removed.
6. A “warning.txt” file may be generated for giving warning information.
7. For list.txt, if there are no species names that are not in the alignment matrices, the “warning.txt” file will not be generated; if there are species names that are not in the alignment matrices, the “warning.txt” file will be generated, and the warning information like “Alignment matrix atpB.fasta does not contain following species:” will be shown.

***Command-line Tool***

perl extract_sub-alignments.pl –i –l –o

run:

perl extract_sub-alignments.pl –i input –l list.txt –o output

parameter:

[-i -input] required: (default: input) input directory containing .fasta alignment matrices.

[-l -list] required: (default: list.txt) input list filename containing species names.

[-o -output] required: (default: output) output directory.

***Example***

1. Input -> atpA.fasta, atpB.fasta, atpF.fasta

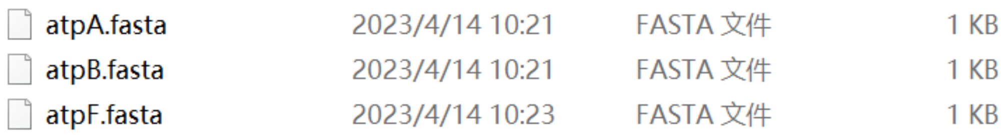

atpA.fasta

```
>sp1
atgtgtgaaaaagacaaagattctgagaaaatgtat------------------------------gaagaagacaaagattctgaggaaatg
>sp2
atgtgt------------------------------------------------------------gaagaaaacaaagattctgaaaaaatg
>sp3
atgtgt------------------------------------------------------------gaggaaaac------------------
>sp4
atgtgt------------------------------------------------------------gaagaagacaaagattctgagaaaatg
>sp5
atgttt------------------------------------------------------------gaggaaaacaaggattctgaaaatatt
>sp6
atgtgtgaaaaagacaaagattctgagaaaatgtatgaagaagacaaagattctgaggaaatgtgtgaagaagacaaagattctgaggaaatg
>sp7
atgtgtgaagaaaacaaagattctgagaaaatg------------------------------tgtgaagaagacaaagattctgagaaaatg
>sp8
atgtgt------------------------------------------------------------gaagaaaacaaagattctgaaaaaatg
```

atpB.fasta

```
>sp1
ATGTGTGAA-N-GACAAAGATTCTGAGAAAATGTATGAAGAAGACAAAGATTCTGAGGAAATGTGTGAAGAAGACAAA
>sp2
ATGTGTGAA-n-AACAAAGATTCTGAAAAAATGTGTGAAGAAGACAAAGATTCTGAAAAAATGTGTGAAGAAGACAAA
>sp3
ATGTGTGAG---AACGAAAACAAGGATTCTGAATATATCGAAACAACCCCCGTAGAAAACGGATACAATTCGGAACGT
>sp8
ATGTGTGAAGAAGACAAAGATTCTGAAAAAATGTGTGAAGAAGACAAAGGTTCTGAAAAAATGTGTCAGGAAAACAAA
>sp9
ATGTGTGAAGAAAACAAAGATTCTGAGAAAATGTGTGAAGAAGACAAAGATTCTGAGAAAATGTGTGAAGAAGACAAA
```

atpF.fasta

```
>sp1
MTLNL-----TPNRSVWNSQVKAIILSTNSGQIGVLKDHAATATAVDIGVLRMLGLDDQWSTMALMGGFARI
>sp2
mtlnl-----tpnrsvwnsqvkaiilstnsgqigvlkdhaatatavdigvlrmlglddqwstmalmggfari
>sp5
MTLNL--X--TPNRSVWNSQVKAIILSTNSGQIGVLKDHAATATAVDIGVLRMLGLDDQWSTMALMGGFARI
>sp6
mtlnl--x--tpnrsvwnsqvkaiilstnsgqigvlkdhaatatavdigvlrmlglddqwstmalmggfari
>sp10
MTLNLMSCVLTPNRSVWNSQVKAIILSTNSGQIGVLKDHAATATAVDIGVLRMLGLDDQWSTMALMGGFARI
```

2. Input -> list.txt

```
sp1
sp2
sp3
sp4
sp5
sp8
```

3. Output -> atpA.fasta, atpB.fasta, atpF.fasta, warning.txt

| | | | |
|---|---|---|---|
| atpA.fasta | 2023/4/14 10:41 | FASTA 文件 | 1 KB |
| atpB.fasta | 2023/4/14 10:41 | FASTA 文件 | 1 KB |
| atpF.fasta | 2023/4/14 10:41 | FASTA 文件 | 1 KB |
| warning.txt | 2023/4/14 10:41 | TXT 文件 | 1 KB |

atpA.fasta

```
>sp1
ATGTGTGAAAAAGACAAAGATTCTGAGAAAATGTATGAAGAAGACAAAGATTCTGAGGAAATG
>sp2
ATGTGT------------------------------GAAGAAAACAAAGATTCTGAAAAAATG
>sp3
ATGTGT------------------------------GAGGAAAAC------------------
>sp4
ATGTGT------------------------------GAAGAAGACAAAGATTCTGAGAAAATG
>sp5
ATGTTT------------------------------GAGGAAAACAAGGATTCTGAAAATATT
>sp8
ATGTGT------------------------------GAAGAAAACAAAGATTCTGAAAAAATG
```

atpB.fasta

```
>sp1
ATGTGTGAA-N-GACAAAGATTCTGAGAAAATGTATGAAGAAGACAAAGATTCTGAGGAAATGTGTGAAGAAGACAAA
>sp2
ATGTGTGAA-N-AACAAAGATTCTGAAAAAATGTGTGAAGAAGACAAAGATTCTGAAAAAATGTGTGAAGAAGACAAA
>sp3
ATGTGTGAG---AACGAAAACAAGGATTCTGAATATATCGAAACAACCCCCGTAGAAAACGGATACAATTCGGAACGT
>sp8
ATGTGTGAAGAAGACAAAGATTCTGAAAAAATGTGTGAAGAAGACAAAGGTTCTGAAAAAATGTGTCAGGAAAACAAA
```

atpF.fasta

```
>sp1
MTLNLTPNRSVWNSQVKAIILSTNSGQIGVLKDHAATATAVDIGVLRMLGLDDQWSTMALMGGFARI
>sp2
MTLNLTPNRSVWNSQVKAIILSTNSGQIGVLKDHAATATAVDIGVLRMLGLDDQWSTMALMGGFARI
>sp5
MTLNLTPNRSVWNSQVKAIILSTNSGQIGVLKDHAATATAVDIGVLRMLGLDDQWSTMALMGGFARI
```

warning.txt

```
Alignment matrix atpB.fasta does not contain following species:
sp4 sp5
Alignment matrix atpF.fasta does not contain following species:
sp3 sp4 sp8
```

#### 7.4. Concatenate Alignments

***Functionality***

This tool can concatenate all fasta format alignment matrices.

***Features***

1. All fasta format files must be alignment matrices. That is to say, all files must be aligned before the concatenation.

2. Missing species are allowed for any fasta format files, and the gaps for the species with missing genes will be displayed with multiple dash (–) lines.

3. If some alignment files contain nucleotide sequences, while others contain amino acid sequences, all of these files will be automatically concatenated without judgement. Because this tool does not determine whether nucleotides and amino acids can be concatenated.

***Command-line Tool***

perl concatenate_alignments.pl –i –o

run:

perl concatenate_alignments.pl –i input –o output

parameter:

[-i -input]  required: (default: input) input directory containing .fasta file(s).

[-o -output]  required: (default: all) output filename.

***Example***

1. Input -> atpA.fasta, atpB.fasta, atpE.fasta

| | | | |
|---|---|---|---|
| atpA.fasta | 2023/4/14 0:12 | FASTA 文件 | 1 KB |
| atpB.fasta | 2023/4/14 0:11 | FASTA 文件 | 1 KB |
| atpE.fasta | 2023/4/14 0:15 | FASTA 文件 | 1 KB |

atpA.fasta

```
>sp1
A-TGA
>sp2
AG-GC
>sp3
AG-AA
>sp4
AGTGA
>sp5
AGTGA
```

atpB.fasta

```
>sp1
ATCA
>sp2
ATCA
>sp3
ATCA
>sp4
ATCA
>sp5
CGCA
>sp6
ATCA
>sp7
ATCA
```

atpE.fasta

```
>sp6
GGCATTTT
>sp7
GGCATTTT
>sp8
GGCATTTT
```

2. Output -> all.fasta

```
>sp1
A-TGAATCA--------
>sp2
AG-GCATCA--------
>sp3
AG-AAATCA--------
>sp4
AGTGAATCA--------
>sp5
AGTGACGCA--------
>sp6
-----ATCAGGCATTTT
>sp7
-----ATCAGGCATTTT
>sp8
---------GGCATTTT
```

#### 7.5. Remove Gaps

***Functionality***

This tool can batchly remove gap sites from multiple alignment matrices.

***Features***

1. Gap sites in multiple alignment matrices or sub-alignments can be batchly removed.
2. Non-aligned sequence matrices cannot be allowed as input files.
3. Nucleotides, amino acids, upper case letters, lower case letters, gap (–), and special letters such as “N”, “n”, “X”, “x” (except 4 types of nucleotides, i.e., A, T, G, C, and 20 types of amino acids, i.e., A, C, D, E, F, G, H, I, K, L, M, N, P, Q, R, S, T, V, W, Y), are allowed in the alignment matrices.
4. For “N” or “n” in the nucleotide alignment matrices, it will not be removed. That is to say, “N” or “n” is not treated as gap site. For “X” or “x” in the amino acid alignment matrices, it will be removed. That is to say, “X” or “x” is treated as gap site.
5. The file “gapsites_atpA.txt” will be generated for each alignment matrix with gap sites, and it contains two information, i.e., “Gap position(s) in alignment matrix” and “Gap site(s) in alignment matrix (i.e. -)”. If the alignment matrix does not contain gap sites, this file will not be generated.

***Command-line Tool***

perl remove_gaps.pl –i –o

run:

perl remove_gaps.pl –i input –o output

parameter:

[-i -input] required: (default: input) input directory containing .fasta format alignment matrices.

[-o -output] required: (default: output) output directory.

***Example***

1. Input -> atpA.fasta, atpB.fasta, atpF.fasta

| | | | |
|---|---|---|---|
| atpA.fasta | 2023/4/14 17:21 | FASTA 文件 | 1 KB |
| atpB.fasta | 2022/12/12 16:18 | FASTA 文件 | 1 KB |
| atpF.fasta | 2023/4/14 17:08 | FASTA 文件 | 1 KB |

atpA.fasta

```
>sp1
atgtgtgaaaaagacaaagattctgagaaaatgtat-----gaagaagacaaagattct
>sp2
atgtgt-----------------------------------gaagaaaacaaagattct
>sp3
atgtgt-----------------------------------gaggaaaac---------
>sp4
atgtgt-----------------------------------gaagaagacaaagattct
>sp5
atgttt-----------------------------------gaggaaaacaaggattct
>sp6
atgtgt-----------------------------------gaagaagacaaagattct
```

atpB.fasta

```
>sp1
ATGCATGCATG-NGTGCTTCCAT-TGTGCATCCATGT
>sp2
TTGCTTCCATG--CTGCTTCCATGAGTGCTTCCATGT
>sp3
GTGCTTCCATG--ATGCATGCATGCGTGCT-CGATGT
>sp4
TTGCTTCCATG--GTGCTTCCATGTGTGCTTCCACGT
```

atpF.fasta

```
>sp1
MTLNL-X-VLTPNRSVWNSQVKAIILSTNSGQIGVLKDHAATATAVDIGVLRMLGLDDQW
>sp2
MTLNL---VLTPNRSVWNSQVKAIILSTNSGQIGVLKDHAATATAVDIGVLRMLGLDDQW
>sp3
MTLNL-------NRSVWNSQVKAIILSTNSGQIGVLKDHAATATAVDIGVLRMLGLDDQW
```

2. Output -> atpA.fasta, atpB.fasta, atpF.fasta, gapsites_atpA.txt, gapsites_atpB.txt, gapsites_atpF.txt

| | | | |
|---|---|---|---|
| atpA.fasta | 2023/4/14 17:21 | FASTA 文件 | 1 KB |
| atpB.fasta | 2023/4/14 17:21 | FASTA 文件 | 1 KB |
| atpF.fasta | 2023/4/14 17:21 | FASTA 文件 | 1 KB |
| gapsites_atpA.txt | 2023/4/14 17:21 | TXT 文件 | 1 KB |
| gapsites_atpB.txt | 2023/4/14 17:21 | TXT 文件 | 1 KB |
| gapsites_atpF.txt | 2023/4/14 17:21 | TXT 文件 | 1 KB |

atpA.fasta

```
>sp1
ATGTGTGAAAAAGACAAAGATTCTGAGAAAATGTATGAAGAAGACAAAGATTCT
>sp2
ATGTGT------------------------------GAAGAAAACAAAGATTCT
>sp3
ATGTGT------------------------------GAGGAAAAC---------
>sp4
ATGTGT------------------------------GAAGAAGACAAAGATTCT
>sp5
ATGTTT------------------------------GAGGAAAACAAGGATTCT
>sp6
ATGTGT------------------------------GAAGAAGACAAAGATTCT
```

atpB.fasta

```
>sp1
ATGCATGCATGNGTGCTTCCAT-TGTGCATCCATGT
>sp2
TTGCTTCCATG-CTGCTTCCATGAGTGCTTCCATGT
>sp3
GTGCTTCCATG-ATGCATGCATGCGTGCT-CGATGT
>sp4
TTGCTTCCATG-GTGCTTCCATGTGTGCTTCCACGT
```

atpF.fasta

```
>sp1
MTLNLVLTPNRSVWNSQVKAIILSTNSGQIGVLKDHAATATAVDIGVLRMLGLDDQW
>sp2
MTLNLVLTPNRSVWNSQVKAIILSTNSGQIGVLKDHAATATAVDIGVLRMLGLDDQW
>sp3
MTLNL----NRSVWNSQVKAIILSTNSGQIGVLKDHAATATAVDIGVLRMLGLDDQW

```

gapsites_atpA.txt

```
Gap position(s) in alignment matrix:
37 38 39 40 41

Gap site(s) in alignment matrix (i.e. -):
>sp1
-----
>sp2
-----
>sp3
-----
>sp4
-----
>sp5
-----
>sp6
-----

```

gapsites_atpB.txt

```
Gap position(s) in alignment matrix:
12

Gap site(s) in alignment matrix (i.e. -):
>sp1
-
>sp2
-
>sp3
-
>sp4
-

```

gapsites_atpF.txt

```
Gap position(s) in alignment matrix:
6 7 8

Gap site(s) in alignment matrix (i.e. -):
>sp1
-X-
>sp2
---
>sp3
---

```

##### 7.6. Generate PartitionFinder

***Functionality***

This tool will generate PartitionFinder format file based on fasta or phylip format multiple alignment matrices for further phylogenetic reconstruction.

***Features***

1. The fasta format (.fasta/.fas/.fa/.fsa/.phy) or phyip (.phylip/.phy) format alignment matrices are all allowed.
2. Non-aligned sequence matrices cannot be allowed as input files.
3. A "PartitionFinder.txt" file will be generated for further phylogenetic reconstruction.

***Command-line Tool***

perl generate_PartitionFinder.pl –i –o

run:

perl generate_PartitionFinder.pl –i input –o output

parameter:

[-i -input]  required: (default: input) input directory name containing fasta or phylip alignment matrices.

[-o -output]  required: (default: output) output file name.

***Example***

1. Input -> atpA.fasta, atpB.phy

| | | | |
|---|---|---|---|
| atpA.fasta | 2023/4/14 17:54 | FASTA 文件 | 1 KB |
| atpB.phy | 2023/4/14 17:53 | PHY 文件 | 1 KB |

atpA.fasta

```
>sp1
ATGTGTGAAAAAGACAAAGATTCTGAGAAAATGTAT---GAAGAAGACAAAGATTCTGAGGAAATG
>sp2
ATGTGT---------------------------------GAAGAAAACAAAGATTCTGAAAAAATG
>sp3
ATGTGT---------------------------------GAGGAAAAC------------------
>sp4
ATGTGT---------------------------------GAAGAAGACAAAGATTCTGAGAAAATG
>sp5
ATGTTT---------------------------------GAGGAAAACAAGGATTCTGAAAATATT
>sp6
ATGTGTGAAGAAAACAAAGATTCTGAGAAAATG---TGTGAAGAAGACAAAGATTCTGAGAAAATG
```

atpB.phy

```
 4 35
sp1         ATGCATGCATGGTGCTTCCAT-TGTGCATCCATGT
sp2         TTGCTTCCATGCTGCTTCCATGAGTGCTTCCATGT
sp3         GTGCTTCCATGATGCATGCATGCGTGCT-CGATGT
sp4         TTGCTTCCATGGTGCTTCCATGTGTGCTTCCACGT
```

2. Output -> PartitionFinder.txt

```
atpA = 1-66;
atpB = 67-101;
```

**8. Phylogeny**

***Functionality***

This tool can batchly perform maximum likelihood phylogenetic inference for each sequence matrix.

***Features***

1. Phylogenetic trees for each of alignment matrices can be batchly inferred by the frequently used phylogenetic analyses tool–RAxML. If you use it, please cite following paper.

Alexandros Stamatakis, RAxML version 8: a tool for phylogenetic analysis and post-analysis of large phylogenies, Bioinformatics, 2014, 30(9), 1312–1313.

2. The command of RAxML phylogenetic analyses will be set to this mode "raxmlHPC -f a -x 12345 -p 12345 -k -N 100 -m GTRGAMMA -s atpA.fasta -n atpA.result". In this command, the parameters "-N" and "-m" can be adjusted, with 100/200/500/1000 and GTRGAMMA/GTRGAMMAI/GTRCAT/GTRCATI, PROTGAMMAGTR/PROTCATGTR/, respectively.

3. If duplicated species are existed within the same sequence matrix file, RAxML phylogenetic inference will not be run, and the warning information like "Duplicated species name Arabidopsis_thaliana for the gene atpA!" will be outputed to the "warning.txt" file.

4. If the sequence matrix is satisfied with species number <= 50 and sequence length <= 10,000, RAxML phylogenetic inference will be normally run; if the sequence matrix is satisfied with species number > 50 or if the sequence matrix is satisfied with sequence length > 10,000, RAxML phylogenetic inference will not be run, and the warning information like "Species number larger than 50 for the gene atpA!" or "Sequence length longer than 10 kb for the gene atpA!" will be outputed to the "warning.txt" file.

***Command-line Tool***

perl batch_RAxML.pl –i –r –m –o

run:

perl batch_RAxML.pl -i input -r 100 -m GTRGAMMA -o output

parameter:

[-i -input] required: (default: input) input directory containing .fasta sequence matrices.

[-r -runs] required: (default: 100) number of runs, i.e., 100/200/500/1000.

[-m -model] required: (default: GTRGAMMA) evolutionary models, i.e., GTRGAMMA/GTRGAMMAI/GTRCAT/GTRCATI, PROTGAMMAGTR/ /PROTCATGTR/.

[-o -output] required: (default: output) output directory.

***Example***

1. Input -> atpA.fasta, atpB.fasta, ycf1.fasta

| | | | |
|---|---|---|---|
| atpA.fasta | 2023/4/14 21:46 | FASTA 文件 | 10 KB |
| atpB.fasta | 2022/12/14 10:23 | FASTA 文件 | 10 KB |
| ycf1.fasta | 2023/4/14 22:10 | FASTA 文件 | 73 KB |

2. Output -> RAxML_bestTree.atpA.result, RAxML_bipartitions.atpA.result, RAxML_bipartitionsBranchLabels.atpA.result, RAxML_bootstrap.atpA.result, RAxML_info.atpA.result

| | | | |
|---|---|---|---|
| RAxML_bestTree.atpA.result | 2023/4/14 22:11 | RESULT 文件 | 1 KB |
| RAxML_bestTree.atpB.result | 2023/4/14 22:11 | RESULT 文件 | 1 KB |
| RAxML_bipartitions.atpA.result | 2023/4/14 22:11 | RESULT 文件 | 1 KB |
| RAxML_bipartitions.atpB.result | 2023/4/14 22:11 | RESULT 文件 | 1 KB |
| RAxML_bipartitionsBranchLabels.atpA.result | 2023/4/14 22:11 | RESULT 文件 | 1 KB |
| RAxML_bipartitionsBranchLabels.atpB.result | 2023/4/14 22:11 | RESULT 文件 | 1 KB |
| RAxML_bootstrap.atpA.result | 2023/4/14 22:11 | RESULT 文件 | 34 KB |
| RAxML_bootstrap.atpB.result | 2023/4/14 22:11 | RESULT 文件 | 34 KB |
| RAxML_info.atpA.result | 2023/4/14 22:11 | RESULT 文件 | 15 KB |
| RAxML_info.atpB.result | 2023/4/14 22:11 | RESULT 文件 | 15 KB |
| warning.txt | 2023/4/14 22:11 | TXT 文件 | 1 KB |

3. Output -> Warning.txt

```
Sequence length longer than 10 kb for the gene ycf1!
```

**9. Barcoding**

***Functionality***

This tool can be used to batchly calculate and extract variable, invariable, and gap sites from multiple alignment matrices.

***Features***

1. The variable, invariable, and gap sites can be batchly calculated and extracted from multiple fasta format alignment matrices.

2. Number, percentage, position, and nucleotides of variable sites or parsimony informative sites for each alignment matrix can be calculated and extracted. The files like "atpA_variable_sites_allnucleotides.fasta" or "atpA _parsimony_informative_sites.fasta" will be generated for each alignment matrix, and it contains nucleotides of focused sites. The files like "atpA _variable_sites_allnucleotides.txt" and "atpA _parsimony_informative_sites.txt" will be simultaneously generated for each alignment matrix, and it contains number, percentage, position, and nucleotides of focused sites.

3. The alignment matrices can be sorted according to percentage of variable sites or parsimony informative sites, which can assist users to judge which matrices can be selected as potential markers. The files like "markers_by_parsimony_informative_sites.txt" or "markers_by_variable_sites.txt" will be generated for all alignment matrices, and they contain the quantitative sorting information.

4. There are 10 statistic patterns within this tool:

1 = variable sites all nucleotides;

2 = variable sites 2 nucleotides;

3 = variable sites 3 nucleotides;

4 = variable sites 4 nucleotides;

5 = parsimony informative sites;

6 = non-parsimony informative sites;

7 = invariable sites;

8 = gap sites;

9 = 1+5;

10 = 1+2+3+4+5+6+7+8.

5. All sites = Statistic pattern 1 + Statistic pattern 7 + Statistic pattern 8. That is to say, "all sites" = "1 = variable sites all nucleotides" + "7 = invariable sites" + "8 = gap sites".

Statistic pattern 1 = Statistic pattern 2 + Statistic pattern 3 + Statistic pattern 4. That is to say, "1 = variable sites all nucleotides" = "2 = variable sites 2 nucleotides" + "3 = variable sites 3 nucleotides" + "4 = variable sites 4 nucleotides".

Statistic pattern 1 = Statistic pattern 5 + Statistic pattern 6. That is to say, "1 = variable sites all nucleotides" = "5 = parsimony informative sites" + "6 = non-parsimony informative sites".

***Principles***

1. Variable sites are equal to polymorphic sites. Variable sites are defined as >= 2 types of nucleotides (>= 2 species), which is equal to above mentioned "1 = variable sites all nucleotides". There are three types of variable sites: variable sites with 2 types of nucleotides (>= 2 species), variable sites with 3 types of nucleotides (>= 3 species), and variable sites with 4 types of nucleotides (>= 4 species), which are equal to above mentioned "2 = variable sites 2 nucleotides",

“3 = variable sites 3 nucleotides”, and “4 = variable sites 4 nucleotides”, respectively.

Variable sites can also be divided into parsimony informative sites and non-parsimony informative sites. Parsimony informative sites are defined as all columns with >= 2 types of nucleotides and each nucleotide occurred >= 2 times, which is equal to above mentioned “5 = parsimony informative sites”. Non-parsimony informative sites are the parts of variable sites other than parsimony informative sites, which is equal to above mentioned “6 = non-parsimony informative sites”. For non-parsimony informative sites, if there are only 2 species in the matrix, Statistic pattern 6 = Statistic pattern 1 = Statistic pattern 2; if there are only 3 species in the matrix, Statistic pattern 6 = Statistic pattern 1 = Statistic pattern 2 + Statistic pattern 3; if there are >= 4 species in the matrix, Statistic pattern 6 = Statistic pattern 1 – Statistic pattern 5 = Statistic pattern 2 + Statistic pattern 3 + Statistic pattern 4 – Statistic pattern 5.

2. Invariable sites are equal to monomorphic sites or constant sites. Invariable sites are defined as all columns with identical nucleotides and without gaps, which is equal to above mentioned “7 = invariable sites”.

3. Gap sites are equal to sites with gaps or missing data. Gap sites are defined as sites with >= 1 gap (>= 2 species), which is equal to above mentioned “8 = gap sites”.

4. If there are only 2 species in the matrix, you can calculate Statistic pattern 1, Statistic pattern 2, Statistic pattern 6, Statistic pattern 7, and Statistic pattern 8; and Statistic pattern 1 = Statistic pattern 2 = Statistic pattern 6.

If there are only 3 species in the matrix, you can calculate Statistic pattern 1, Statistic pattern 2, Statistic pattern 3, Statistic pattern 6, Statistic pattern 7, and Statistic pattern 8; and Statistic pattern 1 = Statistic pattern 2 + Statistic pattern 3 = Statistic pattern 6.

If there are >= 4 species in the matrix, you can calculate Statistic pattern 1, Statistic pattern 2, Statistic pattern 3, Statistic pattern 4, Statistic pattern 5, Statistic pattern 6, Statistic pattern 7, and Statistic pattern 8; and Statistic pattern 1 = Statistic pattern 2 + Statistic pattern 3 + Statistic pattern 4 = Statistic pattern 5 + Statistic pattern 6.

***Command-line Tool***

perl barcoding.pl –i –p –o

run:

perl barcoding.pl –i input –p 9 –o output

parameter:

[-i -input] required: (default: input) input directory name containing fasta alignment matrices.

[-p -pattern] required: (default: 9) 1/2/3/4/5/6/7/8/9/10, statistic pattern.
1=variable sites all nucleotides, 2=variable sites 2 nucleotides,
3=variable sites 3 nucleotides, 4=variable sites 4 nucleotides,
5=parsimony informative sites, 6=non-parsimony informative sites,
7=invariable sites, 8=gap sites, 9=1+5, 10=1+2+3+4+5+6+7+8.

[-o -output] required: (default: output) output directory name.

***Example***

1. Input -> test1.fasta, test2.fasta, test3.fasta, test4.fasta

| | | | |
|---|---|---|---|
| test1.fasta | 2022/12/14 17:47 | FASTA 文件 | 1 KB |
| test2.fasta | 2022/12/15 12:36 | FASTA 文件 | 1 KB |
| test3.fasta | 2022/12/15 12:35 | FASTA 文件 | 1 KB |
| test4.fasta | 2022/12/15 17:07 | FASTA 文件 | 1 KB |

test1.fasta

```
>sp1
AT--
>sp2
AGC-

```

test2.fasta

```
>sp1
ATC--
>sp2
AGGC-
>sp3
AGAC-

```

test3.fasta

```
>sp1
ATCCT-------
>sp2
AGCGC---AGG-
>sp3
AGGAG-AAATC-
>sp4
AGGAAGAGATA-

```

test4.fasta

```
>sp1
ATCCT----?--
>sp2
AGCGC---AGG-
>sp3
AGGAG-AAATC-
>sp4
AGGAAGAGATA-

```

2. Output -> markers_by_parsimony_informative_sites.txt, markers_by_variable_sites.txt, test3_p5_parsimony_informative_sites.fasta, test3_p5_parsimony_informative_sites.txt, test3_p1_variable_sites_allnucleotides.fasta, test3_p1_variable_sites_allnucleotides.txt

| | | | |
|---|---|---|---|
| markers_by_parsimony_informative_sites.txt | 2023/4/18 0:34 | TXT 文件 | 1 KB |
| markers_by_variable_sites.txt | 2023/4/18 0:34 | TXT 文件 | 1 KB |
| test1_p1_variable_sites_allnucleotides.fasta | 2023/4/18 0:34 | FASTA 文件 | 1 KB |
| test1_p1_variable_sites_allnucleotides.txt | 2023/4/18 0:34 | TXT 文件 | 1 KB |
| test2_p1_variable_sites_allnucleotides.fasta | 2023/4/18 0:34 | FASTA 文件 | 1 KB |
| test2_p1_variable_sites_allnucleotides.txt | 2023/4/18 0:34 | TXT 文件 | 1 KB |
| test3_p1_variable_sites_allnucleotides.fasta | 2023/4/18 0:34 | FASTA 文件 | 1 KB |
| test3_p1_variable_sites_allnucleotides.txt | 2023/4/18 0:34 | TXT 文件 | 1 KB |
| test3_p5_parsimony_informative_sites.fasta | 2023/4/18 0:34 | FASTA 文件 | 1 KB |
| test3_p5_parsimony_informative_sites.txt | 2023/4/18 0:34 | TXT 文件 | 1 KB |
| test4_p1_variable_sites_allnucleotides.fasta | 2023/4/18 0:34 | FASTA 文件 | 1 KB |
| test4_p1_variable_sites_allnucleotides.txt | 2023/4/18 0:34 | TXT 文件 | 1 KB |
| test4_p5_parsimony_informative_sites.fasta | 2023/4/18 0:34 | FASTA 文件 | 1 KB |
| test4_p5_parsimony_informative_sites.txt | 2023/4/18 0:34 | TXT 文件 | 1 KB |

markers_by_parsimony_informative_sites.txt

```
Sequence_matrix percentage_of_parsimony_informative_sites        number_of_parsimony_informative_sites    total_length_of_matrix
test4   0.0833  1       12
test3   0.0833  1       12
```

markers_by_variable_sites.txt

```
Sequence_matrix percentage_of_variable_sites     number_of_variable_sites        total_length_of_matrix
test2   0.4000  2       5
test3   0.3333  4       12
test4   0.3333  4       12
test1   0.2500  1       4
```

test3_p5_parsimony_informative_sites.txt

```
Total sequence length of alignment matrix test3:
12

Number of parsimony informative site(s) in alignment matrix test3:
1

Percentage of variable/polymorphic site(s) with all nucleotides in alignment matrix test3:
0.0833

Position of parsimony informative site(s) in alignment matrix test3:
3

Parsimony informative site(s) in alignment matrix test3:
>sp1
C
>sp2
C
>sp3
G
>sp4
G
```

test3_p5_parsimony_informative_sites.fasta

```
>sp1
C
>sp2
C
>sp3
G
>sp4
G
```

test3_p1_variable_sites_allnucleotides.txt

```
Total sequence length of alignment matrix test3:
12

Number of variable/polymorphic site(s) with all nucleotides in alignment matrix test3:
4

Percentage of variable/polymorphic site(s) with all nucleotides in alignment matrix test3:
0.3333

Position of variable/polymorphic site(s) with all nucleotides in alignment matrix test3:
2 3 4 5

Variable/polymorphic site(s) with all nucleotides in alignment matrix test3:
>sp1
TCCT
>sp2
GCGC
>sp3
GGAG
>sp4
GGAA
```

test3_p1_variable_sites_allnucleotides.fasta

```
>sp1
TCCT
>sp2
GCGC
>sp3
GGAG
>sp4
GGAA
```

#### 10. Plastomics

At present, this tool kit contains two tools that can complete comparative analysis of plastomes, such as "10.1. Generate mVISTA, and 10.2. Gene Homology".

##### 10.1. Generate mVISTA

***Functionality***

This tool can batchly generate input files for mVISTA alignment based on GenBank flatfiles.

***Features***

1. Input files for mVISTA alignment can be batchly generated from GenBank format flatfiles (.gb/.gbk/.gbf).

2. Only essential information from GenBank flatfiles are retained in input files for mVISTA alignment, such as gene names (*psbA*), coordinates of genes (378     1439), forward (>) or reverse (<) coding strands, and exon of protein-coding genes or utr of tRNA and rRNA genes.

***Command-line Tool***

perl mVISTA.pl –i –o

run:
perl mVISTA.pl –i input –o output

parameter:
[-i -input]     required: (default: input) input directory name containing GenBank flatfiles.
[-o -output]     required: (default: output) output directory name.

***Example***

1. Input -> Rosa_roxburghii.gb

```
LOCUS       Rosa_roxburghii  156749 bp    DNA     circular PLN 02-DEC-2022
FEATURES             Location/Qualifiers
     source          1..156749
                     /organism="Rosa_roxburghii"
                     /organelle="plastid:chloroplast"
                     /mol_type="genomic DNA"
                     /note="Annotation Method :: PGA-Plastid Genome Annotator"
     repeat_region   85853..111905
                     /note="inverted repeat B"
                     /rpt_type="inverted"
     repeat_region   complement(130697..156749)
                     /note="inverted repeat A"
                     /rpt_type="inverted"
     gene            58781..60244
                     /gene="accD"
     CDS             58781..60244
                     /gene="accD"
                     /codon_start=1
                     /transl_table=11
                     /product="acetyl-CoA carboxylase carboxyltransferase beta subunit"
     gene            complement(10635..12158)
                     /gene="atpA"
     CDS             complement(10635..12158)
                     /gene="atpA"
                     /codon_start=1
                     /transl_table=11
                     /product="ATP synthase CF1 alpha subunit"
```

2. Output -> Rosa_roxburghii_mVISTA.txt

```
<	4	77	trnH-GUG
4	77	utr
<	378	1439	psbA
378	1439	exon
<	1704	4278	trnK-UUU
1704	1738	utr
4242	4278	utr
<	1998	3509	matK
1998	3509	exon
<	5247	6381	rps16
5247	5476	exon
6342	6381	exon
<	7213	7284	trnQ-UUG
7213	7284	utr
>	7704	7889	psbK
7704	7889	exon
>	8126	8236	psbI
8126	8236	exon
<	8379	8466	trnS-GCU
8379	8466	utr
```

**10.2. Gene Homology**

***Functionality***

This tool can be used to generate circular view of two plastomes with homologous genes connected and non-connected genes marked. First, the homologous genes in the target plastome and their corresponding genes in the reference plastome can be connected by this tool. So, collinear state of these two plastomes can be judged by the connection status of homologous genes. Second, the genes that are not connected in reference or target plastome can be marked by this tool. Therefore, annotation completeness of plastomes can be assessed by comparing gene homology and marking non-connected genes.

***Features***

1. The circular view of two plastomes, with connected homologous genes, is generated based on the Circos tool. If you use it, we suggest you to cite following paper.

Krzywinski M, Schein J, Birol I, et al. Circos: an information aesthetic for comparative genomics. Genome Research, 2009, 19(9): 1639–1645.2. Both SVG and PNG format figures are exported based on the Circos tool.

2. Both SVG and PNG format figures are exported. All intermediate documents for generating the circular plastome map are also provided for downloading.

3. The color type of gene blocks follows the OGDRAW tool, and the color block illustration figure (color_block_illustration.pdf) can be individually downloaded.

4. The connection status of homologous genes depends largely on the reference plastome selected by the user.

5. The genes that are not connected in reference or target plastome will be marked with red triangle symbol, which indicates that these genes need to be carefully checked.

***Recommended Suggestions***

1. The standard GenBank format flatfiles should contain information of correct quadripartite structure. That is to say, from 0 bp to 156749 bp (for the whole length of Rosa roxburghii plastome), the quadripartite structure must be LSC-IRb-SSC-IRa.

2. Whether the repeat region (i.e., IRb and IRa) of plastome are annotated in the GenBank flatfiles or not does not affect the display of quadripartite structure (i.e., LSC, SSC, IRb, and IRa).

***Command-line Tool***

perl gene_homology.pl -r -t –o

perl gene_homology.pl -r -t -m -a -p -r -c -b –o

run:

perl gene_homology.pl -r reference/Amborella_trichopoda.gb -t target/Rosa_roxburghii.gb -o output

perl gene_homology.pl -r reference/Amborella_trichopoda.gb -t target/Rosa_roxburghii.gb -m 1000 -a -90 -p 1to1 -r no -c 0.50 -b 0.25 -o output

perl gene_homology.pl -r reference/Amborella_trichopoda.gb -t target/Rosa_roxburghii.gb -m 1000 -a 270 -p 1to1 -r n -c 0.50 -b 0.25 -o output parameter:

[-r -reference] required: (default: reference.gb) input reference file of GenBank-formatted file.

[-t -target] required: (default: target.gb) input target file of GenBank-formatted file.

[-m -minir] required: (default: 1000) minimum allowed length for the longest inverted-repeat (IR).

[-a -angle] required: (default: -90) any rotation angle of the homologous gene map, -90=270.

[-p -pattern] required: (default: 1to1) patterns (1to1/1tomore) for linking homologous genes.

[-r -ribbon] required: (default: no) ribbon (yes/y) or line (no/n) for linking homologous genes.

[-c -crest] required: (default: 0.50) crest peak distance of linking lines or ribbons, and the alternative values are [0.01/0.25/0.50/0.75/1.00].

[-b -bezier] required: (default: 0.25) bezier curve degree of linking lines or ribbons, and the alternative values are [0.01/0.25/0.50/0.75/1.00].

[-o -output] required: (default: output) output directory name.

***Parameter Interpretation***

1. Only two GenBank files are allowed to upload. That is to say, only one reference plastome and one target plastome can be compared at a time.

2. The filenames of these two GenBank files can be sorted in forward or reverse ASCII/alphabetic order.

3. The longest inverted-repeat (IR) is generated by blast search the complete plastome against itself, and the minimum allowed length (with default value of 1000 bp) can be changed by users.

4. The rotation angle of the homologous gene map can be freely adjusted according to users.

5. If one IR copy or two IR copies are lost from any of reference or target plastomes, we suggest you to use "1tomore" pattern, which can not only link syntenic orthologous genes but also link paralogous genes. Therefore, you can find the position of genes occurred two or more times.

If no IR copy/copies are lost from both reference and target plastomes, we suggest you to use "1to1" pattern, which can link orthologous genes. In this case, most genes are linked with 1to1 pattern. However, several genes that not occurred in IR of target plastome but occurred in IR of reference plastome will be linked with 1tomore pattern, which can assist users to find the genes experienced IR expansion or contraction.

6. Ribbon or line for linking homologous genes can be freely selected by the users.

7. Both crest and bezier parameters can determine the bending degree of the connecting lines.

***Principles***

Micro-synteny comparision of each gene in target and reference plastomes is shown in Figure 1.

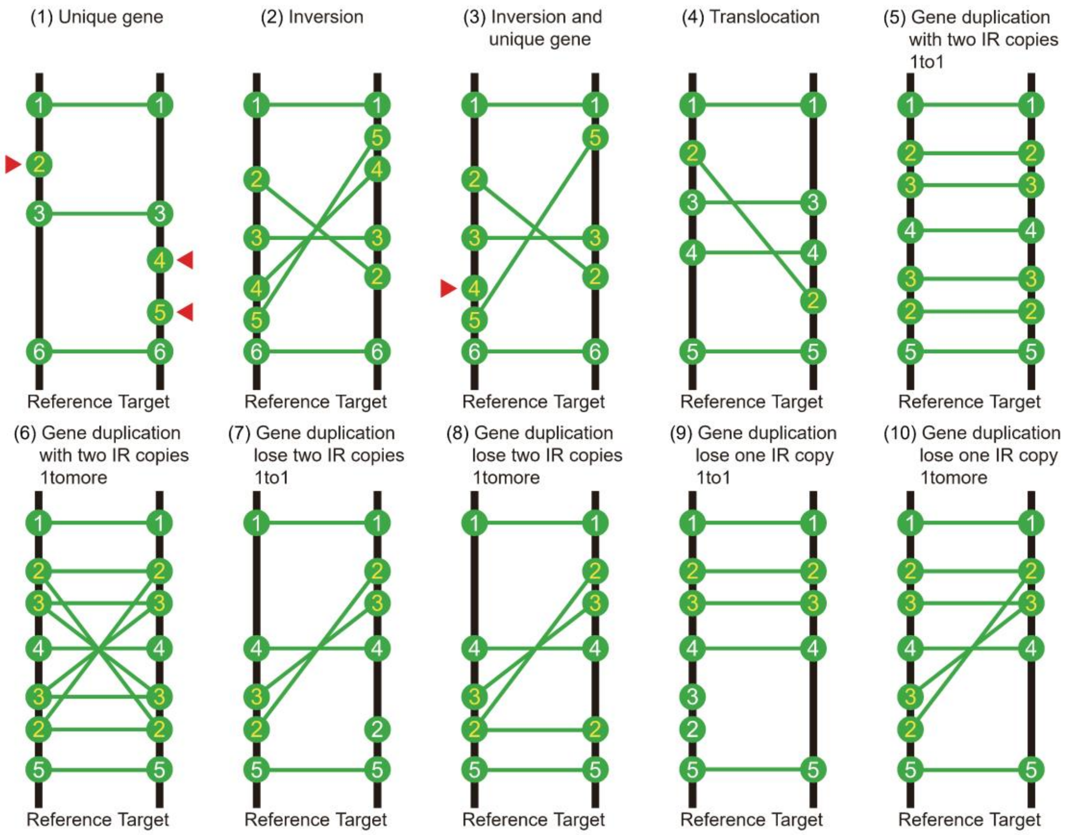


Figure 1. Micro-synteny comparision of each gene in target and reference plastomes.

(1) Unique gene. Unique genes in target or reference plastome will be marked with red triangle symbol, waiting to be carefully checked, for evaluating the annotation quality of plastomes.

(2) Inversion. Inversion, common factor in shaping genome structures, can be shown by connecting the genes in the reciprocally inverted segments.

(3) Inversion and unique gene. Not all genes in the inverted segments are shared between target and reference, so the unique genes in the inverted segments are also be marked with red triangle symbol.

(4) Translocation. Translocation is common factor in reconstructing genome structures.

(5) Gene duplication, with two IR copies, 1to1. For gene duplication, if no IR copies are lost from both plastomes, and if the 1to1 pattern is selected, the tool will only connect shared genes with the same order.

(6) Gene duplication, with two IR copies, 1tomore. For gene duplication, if no IR copies are lost from both plastomes, and if the 1tomore pattern is selected, the tool will connect shared genes with both the same and opposite order.

(7) Gene duplication, lose two IR copies, 1to1. For gene duplication, if two IR copies are lost from both plastomes, and if the 1to1 pattern is selected, the tool will only connect shared genes with same or opposite order.

(8) Gene duplication, lose two IR copies, 1tomore. For gene duplication, if two IR copies are lost from both plastomes, and if the 1tomore pattern is selected, the tool will connect shared genes with both same and opposite order.

(9) Gene duplication, lose one IR copy, 1to1. For gene duplication, if only one IR copy is lost from target or reference, and if the 1to1 pattern is selected, the tool will only connect shared genes with same or opposite order.

(10) Gene duplication, lose one IR copy, 1tomore. For gene duplication, if only one IR copy is lost from target or reference, and if the 1tomore pattern is selected, the tool will connect shared genes with both the same and opposite order.

***Color block illustration***

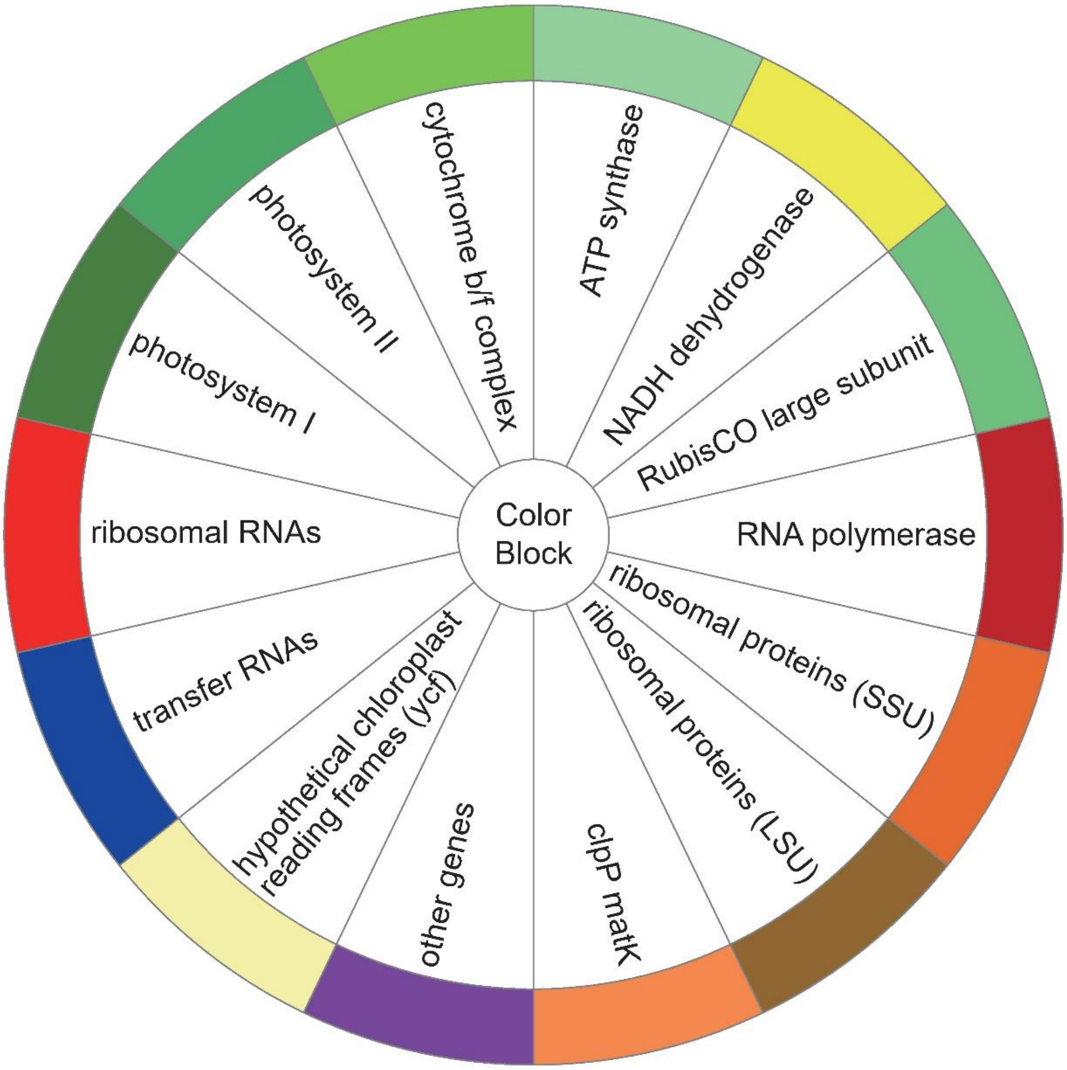

***Example***

Input:

Amborella_trichopoda.gb

Rosa_roxburghii.gb

Output:

Amborella_trichopoda_innergenes.label.txt

Amborella_trichopoda_innergenes.position.txt

Amborella_trichopoda_outergenes.label.txt

Amborella_trichopoda_outergenes.position.txt

Amborella_trichopoda_quadripartite.IRb.IRa.txt

genes.link.txt

genes.unique.reference.txt

genes.unique.target.txt

homology.conf

homology.png

homology.svg

ideogram.conf

karyotype.txt

Rosa_roxburghii_innergenes.label.txt

Rosa_roxburghii_innergenes.position.txt

Rosa_roxburghii_outergenes.label.txt

Rosa_roxburghii_outergenes.position.txt

Rosa_roxburghii_quadripartite.IRb.IRa.txt

species_length.txt

ticks.conf

1. Input -> Amborella_trichopoda.gb

```
LOCUS       Amborella_trichopoda      162686 bp    DNA     circular UNA 08-JUN-2015
DEFINITION  Amborella trichopoda chloroplast genomic DNA, complete sequence.
ACCESSION   AJ506156
VERSION     AJ506156.2  GI:34481608
KEYWORDS    complete genome.
SOURCE      chloroplast Amborella trichopoda
  ORGANISM  Amborella trichopoda
            Eukaryota; Viridiplantae; Streptophyta; Embryophyta; Tracheophyta;
            Spermatophyta; Magnoliophyta; basal Magnoliophyta; Amborellales;
            Amborellaceae; Amborella.
FEATURES          Location/Qualifiers
     source          1..162686
                    /organism="Amborella trichopoda"
                    /mol_type="genomic DNA"
     repeat_region    90951..117611
                    /note="inverted repeat region B; IRB repeat region"
                    /rpt_type="inverted"
     rRNA          complement(139284..142097)
                    /gene="rrn23"
                    /product="23S ribosomal RNA"
     gene           complement(139284..142097)
                    /gene="rrn23"
     tRNA          join(complement(4472..4508), complement(1840..1874))
                    /gene="trnK-UUU"
                    /product="tRNA-Lys"
     gene           complement(1840..4508)
                    /gene="trnK-UUU"
     CDS           join(complement(16186..16330), complement(14506..14915))
                    /gene="atpF"
                    /codon_start=1
                    /transl_table=11
                    /product="ATPase I subunit"
                    /translation="MKNVTDSFVSLGHWPSAGSFGFNTDIFATNPINLSVVLGVLIFF
                    GKGVLSDLLDNRKQRILSTIRNSEELRGGAIEQLEKARARLRKVEIEADEFRVNGYSE
                    IEREKSNLINAAYENLERLENYKNESIHFEQQRAMNQVRQRVFQQALQGALETLNSYL
                    NSELHLRTISANIGMLGTMKNITD"
     gene           complement(14506..16330)
                    /gene="atpF"
```

2. Input -> Rosa_roxburghii.gb

```
LOCUS       Rosa_roxburghii  156749 bp    DNA     circular PLN 02-DEC-2022
FEATURES             Location/Qualifiers
     source          1..156749
                     /organism="Rosa_roxburghii"
                     /organelle="plastid:chloroplast"
                     /mol_type="genomic DNA"
                     /note="Annotation Method :: PGA-Plastid Genome Annotator"
     repeat_region   85853..111905
                     /note="inverted repeat B"
                     /rpt_type="inverted"
     repeat_region   complement(130697..156749)
                     /note="inverted repeat A"
                     /rpt_type="inverted"
     gene            58781..60244
                     /gene="accD"
     CDS             58781..60244
                     /gene="accD"
                     /codon_start=1
                     /transl_table=11
                     /product="acetyl-CoA carboxylase carboxyltransferase beta subunit"
     gene            complement(10635..12158)
                     /gene="atpA"
     CDS             complement(10635..12158)
                     /gene="atpA"
                     /codon_start=1
                     /transl_table=11
                     /product="ATP synthase CF1 alpha subunit"
```

3. Output -> Amborella_trichopoda_innergenes.label.txt

```
Amborella_trichopoda	1	64	trnH-GUG
Amborella_trichopoda	428	1480	psbA
Amborella_trichopoda	1840	4508	trnK-UUU
Amborella_trichopoda	2197	3702	matK
Amborella_trichopoda	6144	7204	rps16
Amborella_trichopoda	8596	8667	trnQ-UUG
Amborella_trichopoda	9852	9939	trnS-GCU
Amborella_trichopoda	12897	14420	atpA
Amborella_trichopoda	14506	16330	atpF
Amborella_trichopoda	16686	16931	atpH
Amborella_trichopoda	17989	18735	atpI
Amborella_trichopoda	19009	19719	rps2
Amborella_trichopoda	19978	24087	rpoC2
Amborella_trichopoda	24263	27026	rpoC1
Amborella_trichopoda	27053	30271	rpoB
Amborella_trichopoda	33691	33798	psbM
Amborella_trichopoda	34739	34812	trnD-GUC
Amborella_trichopoda	35359	35442	trnY-GUA
Amborella_trichopoda	35513	35585	trnE-UUC
Amborella_trichopoda	40465	40557	trnS-UGA
```

4. Output -> Amborella_trichopoda_innergenes.position.txt

```
Amborella_trichopoda	1	64	fill_color=chr14
Amborella_trichopoda	428	1480	fill_color=dgreen
Amborella_trichopoda	1840	4508	fill_color=chr14
Amborella_trichopoda	2197	3702	fill_color=orange
Amborella_trichopoda	6144	7204	fill_color=dorange
Amborella_trichopoda	8596	8667	fill_color=chr14
Amborella_trichopoda	9852	9939	fill_color=chr14
Amborella_trichopoda	12897	14420	fill_color=lgreen
Amborella_trichopoda	14506	16330	fill_color=lgreen
Amborella_trichopoda	16686	16931	fill_color=lgreen
Amborella_trichopoda	17989	18735	fill_color=lgreen
Amborella_trichopoda	19009	19719	fill_color=dorange
Amborella_trichopoda	19978	24087	fill_color=chr4
Amborella_trichopoda	24263	27026	fill_color=chr4
Amborella_trichopoda	27053	30271	fill_color=chr4
Amborella_trichopoda	33691	33798	fill_color=dgreen
Amborella_trichopoda	34739	34812	fill_color=chr14
Amborella_trichopoda	35359	35442	fill_color=chr14
Amborella_trichopoda	35513	35585	fill_color=chr14
Amborella_trichopoda	40465	40557	fill_color=chr14
```

5. Output -> Amborella_trichopoda_outergenes.label.txt

```
Amborella_trichopoda	9027	9212	psbK
Amborella_trichopoda	9615	9725	psbI
Amborella_trichopoda	11684	12534	trnG-UCC
Amborella_trichopoda	12694	12765	trnR-UCU
Amborella_trichopoda	31492	31562	trnC-GCA
Amborella_trichopoda	32567	32656	petN
Amborella_trichopoda	36404	36475	trnT-GGU
Amborella_trichopoda	37818	38879	psbD
Amborella_trichopoda	38827	40248	psbC
Amborella_trichopoda	40893	41081	psbZ
Amborella_trichopoda	41480	41550	trnG-GCC
Amborella_trichopoda	50207	50293	trnS-GGA
Amborella_trichopoda	52167	52725	trnL-UAA
Amborella_trichopoda	53101	53173	trnF-GAA
Amborella_trichopoda	58030	58102	trnM-CAU
Amborella_trichopoda	60985	62412	rbcL
Amborella_trichopoda	63125	64723	accD
Amborella_trichopoda	65349	65459	psaI
Amborella_trichopoda	65914	66621	ycf4
Amborella_trichopoda	67486	68175	cemA
```

6. Output -> Amborella_trichopoda_outergenes.position.txt

```
Amborella_trichopoda    9027    9212    fill_color=dgreen
Amborella_trichopoda    9615    9725    fill_color=dgreen
Amborella_trichopoda    11684   12534   fill_color=chr14
Amborella_trichopoda    12694   12765   fill_color=chr14
Amborella_trichopoda    31492   31562   fill_color=chr14
Amborella_trichopoda    32567   32656   fill_color=chrun
Amborella_trichopoda    36404   36475   fill_color=chr14
Amborella_trichopoda    37818   38879   fill_color=dgreen
Amborella_trichopoda    38827   40248   fill_color=dgreen
Amborella_trichopoda    40893   41081   fill_color=dgreen
Amborella_trichopoda    41480   41550   fill_color=chr14
Amborella_trichopoda    50207   50293   fill_color=chr14
Amborella_trichopoda    52167   52725   fill_color=chr14
Amborella_trichopoda    53101   53173   fill_color=chr14
Amborella_trichopoda    58030   58102   fill_color=chr14
Amborella_trichopoda    60985   62412   fill_color=green
Amborella_trichopoda    63125   64723   fill_color=chr19
Amborella_trichopoda    65349   65459   fill_color=chr13
Amborella_trichopoda    65914   66621   fill_color=vlyellow
Amborella_trichopoda    67486   68175   fill_color=chr19
```

7. Output -> Amborella_trichopoda_quadripartite.IRb.IRa.txt

```
Amborella_trichopoda    90951   117611  fill_color=black,z=0
Amborella_trichopoda    136026  162686  fill_color=black,z=0
```

8. Output -> genes.link.txt

```
Amborella_trichopoda    1       64      Rosa_roxburghii 4       77
Amborella_trichopoda    428     1480    Rosa_roxburghii 378     1439
Amborella_trichopoda    1840    4508    Rosa_roxburghii 1704    4278
Amborella_trichopoda    2197    3702    Rosa_roxburghii 1998    3509
Amborella_trichopoda    6144    7204    Rosa_roxburghii 5247    6381
Amborella_trichopoda    8596    8667    Rosa_roxburghii 7213    7284
Amborella_trichopoda    9027    9212    Rosa_roxburghii 7704    7889
Amborella_trichopoda    9615    9725    Rosa_roxburghii 8126    8236
Amborella_trichopoda    9852    9939    Rosa_roxburghii 8379    8466
Amborella_trichopoda    11684   12534   Rosa_roxburghii 9069    9834
Amborella_trichopoda    12694   12765   Rosa_roxburghii 10027   10098
Amborella_trichopoda    12897   14420   Rosa_roxburghii 10635   12158
Amborella_trichopoda    14506   16330   Rosa_roxburghii 12213   12767
Amborella_trichopoda    16686   16931   Rosa_roxburghii 13249   13494
Amborella_trichopoda    17989   18735   Rosa_roxburghii 14427   15170
Amborella_trichopoda    19009   19719   Rosa_roxburghii 15396   16106
Amborella_trichopoda    19978   24087   Rosa_roxburghii 16351   20505
Amborella_trichopoda    24263   27026   Rosa_roxburghii 20683   23514
Amborella_trichopoda    27053   30271   Rosa_roxburghii 23525   26737
Amborella_trichopoda    31492   31562   Rosa_roxburghii 27932   28002
```

9. Output -> genes.unique.reference.txt

10. Output -> genes.unique.target.txt

11. Output -> homology.conf

```
<<include etc/colors_fonts_patterns.conf>>
<<include etc/housekeeping.conf>>

<<include ideogram.conf>>
<<include ticks.conf>>

<image>
<<include etc/image.conf>>
angle_orientation* = counterclockwise
#background* = transparent
#radius* = 1500p
angle_offset* = -90
</image>

karyotype    = homology\output/karyotype.txt
chromosomes_units           = 1000
chromosomes_display_default = yes
chromosomes_reverse = /Rosa_roxburghii/
```

12. Output -> homology.png

13. Output -> homology.svg

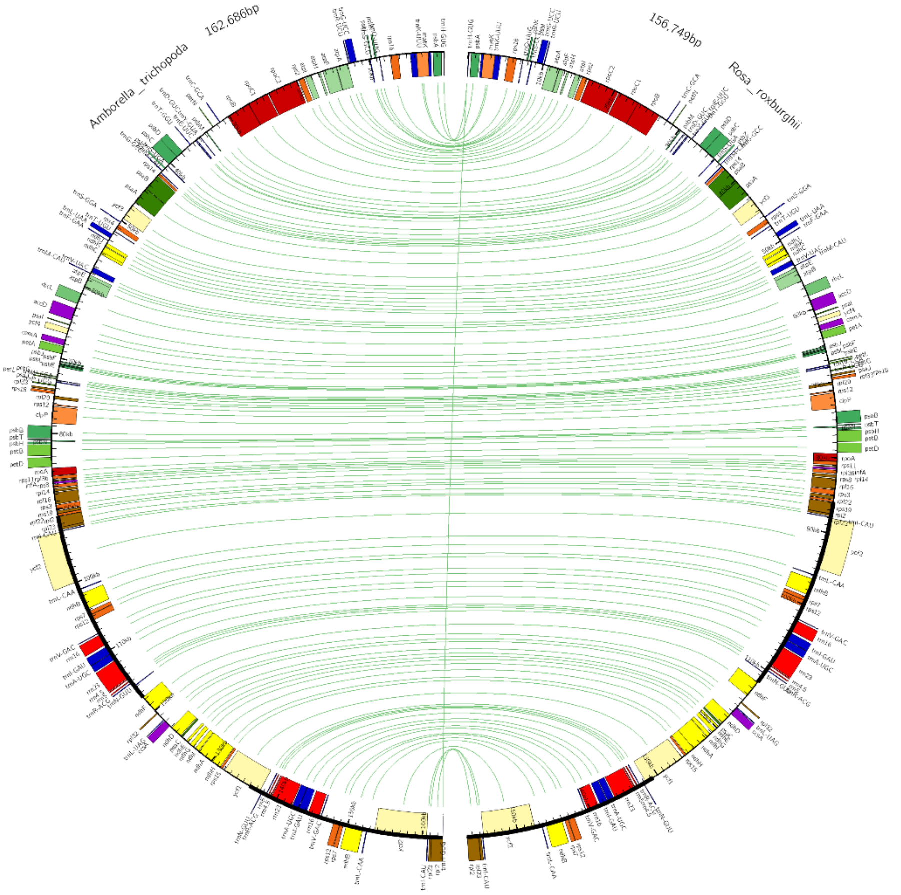


14. Output -> ideogram.conf

```
<ideogram>

<spacing>
default = 0.01r
break   = 0
</spacing>


radius           = 0.8r
thickness        = 2p
fill             = yes

fill_color       = gpos100
stroke_thickness = 2
stroke_color     = black

</ideogram>
```

15. Output -> karyotype.txt

```
chr - Amborella_trichopoda Amborella_trichopoda 0 162686 black
chr - Rosa_roxburghii Rosa_roxburghii 0 156749 black
```

16. Output -> Rosa_roxburghii_innergenes.label.txt

```
Rosa_roxburghii 4       77      trnH-GUG
Rosa_roxburghii 378     1439    psbA
Rosa_roxburghii 1704    4278    trnK-UUU
Rosa_roxburghii 1998    3509    matK
Rosa_roxburghii 5247    6381    rps16
Rosa_roxburghii 7213    7284    trnQ-UUG
Rosa_roxburghii 8379    8466    trnS-GCU
Rosa_roxburghii 10635   12158   atpA
Rosa_roxburghii 12213   12767   atpF
Rosa_roxburghii 13249   13494   atpH
Rosa_roxburghii 14427   15170   atpI
Rosa_roxburghii 15396   16106   rps2
Rosa_roxburghii 16351   20505   rpoC2
Rosa_roxburghii 20683   23514   rpoC1
Rosa_roxburghii 23525   26737   rpoB
Rosa_roxburghii 30165   30269   psbM
Rosa_roxburghii 30793   30866   trnD-GUC
Rosa_roxburghii 31287   31370   trnY-GUA
Rosa_roxburghii 31430   31502   trnE-UUC
Rosa_roxburghii 36070   36162   trnS-UGA
```

17. Output -> Rosa_roxburghii_innergenes.position.txt

```
Rosa_roxburghii 4       77      fill_color=chr14
Rosa_roxburghii 378     1439    fill_color=dgreen
Rosa_roxburghii 1704    4278    fill_color=chr14
Rosa_roxburghii 1998    3509    fill_color=orange
Rosa_roxburghii 5247    6381    fill_color=dorange
Rosa_roxburghii 7213    7284    fill_color=chr14
Rosa_roxburghii 8379    8466    fill_color=chr14
Rosa_roxburghii 10635   12158   fill_color=lgreen
Rosa_roxburghii 12213   12767   fill_color=lgreen
Rosa_roxburghii 13249   13494   fill_color=lgreen
Rosa_roxburghii 14427   15170   fill_color=lgreen
Rosa_roxburghii 15396   16106   fill_color=dorange
Rosa_roxburghii 16351   20505   fill_color=chr4
Rosa_roxburghii 20683   23514   fill_color=chr4
Rosa_roxburghii 23525   26737   fill_color=chr4
Rosa_roxburghii 30165   30269   fill_color=dgreen
Rosa_roxburghii 30793   30866   fill_color=chr14
Rosa_roxburghii 31287   31370   fill_color=chr14
Rosa_roxburghii 31430   31502   fill_color=chr14
Rosa_roxburghii 36070   36162   fill_color=chr14
```

18. Output -> Rosa_roxburghii_outergenes.label.txt

```
Rosa_roxburghii 7704    7889    psbK
Rosa_roxburghii 8126    8236    psbI
Rosa_roxburghii 9069    9834    trnG-UCC
Rosa_roxburghii 10027   10098   trnR-UCU
Rosa_roxburghii 27932   28002   trnC-GCA
Rosa_roxburghii 28813   28902   petN
Rosa_roxburghii 32018   32089   trnT-GGU
Rosa_roxburghii 33399   34460   psbD
Rosa_roxburghii 34408   35829   psbC
Rosa_roxburghii 36549   36737   psbZ
Rosa_roxburghii 37110   37180   trnG-GCC
Rosa_roxburghii 46148   46234   trnS-GGA
Rosa_roxburghii 48741   49374   trnL-UAA
Rosa_roxburghii 49780   49852   trnF-GAA
Rosa_roxburghii 53733   53805   trnM-CAU
Rosa_roxburghii 56695   58122   rbcL
Rosa_roxburghii 58781   60244   accD
Rosa_roxburghii 60727   60840   psaI
Rosa_roxburghii 61261   61815   ycf4
Rosa_roxburghii 62269   62958   cemA
```

19. Output -> Rosa_roxburghii_outergenes.position.txt

```
Rosa_roxburghii 7704    7889    fill_color=dgreen
Rosa_roxburghii 8126    8236    fill_color=dgreen
Rosa_roxburghii 9069    9834    fill_color=chr14
Rosa_roxburghii 10027   10098   fill_color=chr14
Rosa_roxburghii 27932   28002   fill_color=chr14
Rosa_roxburghii 28813   28902   fill_color=chrun
Rosa_roxburghii 32018   32089   fill_color=chr14
Rosa_roxburghii 33399   34460   fill_color=dgreen
Rosa_roxburghii 34408   35829   fill_color=dgreen
Rosa_roxburghii 36549   36737   fill_color=dgreen
Rosa_roxburghii 37110   37180   fill_color=chr14
Rosa_roxburghii 46148   46234   fill_color=chr14
Rosa_roxburghii 48741   49374   fill_color=chr14
Rosa_roxburghii 49780   49852   fill_color=chr14
Rosa_roxburghii 53733   53805   fill_color=chr14
Rosa_roxburghii 56695   58122   fill_color=green
Rosa_roxburghii 58781   60244   fill_color=chr19
Rosa_roxburghii 60727   60840   fill_color=chr13
Rosa_roxburghii 61261   61815   fill_color=vlyellow
Rosa_roxburghii 62269   62958   fill_color=chr19
```

20. Output -> Rosa_roxburghii_quadripartite.IRb.IRa.txt

```
Rosa_roxburghii 85853   111905  fill_color=black,z=0
Rosa_roxburghii 130697  156749  fill_color=black,z=0
```

21. Output -> species_length.txt

```
Amborella_trichopoda    18487   29579   162,686bp
Amborella_trichopoda    28049   45190   Amborella_trichopoda
Rosa_roxburghii 17812   28499   156,749bp
Rosa_roxburghii 27025   43541   Rosa_roxburghii
```

22. Output -> ticks.conf

```
show_ticks          = yes
show_tick_labels    = yes
show_grid           = yes

<ticks>
skip_first_label     = no
skip_last_label      = no
radius               = dims(ideogram,radius_outer)
tick_separation      = 2p
min_label_distance_to_edge = 10p
label_separation = 5p
label_offset     = 5p
multiplier       = 1e-3
color            = black

<tick>
spacing        = 1u
size           = 15p
thickness      = 2p
orientation    = in
```
